# SPROUTS_DB: an implemented database of contaminants for extracellular vesicle proteomics studies

**DOI:** 10.1101/2025.05.20.655024

**Authors:** Maria Gaetana Giovanna Pittalà, Loredana Leggio, Greta Paternò, Elena Giusto, Laura Civiero, Vincenzo Cunsolo, Silvia Vivarelli, Antonella Di Francesco, Emanuele Alpi, Nunzio Iraci, Rosaria Saletti

## Abstract

**Background:** Current proteomics techniques allow rapid identification and quantification of proteins within any given biological source. In particular, nanoUHPLC/High-Resolution nanoESI-MS/MS enables the characterization of proteins in complex biological samples due to its high sensitivity, accuracy, and scalability. However, LC-MS/MS proteomics might still be susceptible to laboratory and sample-associated contaminants, which can significantly compromise the quality and reliability of data. Therefore, an accurate identification and annotation of such contaminants is crucial for the development of robust proteomics databases and spectral-libraries related search engines. This approach is of special interest in the field of secretome and extracellular vesicles (EVs), membrane-enclosed nanostructures that contain a variety of proteins crucial for cell-to-cell communication and translational applications.

**Results:** When working in *ex vivo/in vitro* settings, proteins from fetal bovine serum (FBS), commonly employed in standard cell culture media, may interfere with the proteome analysis. To address this issue, we conceived and designed SPROUTS_DB, **S**erum **P**rotein **R**epository **O**f **U**nwanted **T**arget(ed) **S**equences **D**ata**B**ase, a dedicated resource to catalog serum-derived contaminants. Starting from media supplemented with EV-depleted FBS, we simulated cell growth conditions - in the absence of cells - followed by ultracentrifugation. LC-MS/MS analysis of these samples resulted in the identification of a novel set of 1,288 contaminant proteins, which has been deposited in the ProteomeXchange repository (identifier PXD044137).

SPROUTS_DB contains primarily soluble proteins, mainly related to the Gene Ontology categories Extracellular Region and Extracellular Space, in line with the nature of the starting sample. In contrast, only a small fraction of the contaminants is classified as membrane-associated proteins, supporting the limited vesicle contamination in the complete medium, due to the use of EV-depleted FBS.

Of note, we demonstrated that SPROUTS_DB outperforms existing contaminants’ databases, ensuring that only peptide spectra relevant to the examined sample are retained and identified as true positive data.

**Conclusions:** Considering that even proteins from phylogenetically distant organisms share extensive stretches of sequences, SPROUTS_DB is designed to discern contaminants from real sample proteins of interest, minimizing false positive identifications.

To the best of our knowledge, SPROUTS_DB is the most updated database of contaminants useful for proteomics investigations of cellular secretomes and EV-containing samples.

## Background

Deciphering the proteome of cells – and their secretome – in different conditions is an important step to understand the biological processes occurring inside the cells in a specific moment. Mass spectrometry (MS) has greatly expanded its applicability to the proteomics field, thereby revolutionizing the analysis of proteins and other biomolecules thanks to the development of “soft” ionization techniques, such us electrospray ionization (ESI) and matrix-assisted laser desorption/ionization (MALDI) [1]. Although current proteomics approaches allow to obtain accurate results in terms of number of proteins, post-translational modifications etc., there are yet some issues that need to be overcome [2]. In LC-MS/MS-based proteomics, the selection of an appropriate reference database is critical for enhancing true positive identifications while minimizing both false positives and false negatives, thereby directly impacting the accuracy and reliability of the final results [3]. Furthermore, all the main software for proteomics analysis allow users to indicate a supplementary list of proteins – referred to as contaminant database – which includes common laboratory contaminants such as human keratins or standards used for quantification. These proteins, often introduced during sample handling and processing, can lead to false positive identifications if not properly accounted for.

When working with *in vitro* models, one of the main problems is often related to the cell culture composition, where the presence of fetal bovine serum (FBS) may interfere with data interpretation. Several databases of contaminants have been developed to help researchers to deal with this issue [4]. While these databases serve as valuable tools for identifying proteins of genuine interest (true positive), they require further refinement to effectively discriminate against serum-derived contaminants commonly introduced through cell culture media.

A relevant area where contaminant databases may play a key role in discriminating proteins of interest from co-isolated contaminants is related to the study of extracellular vesicles (EVs) [5]. EVs are membranous nanoparticles produced by almost all cells under physiological and pathological states [6]. Based on their biogenesis, it is possible to distinguish two main classes of EVs: (i) exosomes, deriving from the endosomal compartments and (ii) shedding vesicles, directly released from the plasma membrane. Based on their size, EVs are classified as small EVs (< 200 nm, e.g., exosomes and small shedding vesicles), and medium/large EVs (> 200 nm, e.g., bigger microvesicles and oncosomes) [5,7]. However, the biogenesis of EVs is still under investigation, with novel emerging classes of EVs being discovered, whose origin and function(s) remain to be further elucidated [8]. In line with this effort, research strives to recognize specific markers for each sub-population of EVs, for example via proteomics [9]. About their functional potential, EVs are considered an important tool for cell-to-cell communication, carrying several classes of biomolecules such us lipids, DNA, RNAs and proteins, some of which are selectively sorted from the secreting cells [6]. Of note, these cargoes are protected from the action of nucleases and proteases thanks to the presence of a lipid-bilayer. When EVs are released in the extracellular milieu, they can interact with target cells, both in close proximity and travelling within biological fluid, to reach distant sites. Notably, EVs from different donor cells are able to release different cargoes, which can also change in response to alterations in the microenvironment. Thus, EVs can exert different functions depending on the “status” of the donor cell [10]. On their translational potential, EVs purified from almost all body fluids are emerging as source of novel biomarkers for several diseases [11]. Also, EVs are exploited as new therapeutics in nanomedicine, either in their native form, or opportunely engineered to obtain innovative drug delivery systems [11–14].

We recently demonstrated that astrocytes (AS) from the ventral midbrain (VMB) and the striatum (STR) - the two main brain regions involved in Parkinson’s disease (PD) - release a population of small-EVs in a region-specific manner [15,16]. In particular, only EVs from VMB-AS are able to fully recover the mitochondrial activity of injured target neurons, thus suggesting that EVs from distinct brain regions can exert different functions, with neuroprotective implications for PD [15]. These and other *in vitro* EV functional studies call for a deeper understanding of the cargoes shuttled within specific vesicle samples, while discerning from experimental contaminants. In this context, LC-MS/MS proteomics are crucial for EV characterization, function discovery and quality control. Several approaches have been developed to obtain a high yield of vesicles, including ultracentrifugation (UC), size-exclusion chromatography (SEC) and ultrafiltration [17,18]. To date, UC is still one of the most widely used methodologies for EV purification [4–10,19,20]. In *ex vivo*/*in vitro* research, EV-depleted FBS is often used to reduce the possible interference due to bovine serum-derived vesicles. However, it is not possible to exclude the presence of residual FBS-EV-derived proteins or other soluble bovine proteins in the complete media that may co-precipitate with the EVs of interest during EV isolation [19]. This contamination, intrinsic to the experimental procedure, can obscure true biological signals and lead to misleading interpretations.

In this work, we introduced SPROUTS_DB i.e., **S**erum **P**rotein **R**epository **O**f **U**nwanted **T**arget(ed) **S**equences **D**ata**B**ase, a novel integrated collection of contaminants, that will help addressing these issues. We carefully described all the passages that have led to SPROUTS_DB curation, from sample preparation (complete medium without cells), to the nanoUHPLC/High Resolution nanoESI-MS/MS analysis, in data-dependent acquisition (DDA) mode. Next, via Gene Ontology (GO) Enrichment Analysis, we identified the classes of proteins significantly enriched or depleted in our samples after UC, paying particular attention to the eventual contribution of residual bovine EV-derived proteins. We found that most of medium proteins were soluble components of the extracellular *milieu*, in line with the presence of FBS. Crucially, only a few of them were located in cellular membranes, supporting the limited vesicle contamination in the complete medium, due to the use of EV-depleted FBS.

Overall, SPROUTS_DB will contribute to the correct identification of proteins, while minimizing false positive attribution in complex biological samples, including – but not limited to – EV-containing specimens.

## Results

### SPROUTS_DB consists of over 1,200 contaminant proteins combining SwissProt, TrEMBL and Contaminants Database of the Max Planck Institute

To evaluate the presence of bovine-derived proteins (soluble and EV-related), 50 ml of complete DMEM medium containing 10% EV-depleted FBS were incubated for 24 h at 37 °C, 5% CO_2_, mimicking a typical cell culture setting **(Figure 1A)**. After the UC, ∼20 ± 0.5 µg of protein extracts were obtained in each biological replicate. Next, the lysates were treated with dithiothreitol (DTT), iodoacetamide (IAA) and trypsin in order to perform the subsequent analysis in MS, using a proteomic shotgun approach. MS data have been subjected to bioinformatic analysis and, in a single database search, a protein was considered identified if it fulfilled both of the following requirements: (i) a minimum of two peptides with a score above the peptide filtering threshold matches; (ii) at least a unique peptide in the list of the matched peptides. Also, to generate the final list of proteins, only those identified at least in two out of three LC-MS/MS technical replicates and at least in two out of three biological replicates were considered. The whole process used to identify complete medium proteins and to construct the SPROUTS_DB is described in **Figure 1B-C**. Of note, MS data were very reproducible, as shown by the intensity values of the Total Ion Current (TIC) chromatograms of the technical replicates (**Figure S1**).

**Figure 1.**
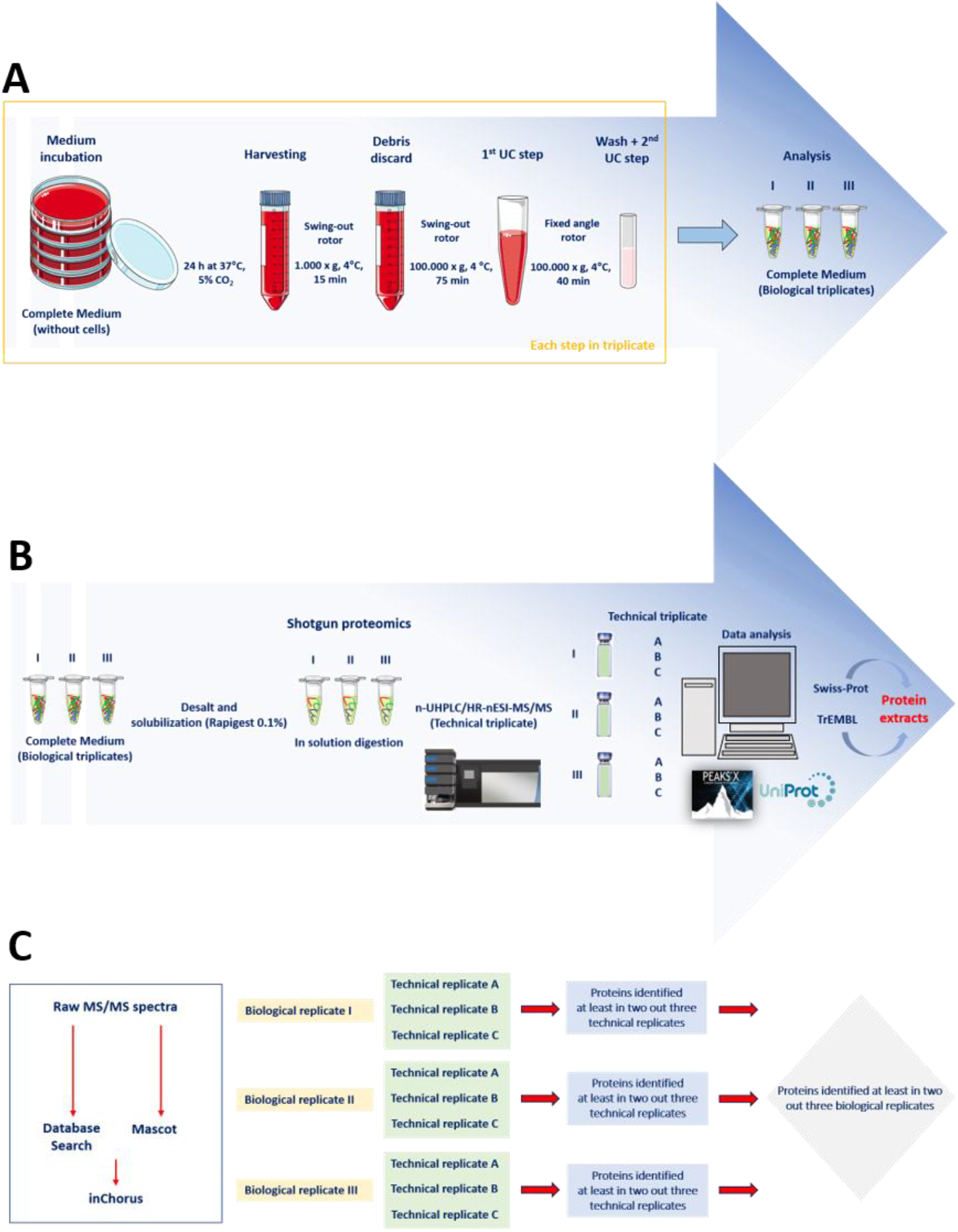
Description of the process leading to complete medium protein analysis. A) Sample preparation via UC. B) Shotgun proteomic approach and MS analysis. C) Database search and protein identification.

The list of medium proteins identified in our sample was obtained by merging the results of the search in the SwissProt section of UniProt (containing reviewed, manually annotated entries), with those from the TrEMBL section of UniProt (containing unreviewed, automatically annotated entries) (**Table S1**). Following the criteria detailed in the Experimental Section, a total of 1,073 proteins were identified. Among these, 536 matched entries in the SwissProt section and 537 in the TrEMBL section of UniProt. Next, to further corroborate the general validity of our database, we performed a similar LC-MS/MS analysis with medium supplemented with an alternative EV-depleted FBS. Interestingly, we found ≈90% of overlap between the 2 complete media preparations, with only 35 additional proteins specific for the new analysis, which were added to our list (underlined in **Table S1**).

Finally, to create a comprehensive contaminant database that also accounts for proteins inadvertently introduced during sample handling and processing (e.g., keratins etc.), we integrated the MaxQuant Contaminants Database from the Max Planck Institute (MPI) of Biochemistry (Martinsried, Germany) into our list of proteins.

The MaxQuant Contaminants Database consists of 245 proteins belonging to various organisms including *Bos taurus* and *Homo sapiens*. The list also contains common laboratory proteins used in enzymatic reactions (such as trypsin) and standards used in quantification (bovine serum albumin). IDs and amino acid sequences of the MaxQuant Contaminants list have been updated using UniProt and other available online resources as REFSEQ (https://www.ncbi.nlm.nih.gov/refseq), H-INV (http://www.h-invitational.jp/), and ENSEMBL (https://www.ensembl.org) databases. To avoid redundancies, we compared our list of proteins found in complete medium with the 245 proteins of this Contaminants Database. We found 65 common proteins between the two groups (highlighted in bold in **Table S1**).

The remaining 180 proteins of the MaxQuant Contaminants Database **(Table S2)** were added to the 1,108 proteins (1,073 + 35) identified in our sample, to obtain a final list of 1,288 (1,108 + 180) proteins **(Figure 2)**, that was used to create a unique contaminant database (See SPROUTS_DB.fasta in the Supplementary Information). Data are also available via ProteomeXchange with identifier PXD044137.

**Figure 2.**
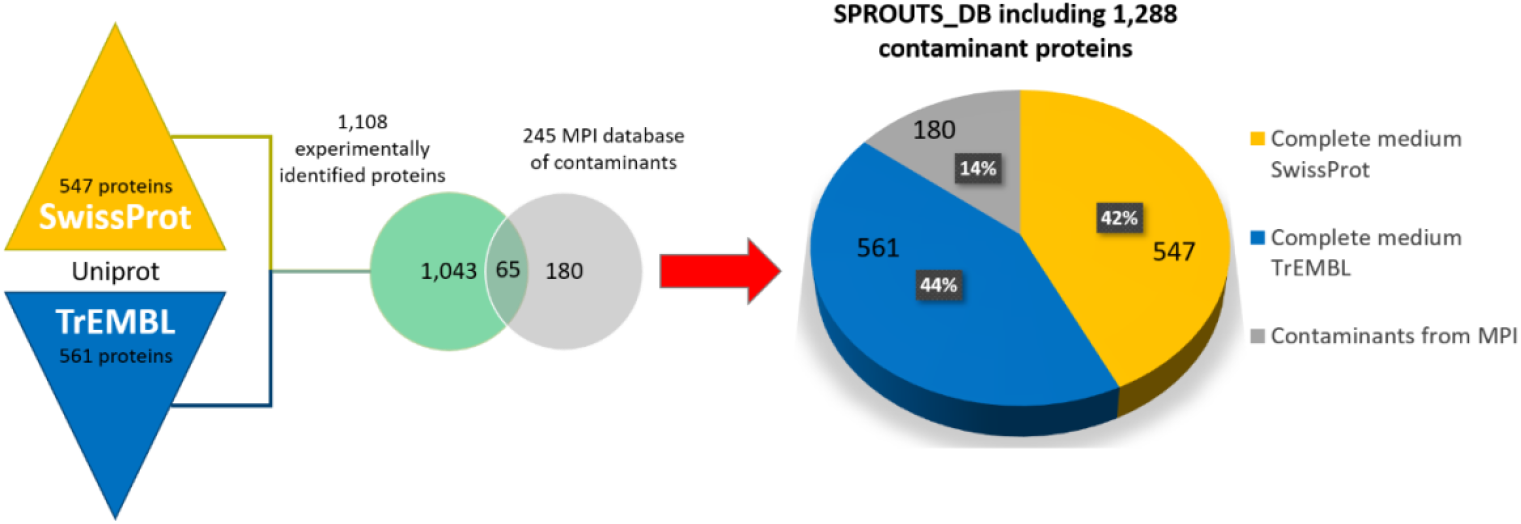
Process leading to SPROUTS_DB definition: from the protein identification in SwissProt and TrEMBL sections of UniProt to the comparison with MPI contaminant database. The exclusive proteins of MPI contaminant database (i.e., 180 proteins) were merged with the 1,108 proteins experimentally identified in the complete medium, to finally obtain SPROUTS_DB, consisting of 1,288 contaminant proteins. The pie chart reveals the contribution each database gave to the SPROUTS_DB composition.

### SPROUTS_DB outperforms existing contaminants’ databases

The entire set of peptides identified by the *Bos taurus* database search (5,294 peptides) was used to query the open source web application Unipept [21]. **Figure 3** shows the tree-graph results of Unipept investigation. The analysis revealed that 84.8% of total peptides are shared by all *Eukariot*a **(Figure 3A)**. The remainder of the peptides are ubiquitous and generically assigned to “organism” (shown in grey in **Figure 3A**). Among the *Eukariota* specific peptides, 61.1% belong to *Mammalia* **(Figure 3B)** and 55.5% of them are common to all *Ruminantia* **(Figure 3C)**. Finally, only 14.4% of these peptides are specific for the *Bos* genus **(Figure 3D)**. The percentage of peptides is calculated considering 100% as the total number of peptides of the previous node. These results highlight how certain protein sequences (e.g., structural proteins, active sites of various enzymes etc.) have been conserved during evolution and are still shared today by phylogenetically distant organisms. Hence, the need to find a method suitable for a better discrimination between proteins specifically belonging to one species rather than another. In other words, the choice of reference databases for contaminants plays a fundamental role in proteomics studies and affects the search results.

**Figure 3.**
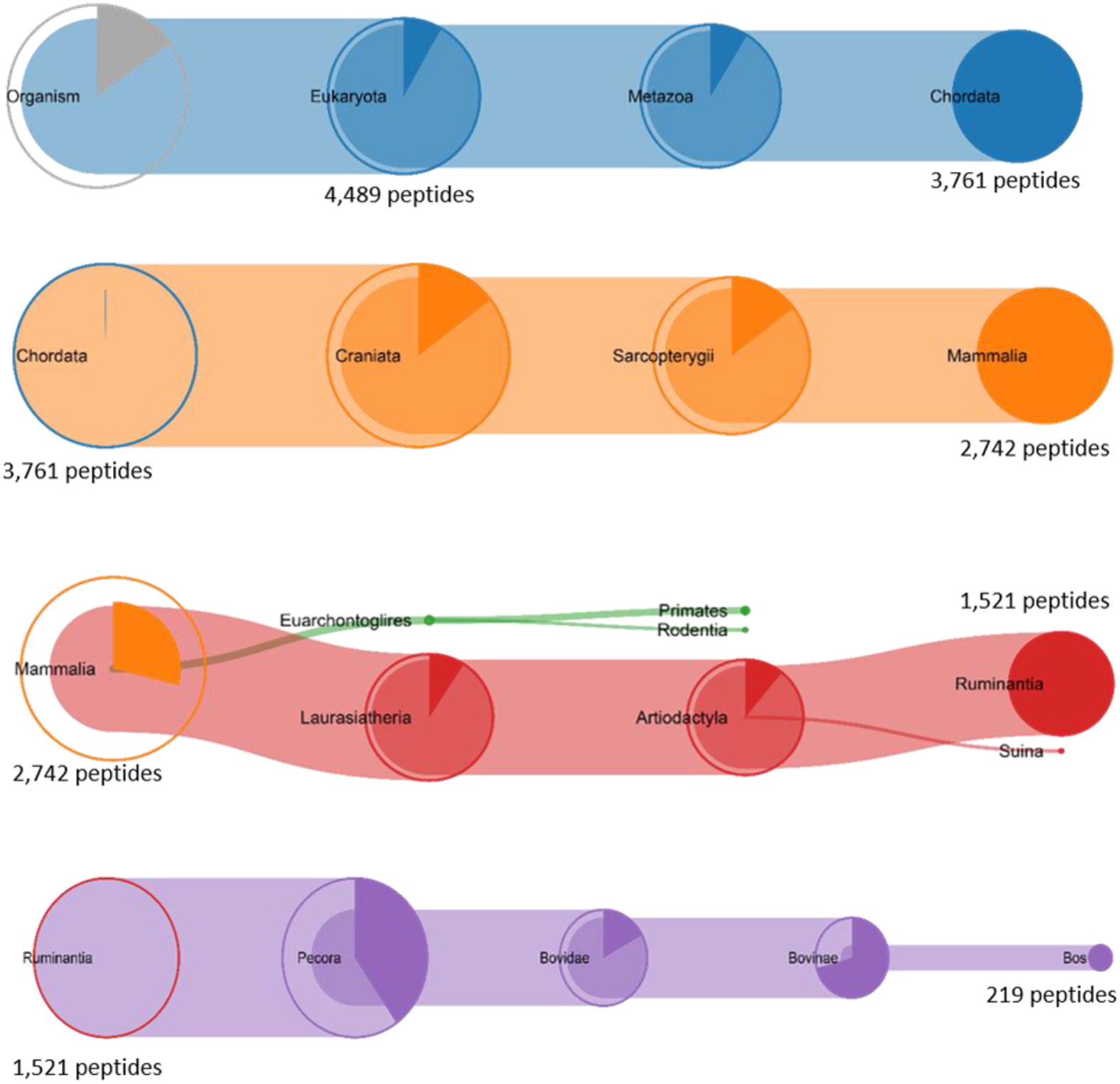
Tree-graph of Unipept application. The analysis of 5,294 culture medium peptides revealing that more than 80% of them are shared between all *Eukaryota*. Instead, only 219 peptides are specific for *Bos*.

To demonstrate the advancements of our database over other studies, we applied SPROUTS_DB to the MS/MS data by Shin et al. (deposited to the PRIDE repository with the data set identifier PXD015143) [22]. In this work, the authors introduced a collection of abundant FBS proteins, called Common Repository of FBS Proteins (cRFP), to be added to the reference database, to reduce false identifications in the MS raw data search of cell secretomes. Employing their raw data files, we compared the results between searches obtained against the Human DB (104,583 entries), the Human DB plus cRFP (104,583 + 199 entries) and the Human DB plus SPROUTS_DB (104,583 + 1,288 entries). The results of these comparisons are detailed in the “Supplementary Information” section and summarized in **Table 1**. Using the SPROUTS_DB together with the reference database (Search 3, **Table 1**) allows for the identification of a higher number of the protein contaminants (267 proteins vs. 56 identified by cRFP), and at the same time a lower number of true human proteins (1,347 proteins vs. 1,369 identified by cRFP).

**Table 1.**
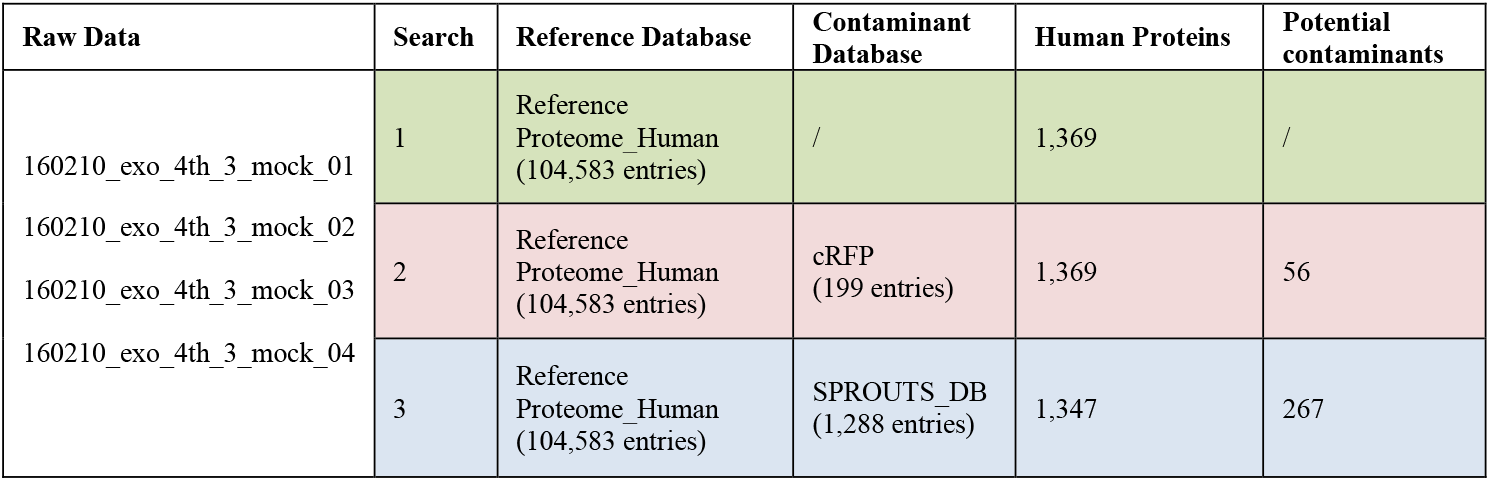
Comparison of results obtained using Human Database (Search 1), Human Database with cRFP (Search 2) and Human Database with SPROUTS_DB (Search 3).

In light of these analyses, we can conclude that SPROUTS_DB shows a greater performance in discriminating bovine contaminants, thus contributing to the correct identification of proteins in the sample of interest.

### Contaminant proteins are enriched in GO terms related to extracellular milieu

Once defined SPROUTS_DB as a unique contaminant database, we performed a relative quantitative analysis of the proteins specifically found in the complete medium after UC. The relative abundance of each protein identified in SwissProt through PEAKS X-Pro software was calculated using the average area in the technical and biological replicates **(Table S3)**. The list of the top 100 most abundant proteins is reported in **Table 2**.

**Table 2.** List of the Top 100 proteins quantified by PEAKS X-Pro software. For each protein is reported: accession number as in the UniProt database, protein name, gene code, score, number of the characterized peptides, number of unique peptides and average area.

|  | Accession number | Protein name | Gene code | Score (-10lgP) | Peptides | Unique peptides | Average area |
| --- | --- | --- | --- | --- | --- | --- | --- |
| 1 | Q7SIH1 | Alpha-2-macroglobulin | A2M | 666.87 | 247 | 247 | 1.59E+10 |
| 2 | P02769 | Albumin | ALB | 547.39 | 96 | 89 | 5.53E+09 |
| 3 | P12763 | Alpha-2-HS-glycoprotein | AHSG | 520.17 | 49 | 49 | 1.56E+09 |
| 4 | Q2UVX4 | Complement C3 | C3 | 590.1 | 116 | 116 | 1.27E+09 |
| 5 | P34955 | Alpha-1-antiproteinase | SERPINA1 | 399.32 | 30 | 30 | 1.24E+09 |
| 6 | P01966 | Hemoglobin subunit alpha | HBA | 310.25 | 10 | 10 | 9.70E+08 |
| 7 | P00735 | Prothrombin | F2 | 481.91 | 50 | 50 | 6.58E+08 |
| 8 | P02081 | Hemoglobin fetal subunit beta | N/A | 395.25 | 25 | 21 | 4.67E+08 |
| 9 | P41361 | Antithrombin-III | SERPINC1 | 419.64 | 28 | 28 | 3.72E+08 |
| 10 | Q29443 | Serotransferrin | TF | 480.42 | 49 | 46 | 2.54E+08 |
| 11 | P17697 | Clusterin | CLU | 345.87 | 18 | 18 | 1.87E+08 |
| 12 | Q95121 | Pigment epithelium-derived factor | SERPINF1 | 382 | 20 | 20 | 1.42E+08 |
| 13 | Q3MHL4 | Adenosylhomocysteinase | AHCY | 363.79 | 23 | 23 | 1.04E+08 |
| 14 | Q58D62 | Fetuin-B | FETUB | 320.17 | 14 | 14 | 1.02E+08 |
| 15 | Q05443 | Lumican | LUM | 283.6 | 14 | 14 | 9.17E+07 |
| 16 | P01017 | Angiotensinogen | AGT | 411.61 | 27 | 7 | 8.96E+07 |
| 17 | Q2KJ83 | Carboxypeptidase N catalytic chain | CPN1 | 340.47 | 13 | 13 | 8.39E+07 |
| 18 | P10096 | Glyceraldehyde-3-phosphate dehydrogenase | GAPDH | 365.54 | 15 | 13 | 8.31E+07 |
| 19 | Q3T052 | Inter-alpha-trypsin inhibitor heavy chain H4 | ITIH4 | 423.89 | 31 | 31 | 7.46E+07 |
| 20 | Q28107 | Coagulation factor V | F5 | 442.83 | 50 | 50 | 7.09E+07 |
| 21 | Q3SZ57 | Alpha-fetoprotein | AFP | 404.84 | 29 | 28 | 6.91E+07 |
| 22 | O46375 | Transthyretin | TTR | 323.88 | 9 | 9 | 6.85E+07 |
| 23 | Q3Y5Z3 | Adiponectin | ADIPOQ | 234.24 | 6 | 6 | 6.48E+07 |
| 24 | P06868 | Plasminogen | PLG | 402.08 | 38 | 38 | 6.33E+07 |
| 25 | P56652 | Inter-alpha-trypsin inhibitor heavy chain H3 | ITIH3 | 362.22 | 23 | 23 | 6.32E+07 |
| 26 | P15497 | Apolipoprotein A-I | APOA1 | 308.48 | 18 | 18 | 5.10E+07 |
| 27 | Q9XSJ4 | Alpha-enolase | ENO1 | 379.4 | 23 | 17 | 4.78E+07 |
| 28 | P81187 | Complement factor B | CFB | 393.22 | 27 | 27 | 4.24E+07 |
| 29 | Q0VCU1 | Cytoplasmic aconitate hydratase | ACO1 | 412.13 | 36 | 36 | 4.03E+07 |
| 30 | Q03247 | Apolipoprotein E | APOE | 297.9 | 11 | 11 | 3.87E+07 |
| 31 | P28800 | Alpha-2-antiplasmin | SERPINF2 | 308.14 | 12 | 12 | 3.70E+07 |
| 32 | Q3ZCJ2 | Aldo-keto reductase family 1 member A1 | AKR1A1 | 375.77 | 23 | 22 | 3.63E+07 |
| 33 | Q3T0P6 | Phosphoglycerate kinase 1 | PGK1 | 364.33 | 19 | 19 | 3.61E+07 |
| 34 | Q9N2I2 | Plasma serine protease inhibitor | SERPINA5 | 330.85 | 13 | 13 | 3.25E+07 |
| 35 | Q3SX14 | Gelsolin | GSN | 374.98 | 23 | 23 | 2.91E+07 |
| 36 | P00978 | Protein AMBP | AMBP | 312.06 | 11 | 11 | 2.83E+07 |
| 37 | Q32LP0 | Fermitin family homolog 3 | FERMT3 | 393.59 | 22 | 22 | 2.75E+07 |
| 38 | Q2KJF1 | Alpha-1B-glycoprotein | A1BG | 374.41 | 17 | 17 | 2.67E+07 |
| 39 | P02584 | Profilin-1 | PFN1 | 246.79 | 7 | 7 | 2.46E+07 |
| 40 | P62935 | Peptidyl-prolyl cis-trans isomerase A | PPIA | 273.82 | 10 | 10 | 2.42E+07 |
| 41 | P52898 | Dihydrodiol dehydrogenase 3 | N/A | 362.45 | 16 | 16 | 2.40E+07 |
| 42 | P01267 | Thyroglobulin | TG | 477.61 | 61 | 61 | 2.40E+07 |
| 43 | P68103 | Elongation factor 1-alpha 1 | EEF1A1 | 303.67 | 11 | 11 | 2.32E+07 |
| 44 | O46415 | Ferritin light chain | FTL | 250.13 | 6 | 6 | 2.28E+07 |
| 45 | A6H768 | Galactokinase | GALK1 | 356.39 | 15 | 15 | 2.23E+07 |
| 46 | Q2KIG3 | Carboxypeptidase B2 | CPB2 | 317.74 | 15 | 15 | 2.21E+07 |
| 47 | Q92176 | Coronin-1A | CORO1A | 304.36 | 16 | 16 | 2.17E+07 |
| 48 | Q3SZR3 | Alpha-1-acid glycoprotein | ORM1 | 283.21 | 8 | 8 | 2.10E+07 |
| 49 | Q2KJH9 | 4-trimethylaminobutyraldehyde dehydrogenase | ALDH9A1 | 277.31 | 13 | 13 | 2.04E+07 |
| 50 | Q2KJH4 | WD repeat-containing protein 1 | WDR1 | 339.39 | 17 | 17 | 1.94E+07 |
| 51 | Q27975 | Heat shock 70 kDa protein 1A | HSPA1A | 370.73 | 19 | 11 | 1.91E+07 |
| 52 | Q27965 | Heat shock 70 kDa protein 1B | HSPA1B | 370.73 | 19 | 11 | 1.91E+07 |
| 53 | P52556 | Flavin reductase (NADPH) | BLVRB | 325.49 | 9 | 9 | 1.80E+07 |
| 54 | Q0VCK0 | Bifunctional purine biosynthesis protein ATIC | ATIC | 322.53 | 20 | 20 | 1.78E+07 |
| 55 | P14568 | Argininosuccinate synthase | ASS1 | 289.04 | 12 | 12 | 1.77E+07 |
| 56 | Q3SZV7 | Hemopexin | HPX | 328.93 | 12 | 12 | 1.75E+07 |
| 57 | P19120 | Heat shock cognate 71 kDa protein | HSPA8 | 380.74 | 21 | 14 | 1.73E+07 |
| 58 | A3KMV5 | Ubiquitin-like modifier-activating enzyme 1 | UBA1 | 389.49 | 27 | 27 | 1.65E+07 |
| 59 | Q3SYV4 | Adenylyl cyclase-associated protein 1 | CAP1 | 333.75 | 14 | 14 | 1.54E+07 |
| 60 | P81948 | Tubulin alpha-4A chain | TUBA4A | 325.66 | 15 | 5 | 1.46E+07 |
| 61 | Q3T054 | GTP-binding nuclear protein Ran | RAN | 195.51 | 8 | 8 | 1.45E+07 |
| 62 | Q3SYU2 | Elongation factor 2 | EEF2 | 315.15 | 23 | 23 | 1.44E+07 |
| 63 | Q3MHN2 | Complement component C9 | C9 | 295.57 | 15 | 15 | 1.39E+07 |
| 64 | P61223 | Ras-related protein Rap-1b | RAP1B | 238.08 | 11 | 11 | 1.30E+07 |
| 65 | P07589 | Fibronectin | FN1 | 417.14 | 38 | 38 | 1.28E+07 |
| 66 | P28801 | Glutathione S-transferase P | GSTP1 | 245.61 | 6 | 6 | 1.24E+07 |
| 67 | P02672 | Fibrinogen alpha chain | FGA | 300.49 | 15 | 15 | 1.22E+07 |
| 68 | P37141 | Glutathione peroxidase 3 | GPX3 | 229.26 | 7 | 7 | 1.21E+07 |
| 69 | O18879 | Glutathione S-transferase A2 | GSTA2 | 148.7 | 4 | 3 | 1.20E+07 |
| 70 | Q2KJD0 | Tubulin beta-5 chain | TUBB5 | 360.2 | 20 | 4 | 1.06E+07 |
| 71 | P12799 | Fibrinogen gamma-B chain | FGG | 264.17 | 11 | 11 | 1.03E+07 |
| 72 | Q76LV1 | Heat shock protein HSP 90-beta | HSP90AB1 | 348.54 | 23 | 11 | 1.01E+07 |
| 73 | Q29RQ1 | Complement component C7 | C7 | 323.89 | 18 | 18 | 9.92E+06 |
| 74 | Q9TTJ5 | Regucalcin | RGN | 241.4 | 9 | 9 | 9.82E+06 |
| 75 | P12378 | UDP-glucose 6-dehydrogenase | UGDH | 290.35 | 16 | 16 | 9.40E+06 |
| 76 | Q76LV2 | Heat shock protein HSP 90-alpha | HSP90AA1 | 353.2 | 26 | 15 | 8.99E+06 |
| 77 | Q95M17 | Acidic mammalian chitinase | CHIA | 254.93 | 10 | 10 | 8.95E+06 |
| 78 | P19217 | Sulfotransferase 1E1 | SULT1E1 | 206.1 | 6 | 6 | 8.81E+06 |
| 79 | Q5EA79 | Galactose mutarotase | GALM | 265.7 | 7 | 7 | 8.78E+06 |
| 80 | P17690 | Beta-2-glycoprotein 1 | APOH | 265.16 | 9 | 9 | 8.36E+06 |
| 81 | P00741 | Coagulation factor IX | F9 | 242.92 | 10 | 10 | 7.92E+06 |
| 82 | P19879 | Mimecan | OGN | 214.97 | 8 | 8 | 7.54E+06 |
| 83 | Q5EAD2 | D-3-phosphoglycerate dehydrogenase | PHGDH | 293.94 | 12 | 12 | 7.49E+06 |
| 84 | P81644 | Apolipoprotein A-II | APOA2 | 202.89 | 6 | 6 | 7.02E+06 |
| 85 | A7E3W2 | Galectin-3-binding protein | LGALS3BP | 240.19 | 10 | 10 | 6.86E+06 |
| 86 | P01044 | Kininogen-1 | KNG1 | 331.08 | 21 | 7 | 6.83E+06 |
| 87 | P16116 | Aldo-keto reductase family 1 member B1 | AKR1B1 | 229.55 | 10 | 9 | 6.54E+06 |
| 88 | Q08DP0 | Phosphoglucosmutase-1 | PGM1 | 288.05 | 12 | 12 | 6.42E+06 |
| 89 | E1BF81 | Corticosteroid-binding globulin | SERPINA6 | 233.44 | 7 | 7 | 6.19E+06 |
| 90 | Q3ZC42 | Alcohol dehydrogenase class-3 | ADH5 | 274.43 | 11 | 11 | 5.89E+06 |
| 91 | Q5E9F7 | Cofilin-1 | CFL1 | 247.13 | 8 | 7 | 5.45E+06 |
| 92 | P02070 | Hemoglobin subunit beta | HBB | 228.08 | 8 | 4 | 5.43E+06 |
| 93 | Q5E9B7 | Chloride intracellular channel protein 1 | CLIC1 | 223.31 | 9 | 8 | 5.41E+06 |
| 94 | Q0VCM5 | Inter-alpha-trypsin inhibitor heavy chain H1 | ITIH1 | 290.35 | 12 | 12 | 5.40E+06 |
| 95 | P13605 | Fibromodulin | FMOD | 216.76 | 7 | 7 | 5.36E+06 |
| 96 | Q28035 | Glutathione S-transferase A1 | GSTA1 | 162.5 | 5 | 4 | 5.35E+06 |
| 97 | Q5EA01 | Beta-1 4-glucuronyltransferase 1 | B4GAT1 | 216.19 | 5 | 5 | 5.29E+06 |
| 98 | Q3SZD7 | Carbonyl reductase [NADPH] 1 | CBR1 | 254.05 | 10 | 10 | 5.07E+06 |
| 99 | Q3ZBD7 | Glucose-6-phosphate isomerase | GPI | 262.55 | 11 | 11 | 4.96E+06 |
| 100 | P50227 | Sulfotransferase 1A1 | SULT1A1 | 237.14 | 9 | 9 | 4.95E+06 |

N/A: non annotated gene.

Alpha-2-macroglobulin (accession ID Q7SIH1) is the first in the list, with an average area under the curve of 1.59 × 10^10^, followed by Albumin with an average area under the curve of 5.53 × 10^9^ (**Table 2** and **Table S3**). Interestingly, these proteins are typically enriched in the serum, in accordance with the presence of FBS in the starting samples [5,7].

To better classify the quantified proteins, the top 100 proteins were subjected to the Gene Ontology (GO) Enrichment Analysis using the BiNGO [23] plugin for Cytoscape software platform [24,25]. First, we determined the statistically over-represented GO aspect *Cellular Component* (CC) **(Table S4)**. In **Figure 4A** the GO terms with a corrected *p-*value < 0.001 are displayed; in particular, the most predominant ones are: (i) extracellular region (GO:0005576), (ii) extracellular space (GO:0005615), (iii) cytosol (GO:0005829). The *p*-values were calculated considering the number of proteins of the medium that are identified in the list of genes of a specific GO term, in relation to the genes in the whole bovine proteome that are annotated to that GO term. As expected, our samples contain mainly soluble proteins, most of which released in the extracellular environment, again in line with the presence of FBS. The “extracellular region” GO term includes 2,743 genes of the whole bovine genome (i.e., 28,791 genes), with a background frequency of 9.53%. In this GO term, we found 54 out of the 99 top proteins (1 protein was not involved in any statistically significant GO term), with a sample frequency of 54.55%. The “extracellular space” GO term has a background frequency of 6.12%, and included the 43.43% of medium proteins. The “cytosol” GO term has a background frequency of 12.83%, with a sample frequency of 36.36% **(Figure 4B)**. Indeed, the percentage of proteins deriving from these cellular compartments is higher than expected, supporting their enrichment in our samples.

**Figure 4.**
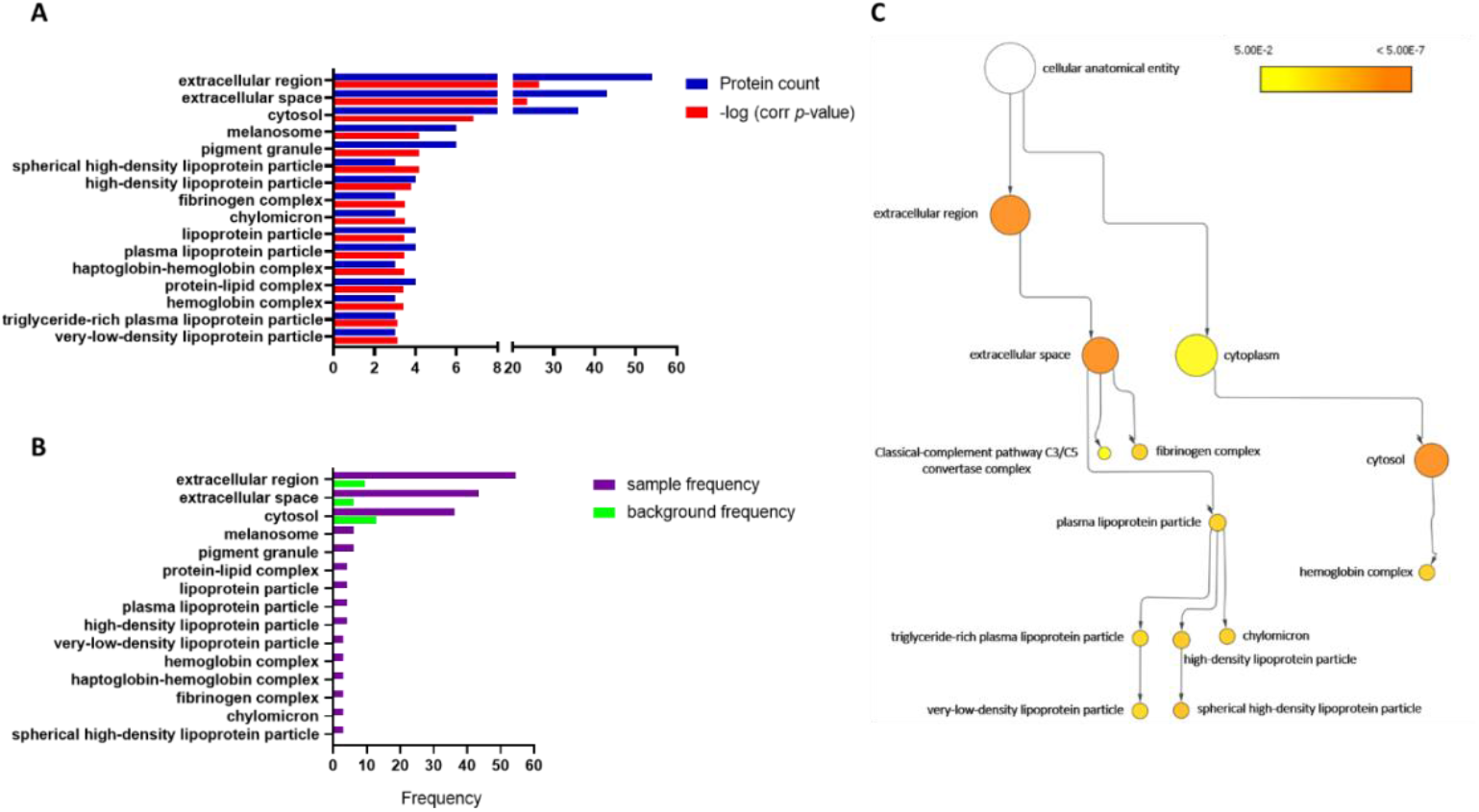
Gene Ontology analysis on the top 100 proteins quantified from LC-MS/MS proteomics. A) GO terms referring to cellular components found to be over-represented in medium proteins. Only GO terms with a corrected *p-*value < 0.001 were included in the graph. B) Graphical representation of background frequency (green bars: number of bovine proteins annotated to a specific GO term divided by the total number of bovine genes in the reference genome) and sample frequency (purple bars: number of medium proteins annotated to a specific GO term over the total number of proteins in our data set). C) Graphical visualization of the over-represented GO terms with parent–child relationships and color coding according to the *p*-value.

To better visualize the CC GO terms that we found over-represented, we used the graphical representations of BiNGO in the Cytoscape platform **(Figure 4C)**. The most significant GO terms showed in **Figure 4A** are linked by a parent–child relationship, with the first term (extracellular region) at the head of the tree. The other terms derived mainly from the extracellular space node and again, they included mainly serum-specific molecules **(Figure 4C)**.

### Membrane-related terms are under-represented in contaminant proteins

In addition, the same top 100 proteins were interrogated with BiNGO to define the under-represented CC GO terms **(Table S5)**. The results show a significant depletion of the terms “organelle” (GO: 043226), “intracellular organelle” (GO: 043229) and “membrane” (GO: 016020). It is noteworthy that only the latter has a corrected *p*-value < 0.01 **(Figure 5A)**. As expected, and contrary to what we found for the over-represented CC, these GO terms had a background frequency (membrane 48.67%, organelle 59.57% and intracellular organelle 58.05%) higher compared to the sample frequency (membrane: 25.25%, organelle: 38.38% and intracellular organelle: 38.38%) **(Figure 5B)**. This imply that in our dataset (i.e., the top 100 proteins quantified by PEAKS X-Pro software) these terms are less represented (lower sample frequency) compared to the genome (higher background frequency).

**Figure 5.**
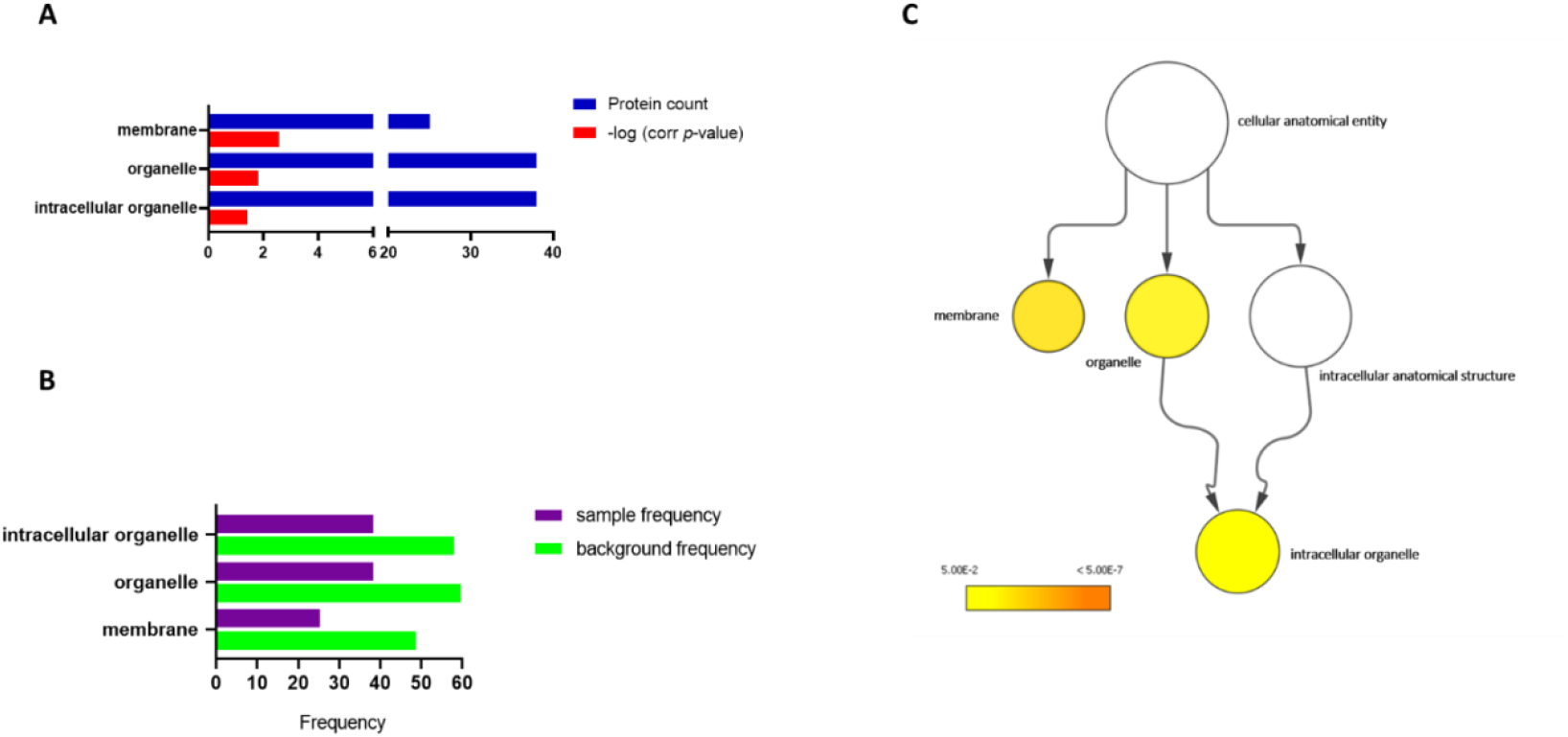
Gene Ontology analysis on the top 100 proteins quantified from LC-MS/MS proteomics. A) GO terms referring to cellular components found to be under-represented in medium proteins. GO terms with a corrected *p-*value < 0.05 were included in the graph. B) Graphical representation of background frequency (green bars: number of bovine proteins annotated to a specific GO term divided by the total number of bovine genes in the reference genome) and sample frequency (purple bars: number or medium proteins annotated to a specific GO term over the total number of proteins in our data set). C) Graphical visualization of the under-represented GO terms with parent– child relationships and color coding according to the *p-*value, obtained after the statistical analysis.

The visualization of the parent-child relationship between GO terms showed the link between organelle and intracellular organelle, with the term membrane as a separated node **(Figure 5C)**. Overall, considering the under-enrichment of membrane proteins (including EV terms), our results confirmed the limited vesicle contamination in the complete medium, in line with the use of EV-depleted FBS.

## Discussion

In the last years MS has expanded its applicability to proteomics studies of the cell secretome [26–28]. Cell secretome can be divided in two broad constituents, a soluble fraction and a vesicular fraction associated with EVs, key mediators of cell-to-cell communication [29,30]. In this context, a major challenge lies in distinguishing the contaminant proteins (usually derived from the FBS in the culture medium) from those of the organism under investigation, considering that even proteins belonging to phylogenetically distant organisms share extensive stretches of sequences. Indeed, the Unipept analysis of the 5,294 peptides identified in our complete medium samples by searching the *Bos taurus* database, indicated that 4,489 of total peptides are shared by all *Eukariota*, and only 219 peptides of them are specific for the *Bos* genus **(Figure 3)**. Moreover, proteins are indirectly identified through the corresponding peptides and, especially in higher eukaryotes, many of these peptide sequences can be assigned to more than one protein, making difficult the discrimination between the sample proteins and the contaminating ones. This issue is part of a much broader context where, in general, a typical shotgun LC-MS/MS experiment is challenged by environmental contaminants that may be introduced accidentally into the samples during the procedure. Contamination can arise from various sources, such as laboratory reagents, sample handling procedures, equipment and airborne particles. While non-protein contaminating molecules (as detergents, nucleic acids and salts) can be easily removed by the PlusOne 2-D Clean-Up kit [31] or using other chemical extraction methods, it is much more difficult to remove protein contaminants from a sample. Keratins, protein digestion enzymes (as trypsin and chymotrypsin of various sources), antibodies, affinity tags (as streptavidin), bovine serum albumin (which can be used as a blocking agent in pull-down workflows) and molecular standards for quantification can all be found on the surfaces in proteomics laboratories. Additionally, components of the culture medium may introduce bovine protein contamination into the cellular secretome and EV samples. These bovine contaminants constitute a relevant problem, as their concentration may be much higher than that of EV-proteins. In fact, highly concentrated protein contaminants can compete with sample proteins in the MS analysis and hide the detection of minority proteins present in low concentrations in the biological sample. For these reasons, an efficient method capable of identifying the sample-specific components needs to be developed. A targeted-MS analysis with an exclusion list may be used in DDA proteomics, but in this way peptides of the sample with similar retention time and m/z could be excluded from the analysis, especially without a high resolution mass analyzer [2].

An alternative approach to targeted-MS is the use of different sequence databases [32]. For example, if the sample is represented by murine EVs from *ex vivo*/*in vitro* cell cultures, the MS/MS data can be compared against various databases including *Mus musculus* (reference), *Bos taurus* (FBS contaminants) and a database consisting of both taxonomies. Furthermore, most of the software used for the analysis of raw MS data allow to indicate, alongside a reference database, also a database of protein sequences considered contaminants. Several resources exist that attempted to create generic lists of protein contaminants, such as (not an exhaustive list) the cRAP collection (https://www.thegpm.org/crap/), the PRIDE Peptidome spectral libraries (https://www.ebi.ac.uk/pride/spectrumlibrary) and the MaxQuant contaminants list (embedded in each MaxQuant version, https://www.maxquant.org/download_asset/maxquant/latest, as the Contaminants.fasta file). However, these databases do not accomplish our needs to carefully distinguish, for instance, between bovine and mouse proteins. Also, care must be taken to check the contents of these publicly available resources, since what is considered to be a contaminant in one case might not be a contaminant in another situation and, as such, valuable PSMs might be lost. Another important factor to consider is the update rate of these resources, encompassing changes to amino acid sequences, corresponding accession numbers or identifiers, and the incorporation of newly available sequences from updated or newly released nucleotide-level data. As an example, for the three resources indicated above, an update would be highly recommended, since they all look like years old.

These considerations collectively support the rationale for creating-and-maintaining a custom list of contaminants suited around the specific scientific question(s) under study, and aligned with the particular sample preparation protocols employed in each experimental context. Hence the idea of developing SPROUTS, a new contaminant database that can be specifically, but not only, used by scientists working in the EV field. SPROUTS_DB has been curated starting from complete DMEM medium supplemented with 10% EV-depleted FBS, and it should be applied primarily to *ex vivo*/*in vitro* approaches. This medium is one of the most used formulations to culture both primary cells (e.g., fibroblasts, neurons, glial cells etc.) and cell lines (e.g., HeLa, 293T, Cos-7 etc.), thus further broadening the potential use of SPROUTS_DB in several contexts. In the case of biofluid-derived samples (e.g., blood, urine etc.), the presence of additional or alternative contaminants is to be expected, although some degree of overlap with entries in SPROUTS_DB is likely, in particular when processing hematic samples.

Through a shotgun proteomic approach and nanoUHPLC/High Resolution nanoESI-MS/MS, we evaluated the presence of bovine-derived proteins, both soluble as well as from residual FBS-EVs in complete medium (without cells). Also, we compared two commercially available EV-depleted FBS, to identify eventual variations depending on different brands. Of note, even if the EV-free FBS preparations were obtained from two companies that used different production protocols, we found a relevant 90% of overlap in terms of identified proteins, further strengthening the accuracy of SPROUTS_DB.

Raw data analysis was performed by querying both sections of UniProt with taxonomy *Bos taurus*. The result of the complete medium profiling was a list of 1,108 proteins, 547 of which identified in SwissProt section and 561 in TrEMBL section. We merged these proteins with the non-redundant 180 proteins of Contaminants Database of MPI, finally obtaining a novel database of contaminants with 1,288 entries. This database, which includes bovine proteins (due to the presence of FBS) and common laboratory contaminants such as keratins, can be considered an improved version of the cRFP, created by Shin et al., in 2019 [22]. SPROUTS_DB showed an increased ability to discriminate contaminating bovine proteins, reducing the number of false identifications in cell secretome studies. The GO Enrichment Analysis performed with BiNGO on the top 100 proteins quantified with our proteomics approach revealed that the most represented Cellular Component categories were specifically related to extracellular milieu. Among the most abundant proteins, we identified albumin (the second in **Table 2**) and other “classic” contaminants for EVs [5,7]. This result was expected, considering the nature (i.e., the presence of FBS) and processing (i.e., UC) of the sample. Even if UC is still one of the most used methodologies for EV purification [5–10], other EV isolation strategies are emerging in the field, that could result in the recovery of different contaminants. In particular, SEC-based approaches are widely recognized as cleaner methods, where the contamination from FBS-derived proteins is reduced [4].

Another critical point in the field is the presence of proteins which co-isolate with EVs, surrounding them on the surface and contributing to their functions, as a dynamic EV corona. In particular, nine “EV corona proteins” were shown to be shared among EVs, viruses and artificial nanoparticles in blood [33]. Five of these (i.e., ApoA1, ApoB, ApoE, complement factor 3 and fibrinogen α-chain) were identified in our SPROUTS_DB. The biological significance of these EV corona proteins remains to be further investigated.

Interestingly, we found that membrane components were under-represented in the BiNGO analysis, in line with the EV-free FBS media used to generate SPROUTS_DB. Thus, our data support the suitability of these media for EV-based studies. However, some proteins often found to be incorporated in EVs - such as GAPDH and HSP70 - are present in our database [20], possibly related to the presence of residual bovine EVs in the EV-depleted FBS preparations. As previously mentioned, proteins listed in SPROUTS_DB - whether soluble or EV-derived - are considered contaminants when serum-containing media are used to culture human or rodent cells for secretome and EV studies. SPROUTS_DB facilitates the discrimination between residual bovine-derived proteins and those genuinely originating from the cells of interest. The decision to retain or exclude a given protein from the dataset can be guided by the relative peptide abundance from each species (e.g., bovine vs. human or mouse), thereby informing the degree of confidence in protein attribution. For instance, when a protein of interest is identified by peptides that are not shared with contaminant proteins, or by marker peptides associated uniquely with a single protein, the use of the SPROUTS database ensures that this protein is not excluded from the final list of proteins present in the sample. On the contrary, a protein is excluded when all of its identified peptides are also found in a contaminant protein, and consequently it must be discarded.

## Conclusions

SPROUTS_DB represents a valuable resource for improving the accuracy of protein identification in samples of interest, by enabling the recognition and exclusion of contaminants, in particular for EV-enriched samples. Considering the importance of EVs for both biomarker discovery and nanotherapeutic applications, this database represents an important tool available for the field. SPROUTS_DB was generated in FASTA format, to be read directly by software used to process proteomics data. The file can be freely downloaded from the Supplementary Information section. Furthermore, the complete list of entries is reported in **Tables S1 and S2**. Future perspective will focus on implementing an automated algorithm to expand and update the database, thus maximizing SPROUTS_DB performance.

To our knowledge, SPROUTS is the most updated and complete database of contaminants suitable for secretome proteomics studies, with a specific focus on EV research.

## Methods

### Complete medium preparation

DMEM (1 g L^−1^ glucose, Sigma Aldrich, D6046) was supplemented with 2 mM L-glutamine (Sigma Aldrich, G7513), 2.5 µg mL^−1^ amphotericin B (Sigma Aldrich, A2942), 1% penicillin/streptomycin (Sigma Aldrich, P0781), and 10% exosome-depleted FBS: (i) System Biosciences, EXO-FBS-250A-1; and (ii) ThermoFisher Scientific, Gibco A2720803. 50 mL of complete medium (without cells) was incubated for 24 h at 37 °C and 5% CO_2_ in 10 cm dishes (Corning, 353803). Then the medium was subjected to the same passages for EV isolation as in Leggio L. et al., 2022.[15] Briefly, the medium was collected and immediately centrifuged at 1,000 ×g at 4 °C for 15 min. Next, the supernatant was subjected to ultracentrifugation in a Sorvall WX100 (Thermo Scientific). The first ultracentrifugation was performed at 100,000 ×g at 4 °C for 75 min, in 2 ultra-cone polyclear centrifuge tubes (Seton, 7067), using the swing-out rotor SureSpin 30 (k-factor: 216, RPM: 23,200). Then the 2 pellets were washed with cold PBS 1× and ultracentrifuged again at the same speed for 40 min in a single thick wall polycarbonate tube (Seton, 2002), using the fixed-angle rotor T-8100 (k-factor: 106, RPM: 41,000). The resulting small pellets were dissolved in 100 µL of 50 mM ammonium bicarbonate (pH 8.3) and subjected to proteomics analysis.

### Proteomics sample preparation

Three biological replicates of complete medium were purified from non-protein contaminating molecules with the Plus One 2-D Clean-Up kit (GE Healthcare Life Sciences,80-6484-51) according to the manufacturer’s instructions. The desalted protein pellet of each sample was suspended in 50 µL of 50 mM ammonium bicarbonate (pH 8.3) (Fluka BioChemika, 09830) and incubated at 4°C for 15 min. Next, 50 µL of 0.2% RapiGest SF (Waters, 186001861) in 50 mM ammonium bicarbonate (pH 8.3) were added and the samples were incubated on ice for 30 min.[36–38] The amount of protein was determined by fluorometric assay using the Qubit Protein Assay kit, and 20 µg of each sample were analyzed (ThermoFisher Scientific, Q33211).[39] Each sample was reduced for 3 h at 25 °C by adding 44 µg of dithiothreitol (DTT) (Sigma, 53815) in 50 mM ammonium bicarbonate with intermittent mixing (pH 8.3): this corresponds to a 50:1 (mol/mol) excess with respect to the estimated protein thiol groups. Alkylation was performed by adding iodoacetamide (IAA) (Sigma, I1149) at 2:1 M ratio with respect to DTT in 50 mM ammonium bicarbonate (pH 8.3) for 1 h in the dark at 25 °C with intermittent mixing. Finally, the reduced and alkylated proteins of each sample were in-solution digested overnight using porcine trypsin (Sequencing Grade Modified Trypsin, Porcine, lyophilized, Promega, 0000546115) at an enzyme/substrate ratio of 1:50 at 37 °C, in 50 mM ammonium bicarbonate (pH 8.3). Samples obtained by in-solution tryptic digestion were dried under vacuum and then reconstituted in 30 µL of 5% formic acid (FA) (Honeywell/Fluka, 94318) aqueous solution. In order to obtain a final concentration of 25 ng µL^−1^, each sample solution was diluted five times with 5% FA aqueous solution.

### Liquid chromatography with tandem mass spectrometry (LC–MS/MS) analysis

MS data were acquired in triplicate for each sample assayed on an Orbitrap Fusion Tribrid (Q-OT-qIT) mass spectrometer (ThermoFisher Scientific, Bremen, Germany) equipped with a ThermoFisher Scientific DionexUltiMate 3000 RSLCnano system (Sunnyvale, CA), to assess the reproducibility of the available MS data, as described in Pittalà M.G.G. et al., 2020 [40]. In details, 1 μL of each sample was loaded onto an Acclaim®Nano Trap C18 column (100 μm i.d. × 2 cm, 5 μm particle size, 100 Å, ThermoFisher Scientific, PN 164564-CMD). After washing the trapping column with solvent A (H_2_O + 0.1% FA) for 3 min at a flow rate of 7 μL/min, the peptides were eluted from the trapping column onto a PepMap® RSLC C18 EASY Spray, 75 μm × 50 cm, 2 μm, 100 Å column (ThermoFisher Scientific, PN ES903) and were separated by elution at a flow rate of 0.250 μL/min, at 40 °C, with a linear gradient of solvent B (CH_3_CN+0.1% FA) in A, 5% for 3 min, followed by 5% to 65% in 82 min, followed by 65% to 95% in 5 min, holding 95% B 5 min, 95% to 5% in 10 min and re-equilibrating at 5% B for 25 min. Eluted peptides were ionized by a nanospray (Easy-spray ion source, Thermo Scientific) using a spray voltage of 1.7 kV and introduced into the mass spectrometer through a heated ion transfer tube (275 °C). Survey scans of peptide precursors in the m/z range 400– 1600 were performed at a resolution of 120,000 (@ 200 m/z) with an AGC target for the Orbitrap survey of 4.0 × 10^5^ and a maximum injection time (MaxIT) of 50 ms. Tandem MS was performed by isolation at 1.6 Th with the quadrupole, and high energy collisional dissociation (HCD) was performed in the Ion Routing Multipole (IRM), using a normalized collision energy of 35 and rapid scan MS analysis in the ion trap. Only those precursors with charge state 1–3 and intensity above the threshold of 5.0 ×·10^3^ were sampled for MS^2^. The dynamic exclusion duration was set to 60 s with a 10 ppm tolerance around the selected precursor and its isotopes. Monoisotopic precursor selection was turned on. AGC target and MaxIT for MS/MS spectra were 1.0 × 10^4^ and 100 ms, respectively. The instrument was run in top speed mode with 3 s cycles, meaning the instrument would continuously perform MS^2^ events until the list of non-excluded precursors diminishes to zero or 3 s, whichever is shorter. MS/MS spectral quality was enhanced enabling the parallelizable time option (i.e., by using all parallelizable time during full scan detection for MS/MS precursor injection and detection). Mass spectrometer calibration was performed using the Pierce® LTQ Velos ESI Positive Ion Calibration Solution (ThermoFisher Scientific, 88323). MS data acquisition was performed using the Xcalibur v. 3.0.63 software (ThermoFisher Scientific, Milan, Italy).

### Database search

LC-MS/MS data were processed using PEAKS *de novo* sequencing software for data analysis (v. X-Pro, Bioinformatics Solutions Inc., Waterloo, ON, Canada). The data were searched against the 6,035 entries “*Bos taurus*” SwissProt database (release October 2022) and against the 41,097 entries “*Bos taurus*” TrEMBL database (release October 2022). The common Repository of Adventitious Proteins (c-RAP) contaminant database was enabled in the database search. Full tryptic peptides with a maximum of 3 missed cleavage sites were subjected to a bioinformatic search. Cysteine carboxyamidomethylation was set as the fixed modification, whereas oxidation of methionine, transformation of N-terminal glutamine and N-terminal glutamic acid residues in the form of pyroglutamic acid and N-terminal protein acetylation were included as variable modifications. The precursor mass tolerance threshold was 10 ppm and the maximum fragment mass error was set to 0.6 Da. Peptide spectral matches (PSM) were validated using Target Decoy PSM Validator node based on q-values at a 0.1% False Discovery Rate (FDR). Proteins codified from different genes though containing the same peptides which could not be differentiated based on MS/MS analysis alone were considered. Finally, all the reviewed protein hits obtained were processed by using the inChorus function of PEAKS. This tool combines the database search results of PEAKS software with those obtained by the Mascot search engine. This strategy allows not only to increase the coverage, but also the confidence of the protein identification [41], since the engines use independent algorithms and therefore the results confirm each other. In a single database search, a protein was considered identified if it fulfilled both of the following requirements: (i) a minimum of two peptides with a score above the peptide filtering threshold matches; (ii) the list of the matched peptides with at least a unique peptide (i.e., marker peptide). In order to produce the final list of proteins, only those identified at least in two out of three LC-MS/MS technical replicates and at least in two out of three biological replicates were considered. The MS proteomics data have been deposited to the ProteomeXchange Consortium via the PRIDE[42] partner repository with the dataset identifier PXD044137.

### GO enrichment analysis

Gene ontology (GO) term enrichment analysis to find statistically over- and under-represented categories was carried out with the open-source BiNGO [23] (a Biological Network Gene Ontology tool) 3.0.5 as a plugin for Cytoscape 3.9.1. [24]. The ontology file in OBO Flat File format 1.4 (release 2023-01-01) and the *Bos taurus* annotations file in GAF format (release 2022-11-12) were downloaded from Gene Ontology (http://geneontology.org/docs/download-ontology/) and Gene Ontology Annotation (GOA) (https://www.ebi.ac.uk/GOA/proteomes) websites, respectively.

The whole *Bos taurus* GOA annotation file was used as a reference set and GO terms referring to the cellular component terms were searched. Hypergeometric test was selected as statistical test and Benjamini & Hochberg False Discovery Rate (FDR) correction as multiple testing correction.

In order to visualize in Cytoscape the over- and under-represented categories after correction, a significance level of 0.05 was chosen.

In Cytoscape, yFiles Hierarchic Layout was selected and finally the most significant nodes were extracted from the entire networks and displayed in the graphs.

### Metaproteomics analysis at peptide level

The metaproteomics analysis was performed through the open-source web application Unipept (Unipept 5.0.8; http://unipept.ugent.be) [21], using all peptides identified in the complete medium with a peptides score (−10lgP) ≥ 43.8 at 0.1% FDR [43,44].

This tool for metaproteomics analysis, realized for tryptic peptides obtained with a shotgun approach, calculate the Lowest Common Ancestors (LCA) of a group of peptides and shows the most specific taxonomic level for each peptide. Ubiquitarian peptides are instead assigned to “organism”.

## Acknowledgements

The work of EA for this publication is not related in any way to Abzena. The authors are grateful to Bianca Marchetti and Salvatore Foti for their valuable support. LL, GP and NI are grateful to the Pharmacology section at the BIOMETEC Department, which hosts the authors’ laboratory. The authors also acknowledge the PON project Bio-nanotech Research and Innovation Tower (BRIT), financed by the Italian Ministry for Education, University and Research (MIUR) (Grant no. PONa3_00136). They are grateful for the technical assistance of the BRIT Team with the ultracentrifuge and the Orbitrap Fusion mass spectrometer.

## Author contributions

The manuscript was written through contributions of all authors. All authors have given approval to the final version of the manuscript. ‡These authors contributed equally.

## Funding

The project has been supported by the “Brain to South” grant (Fondazione con il Sud – Bando Capitale Umano ad Alta Qualificazione 2015). The research program also received support from: the University of Catania, “Bando-Chance”, Ph.D. program in Biotechnology, “Piano della Ricerca di Ateneo PIACERI 2020” grants “ARVEST” and “VDAC”; UniPD (STARs 2019: Supporting TAlents in ReSearch); the Italian Ministry of Health (GR-2016-02363461); and the IRCCS San Camillo Hospital.

## Data Availability

Data are available via ProteomeXchange with the identifier PXD044137. Other data that support the findings of this study are available from the corresponding authors upon reasonable request.

## Competing interests

The authors declare no conflict of interest.

## Notes

https://www.proteomexchange.org

### Competing Interest Statement

The authors have declared no competing interest.

