## Supplementary materials for "SPROUTS_DB: an implemented database of contaminants for extracellular vesicle proteomics studies"

S. Vivarelli
Department of Biomedical and Dental Sciences and Morphofunctional Imaging, Occupational Medicine Section, University of Messina, Messina, Italy

E. Alpi
Abzena, Babraham Research Campus, Cambridge, United Kingdom.

‡ MGGP and LL contributed equally to this work.

**Analysis to test the performance of SPROUTS_DB**

To evaluate the performance of SPROUTS_DB, we used the MS/MS raw data files from Shin et al., 2019[1] (PRIDE repository, identifier PXD015143). In this work, the authors proposed the use of cRFP (Common Repository of FBS Proteins) for the identification of contaminants. We compared the results between searches obtained against the Human DB (Search 1), the Human DB plus cRFP DB (Search 2), and the Human DB plus SPROUTS_DB (Search 3).

LC-MS/MS data were processed using PEAKS *de novo* sequencing software for data analysis (v. X-Pro, Bioinformatics Solutions Inc., Waterloo, ON, Canada). The data were searched using the reference proteome of Homo sapiens (104,583 entries, release July 2024) as reference database and cRFP or SPROUTS as contaminant database. Full tryptic peptides with a maximum of 3 missed cleavage sites were subjected to a bioinformatic search. Cysteine carboxyamidomethylation was set as the fixed modification, whereas oxidation of methionine, transformation of N-terminal glutamine and N- terminal glutamic acid residues in the form of pyroglutamic acid and N-terminal protein acetylation were included as variable modifications. The precursor mass tolerance threshold was 10 ppm and the maximum fragment mass error was set to 0.6 Da. Peptide spectral matches (PSM) were validated using Target Decoy PSM Validator node based on q-values at a 1% False Discovery Rate (FDR), as in Shin 2019[1]. In a single database search, a protein was considered identified if it fulfilled both of the following requirements: (i) a minimum of two peptides with a score above the peptide filtering threshold matches; (ii) at least a unique peptide in the list of the matched peptides. In **Table 1** of the main text (reported for clarity also below) the results of these analyses are shown.

| Raw Data | Search | Reference Database | Contaminant Database | Human Proteins | Potential contaminants |
|---|---|---|---|---|---|
| 160210_exo_4th_3_mock_01<br>160210_exo_4th_3_mock_02<br>160210_exo_4th_3_mock_03<br>160210_exo_4th_3_mock_04 | 1 | Reference Proteome_Human (104,583 entries) | / | 1,369 | / |
| | 2 | Reference Proteome_Human (104,583 entries) | cRFP (199 entries) | 1,369 | 56 |
| | 3 | Reference Proteome_Human (104,583 entries) | SPROUTS_DB (1,288 entries) | 1,347 | 267 |

The results of Searches 1 and 2 both show a total of 1,369 identified proteins. The comparison of the protein lists shows that although 1,315 proteins are common, 54 proteins vary between the two searches.

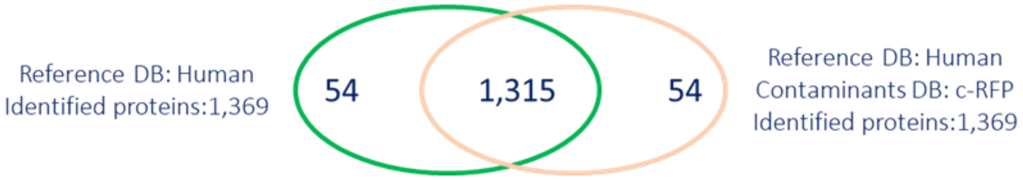


In particular, among the 54 proteins (in the green circle), 11 "human" proteins of the Search 1 correspond to "bovine" homologous proteins identified in the presence of cRFP DB (Search 2), with more peptides and PSMs (**Table A**). According Occam’s razor, we can conclude that these 11 proteins are not human but bovine contaminants.

The same assumption is valid for another 25 proteins identified exclusively in Search 1 which belong to the same protein groups as the 11 proteins reported in **Table A**, identified with the same set of peptides.

The remaining 18 "human" proteins are different in the two searches. Indeed, PSMs that allow the unique identification of a protein in Search 1 may contribute to the identification of a different protein in Search 2. Also, peptides that are unique to a protein in the Human DB, they are not unique in the presence of cRFP. Overall, the addition of the cRFP DB allows the identification of 56 contaminant proteins in Search 2.

**Table A.** Details of identified proteins using Human Database (Search 1) and Human Database with c-RFP as Contaminants Database (Search 2). For each protein is reported: protein name, gene code, number of identified peptides and of PSMs in Search 1 and in Search 2, respectively.

| Protein Group | Reference Database: Human | | | | Reference Database: Human; Contaminants Database: cRFP | | | |
|---|---|---|---|---|---|---|---|---|
| | Human | | | | Bovine | | | |
| | Protein name | Gene Code | N. of peptides | PSMs | Protein name | Gene Code | N. of peptides | PSMs |
| 1 | Coagulation factor V | F5 | 9 | 16 | Coagulation factor V | F5 | 38 | 66 |
| 2 | Prothrombin | F2 | 8 | 29 | Prothrombin | F2 | 21 | 75 |
| 3 | Tetranectin | CLEC3B | 7 | 9 | Tetranectin | CLEC3B | 12 | 16 |
| 4 | Albumin | ALB | 6 | 14 | Serum albumin | ALB | 43 | 101 |
| 5 | Tubulin beta-1 chain | TUBB1 | 6 | 18 | Uncharacterized protein | TUBB1 | 8 | 23 |
| 6 | Alpha-2-HS-glycoprotein | AHSG | 3 | 8 | Alpha-2-HS-glycoprotein | AHSG | 15 | 66 |
| 7 | Hemoglobin subunit beta | HBB | 3 | 16 | Hemoglobin subunit beta | HBB | 5 | 22 |
| 8 | Sex hormone-binding globulin | SHBG | 2 | 2 | SHBG protein | SHBG | 8 | 10 |
| 9 | Periostin | POSTN | 2 | 3 | Periostin variant 7 | POSTN | 3 | 4 |
| 10 | Inter-alpha-trypsin inhibitor heavy chain H2 | ITIH2 | 2 | 4 | Inter-alpha-trypsin inhibitor heavy chain H2 | ITIH2 | 17 | 35 |
| 11 | Alpha-2-antiplasmin | SERPINF2 | 2 | 4 | Alpha-2-antiplasmin | SERPINF2 | 9 | 16 |

Similarly, we compared Search 1 and Search 3:

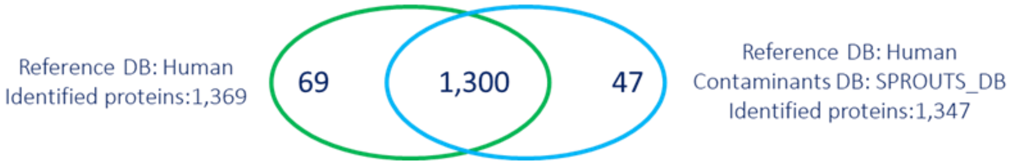


The presence of SPROUTS_DB leads to the identification of a smaller number of human proteins (1,347 proteins of which 47 identified exclusively in Search 3), and a greater number (i.e., 267) of potentially contaminant bovine proteins.

Also, among the 69 proteins (in the green circle), in addition to the 11 proteins present in Table A, SPROUTS_DB allows to identify two other proteins as contaminants, indicated in Table B. Again, bovine proteins are identified with more peptides and PSMs; based on Occam's razor, also these proteins are potential contaminants.

**Table B.** Details of the two proteins identified as contaminants exclusively using SPROUTS_DB (Search 3). For each protein is reported: protein name, gene code, number of identified peptides and of PSMs in Search 1 and in Search 3, respectively.

| Protein Group | Reference Database: Human | | | | Reference Database: Human; Contaminants Database: SPROUTS_DB | | | |
|---|---|---|---|---|---|---|---|---|
| | Human | | | | Bovin | | | |
| | Protein name | Gene Code | N. of peptides | PSMs | Protein name | Gene Code | N. of peptides | PSMs |
| 12 | Gamma-glutamyl hydrolase | GGH | 2 | 3 | Gamma-glutamyl hydrolase | GGH | 7 | 13 |
| 13 | Tubulin beta-1 chain | TUBB1 | 6 | 18 | Uncharacterized protein | TUBB1 | 9 | 23 |

Furthermore, 29 of the proteins identified exclusively in Search 1 belong to the same protein groups as the proteins in Tables A and B, identified with the same set of peptides. All of these proteins have a bovine homolog protein identified in Search 3 with more peptides and PSMs. The remaining "human" proteins (27 out of 69 in Search 1) are different.

In light of these results, we can conclude that: (i) the higher number of the protein contaminants; and (ii) the lower number of human proteins identified in Search 3 correlates with a greater performance of SPROUTS_DB in discriminating bovine contaminants, thus contributing to the correct identification of proteins in the sample of interest.

### FIGURES

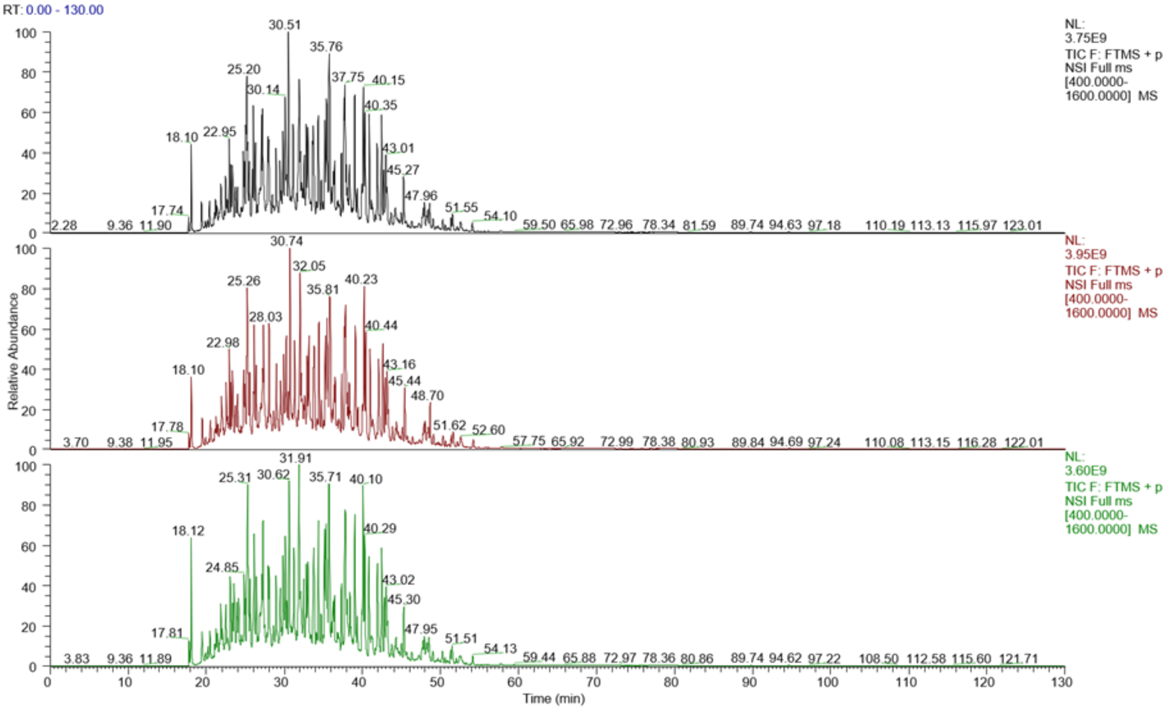


**Figure S1**. Total Ion Current (TIC) chromatograms of the 3 technical replicates of biological replicate 1.

### TABLES

**Table S1.** List of proteins identified in the complete medium. For each protein is reported: Accession Number as in the UniProt database, protein name, gene code, average mass and UniProt section. The 65 common proteins between complete medium and Contaminants database of the Max Planck Institute (MPI) are highlighted in bold. The 35 proteins of the exosome-depleted FBS from ThermoFisher Scientific (Gibco A2720803) are underlined.

| Accession | Protein name | Gene code | Avg. Mass | Section of UniProt |
|---|---|---|---|---|
| A1A4J1 | ATP-dependent 6-phosphofructokinase liver type | PFKL | 85292 | SwissProt |
| A1A4R1 | Histone H2A type 2-C | H2AC20 | 13988 | SwissProt |
| A1L595 | Keratin type I cytoskeletal 17 | KRT17 | 48712 | SwissProt |
| A2I7M9 | Serpin A3-2 | SERPINA3-2 | 46237 | SwissProt |
| **A2I7N1** | **Serpin A3-5** | **SERPINA3-5** | **46397** | **SwissProt** |
| A2I7N2 | Serpin A3-6 | SERPINA3-6 | 46390 | SwissProt |
| **A2I7N3** | **Serpin A3-7** | **SERPINA3-7** | **46942** | **SwissProt** |
| A2VDL6 | Sodium/potassium-transporting ATPase subunit alpha-2 | ATP1A2 | 112179 | SwissProt |
| A2VDS1 | Selenocysteine lyase | SCLY | 47204 | SwissProt |
| A3KMV5 | Ubiquitin-like modifier-activating enzyme 1 | UBA1 | 117830 | SwissProt |
| A3KN12 | Adenylosuccinate lyase | ADSL | 55484 | SwissProt |
| A4FUA8 | F-actin-capping protein subunit alpha-1 | CAPZA1 | 32932 | SwissProt |
| A4FV54 | Ras-related protein Rab-8A | RAB8A | 23684 | SwissProt |
| A4IF69 | NHL repeat-containing protein 2 | NHLRC2 | 79314 | SwissProt |
| A5D785 | Exportin-2 | CSE1L | 110375 | SwissProt |
| A5D7I4 | Exostosin-1 | EXT1 | 86299 | SwissProt |
| A5PJI5 | ATPase GET3 | GET3 | 38793 | SwissProt |
| A5PK51 | Nicotinate phosphoribosyltransferase | NAPRT | 57871 | SwissProt |
| A6H742 | Plastin-1 | PLS1 | 70532 | SwissProt |
| A6H767 | Nucleosome assembly protein 1-like 1 | NAP1L1 | 45376 | SwissProt |
| A6H768 | Galactokinase | GALK1 | 42227 | SwissProt |
| A6N9I4 | RAS guanyl-releasing protein 2 | RASGRP2 | 69180 | SwissProt |
| A6QLP2 | Adenosylhomocysteinase 3 | AHCYL2 | 66774 | SwissProt |
| A6QLR4 | Flotillin-2 | FLOT2 | 46995 | SwissProt |
| A6QPQ2 | Serpin A3-8 | SERPINA3-8 | 46964 | SwissProt |
| A6QR46 | Ras-related protein Rab-6B | RAB6B | 23462 | SwissProt |
| A6QR56 | Aldehyde dehydrogenase family 16 member A1 | ALDH16A1 | 85168 | SwissProt |
| A7E3W2 | Galectin-3-binding protein | LGALS3BP | 62127 | SwissProt |
| A7MAZ5 | Histone H1.3 | H1-3 | 22154 | SwissProt |
| A7MB62 | Actin-related protein 2 | ACTR2 | 44761 | SwissProt |

| | | | | |
|---|---|---|---|---|
| A7MBI7 | Catechol O-methyltransferase | COMT | 30484 | SwissProt |
| A7MBJ4 | Receptor-type tyrosine-protein phosphatase F | PTPRF | 211381 | SwissProt |
| A7MBJ5 | Cullin-associated NEDD8-dissociated protein 1 | CAND1 | 136375 | SwissProt |
| A7YW98 | Arginine--tRNA ligase cytoplasmic | RARS1 | 75561 | SwissProt |
| A7YWG4 | Gamma-glutamyl hydrolase | GGH | 35683 | SwissProt |
| A7YWP4 | Histidine ammonia-lyase | HAL | 72291 | SwissProt |
| D3K0R6 | Plasma membrane calcium-transporting ATPase 4 | ATP2B4 | 133792 | SwissProt |
| E1BB52 | Cyclin-dependent kinase 13 | CDK13 | 164716 | SwissProt |
| E1BF81 | Corticosteroid-binding globulin | SERPINA6 | 45102 | SwissProt |
| F1N152 | Serine protease HTRA1 | HTRA1 | 51907 | SwissProt |
| G3MYZ3 | Afamin | AFM | 69562 | SwissProt |
| O02659 | Mannose-binding protein C | MBL | 26471 | SwissProt |
| O02675 | Dihydropyrimidinase-related protein 2 | DPYSL2 | 62278 | SwissProt |
| O02739 | Serpin B6 | SERPINB6 | 42561 | SwissProt |
| O18879 | Glutathione S-transferase A2 | GSTA2 | 25717 | SwissProt |
| O46375 | Transthyretin | TTR | 15727 | SwissProt |
| O46414 | Ferritin heavy chain | FTH1 | 21052 | SwissProt |
| O46415 | Ferritin light chain | FTL | 19988 | SwissProt |
| O77742 | Osteomodulin | OMD | 49116 | SwissProt |
| O77783 | Exostosin-2 | EXT2 | 81887 | SwissProt |
| O77834 | Peroxiredoxin-6 | PRDX6 | 25067 | SwissProt |
| O97764 | Zeta-crystallin | CRYZ | 35383 | SwissProt |
| P00366 | Glutamate dehydrogenase 1 mitochondrial | GLUD1 | 61512 | SwissProt |
| P00376 | Dihydrofolate reductase | DHFR | 21604 | SwissProt |
| P00432 | Catalase | CAT | 59915 | SwissProt |
| P00435 | Glutathione peroxidase 1 | GPX1 | 22659 | SwissProt |
| P00514 | cAMP-dependent protein kinase type I-alpha regulatory subunit | PRKAR1A | 42893 | SwissProt |
| P00515 | cAMP-dependent protein kinase type II-alpha regulatory subunit | PRKAR2A | 45094 | SwissProt |
| P00517 | cAMP-dependent protein kinase catalytic subunit alpha | PRKACA | 40620 | SwissProt |
| **P00735** | **Prothrombin** | **F2** | **70506** | **SwissProt** |
| P00741 | Coagulation factor IX | F9 | 52046 | SwissProt |
| P00743 | Coagulation factor X | F10 | 54510 | SwissProt |
| P00744 | Vitamin K-dependent protein Z | PROZ | 43113 | SwissProt |
| P00745 | Vitamin K-dependent protein C (Fragment) | PROC | 51416 | SwissProt |
| P00794 | Chymosin | CYM | 42180 | SwissProt |

| | | | | |
|---|---|---|---|---|
| P00829 | ATP synthase subunit beta mitochondrial | ATP5F1B | 56284 | SwissProt |
| **P00978** | **Protein AMBP** | **AMBP** | **39235** | **SwissProt** |
| P01017 | Angiotensinogen | AGT | 51424 | SwissProt |
| **P01030** | **Complement C4 (Fragments)** | **C4** | **101908** | **SwissProt** |
| **P01044** | **Kininogen-1** | **KNG1** | **68890** | **SwissProt** |
| **P01045** | **Kininogen-2** | **KNG2** | **68710** | **SwissProt** |
| P01131 | Low-density lipoprotein receptor | LDLR | 92869 | SwissProt |
| P01267 | Thyroglobulin | TG | 303221 | SwissProt |
| **P01966** | **Hemoglobin subunit alpha** | **HBA** | **15184** | **SwissProt** |
| **P02070** | **Hemoglobin subunit beta** | **HBB** | **15954** | **SwissProt** |
| P02081 | Hemoglobin fetal subunit beta | N/A | 15859 | SwissProt |
| P02253 | Histone H1.2 | H1-2 | 21356 | SwissProt |
| P02453 | Collagen alpha-1(I) chain | COL1A1 | 138939 | SwissProt |
| P02459 | Collagen alpha-1(II) chain | COL2A1 | 141829 | SwissProt |
| P02465 | Collagen alpha-2(I) chain | COL1A2 | 129064 | SwissProt |
| **P02584** | **Profilin-1** | **PFN1** | **15057** | **SwissProt** |
| **P02672** | **Fibrinogen alpha chain** | **FGA** | **67012** | **SwissProt** |
| **P02676** | **Fibrinogen beta chain** | **FGB** | **53340** | **SwissProt** |
| P02722 | ADP/ATP translocase 1 | SLC25A4 | 32967 | SwissProt |
| **P02769** | **Albumin** | **ALB** | **69294** | **SwissProt** |
| **P04258** | **Collagen alpha-1(III) chain** | **COL3A1** | **93651** | **SwissProt** |
| P04272 | Annexin A2 | ANXA2 | 38612 | SwissProt |
| P04695 | Guanine nucleotide-binding protein G(t) subunit alpha-1 | GNAT1 | 39966 | SwissProt |
| P04696 | Guanine nucleotide-binding protein G(t) subunit alpha-2 | GNAT2 | 40144 | SwissProt |
| P04896 | Guanine nucleotide-binding protein G(s) subunit alpha isoforms short | GNAS | 45709 | SwissProt |
| P05131 | cAMP-dependent protein kinase catalytic subunit beta | PRKACB | 40594 | SwissProt |
| P05307 | Protein disulfide-isomerase | P4HB | 57266 | SwissProt |
| P05785 | Keratin type I cytoskeletal 14 (Fragment) | KRT14 | 10722 | SwissProt |
| P05786 | Keratin type II cytoskeletal 8 | KRT8 | 53627 | SwissProt |
| P05980 | Prostaglandin F synthase 1 | N/A | 36720 | SwissProt |
| P06394 | Keratin type I cytoskeletal 10 | KRT10 | 54848 | SwissProt |
| **P06868** | **Plasminogen** | **PLG** | **91216** | **SwissProt** |
| **P07224** | **Vitamin K-dependent protein S** | **PROS1** | **75133** | **SwissProt** |
| P07507 | Matrix Gla protein | MGP | 12217 | SwissProt |
| P07514 | NADH-cytochrome b5 reductase 3 | CYB5R3 | 34122 | SwissProt |
| P07589 | Fibronectin | FN1 | 272151 | SwissProt |
| P07688 | Cathepsin B | CTSB | 36661 | SwissProt |

| | | | | |
|---|---|---|---|---|
| P08037 | Beta-1 4-galactosyltransferase 1 | B4GALT1 | 44843 | SwissProt |
| P08169 | Cation-independent mannose-6-phosphate receptor | IGF2R | 274527 | SwissProt |
| P08239 | Guanine nucleotide-binding protein G(o) subunit alpha | GNAO1 | 40067 | SwissProt |
| P08728 | Keratin type I cytoskeletal 19 | KRT19 | 43885 | SwissProt |
| P0C7Q4 | Guanine nucleotide-binding protein G(t) subunit alpha-3 | GNAT3 | 40332 | SwissProt |
| P0CB32 | Heat shock 70 kDa protein 1-like | HSPA1L | 70389 | SwissProt |
| P0CG53 | Polyubiquitin-B | UBB | 34308 | SwissProt |
| P0CH28 | Polyubiquitin-C | UBC | 77570 | SwissProt |
| P10096 | Glyceraldehyde-3-phosphate dehydrogenase | GAPDH | 35868 | SwissProt |
| P11017 | Guanine nucleotide-binding protein G(I)/G(S)/G(T) subunit beta-2 | GNB2 | 37331 | SwissProt |
| P11064 | Low molecular weight phosphotyrosine protein phosphatase | ACP1 | 18054 | SwissProt |
| P11151 | Lipoprotein lipase | LPL | 53378 | SwissProt |
| P12260 | Coagulation factor XIII A chain (Fragment) | F13A1 | 22745 | SwissProt |
| P12378 | UDP-glucose 6-dehydrogenase | UGDH | 55136 | SwissProt |
| P12624 | Myristoylated alanine-rich C-kinase substrate | MARCKS | 31665 | SwissProt |
| **P12763** | **Alpha-2-HS-glycoprotein** | **AHSG** | **38419** | **SwissProt** |
| P12799 | Fibrinogen gamma-B chain | FGG | 50244 | SwissProt |
| P13384 | Insulin-like growth factor-binding protein 2 | IGFBP2 | 34015 | SwissProt |
| P13605 | Fibromodulin | FMOD | 43038 | SwissProt |
| P13696 | Phosphatidylethanolamine-binding protein 1 | PEBP1 | 20986 | SwissProt |
| P14568 | Argininosuccinate synthase | ASS1 | 46417 | SwissProt |
| P15246 | Protein-L-isoaspartate(D-aspartate) O-methyltransferase | PCMT1 | 24565 | SwissProt |
| **P15497** | **Apolipoprotein A-I** | **APOA1** | **30276** | **SwissProt** |
| P16116 | Aldo-keto reductase family 1 member B1 | AKR1B1 | 35919 | SwissProt |
| P17248 | Tryptophan--tRNA ligase cytoplasmic | WARS1 | 53812 | SwissProt |
| **P17690** | **Beta-2-glycoprotein 1** | **APOH** | **38252** | **SwissProt** |
| **P17697** | **Clusterin** | **CLU** | **51114** | **SwissProt** |
| P18902 | Retinol-binding protein 4 | RBP4 | 21069 | SwissProt |
| P19035 | Apolipoprotein C-III | APOC3 | 10692 | SwissProt |
| P19120 | Heat shock cognate 71 kDa protein | HSPA8 | 71241 | SwissProt |
| P19217 | Sulfotransferase 1E1 | SULT1E1 | 34640 | SwissProt |

| | | | | |
|---|---|---|---|---|
| P19483 | ATP synthase subunit alpha mitochondrial | ATP5F1A | 59720 | SwissProt |
| P19687 | Guanylate cyclase soluble subunit alpha-1 | GUCY1A1 | 77533 | SwissProt |
| P19803 | Rho GDP-dissociation inhibitor 1 | ARHGDIA | 23421 | SwissProt |
| P19858 | L-lactate dehydrogenase A chain | LDHA | 36598 | SwissProt |
| P19879 | Mimecan | OGN | 34209 | SwissProt |
| P20000 | Aldehyde dehydrogenase mitochondrial | ALDH2 | 56653 | SwissProt |
| P21809 | Biglycan | BGN | 41590 | SwissProt |
| P21856 | Rab GDP dissociation inhibitor alpha | GDI1 | 50566 | SwissProt |
| P22226 | Cathelicidin-1 | CATHL1 | 17600 | SwissProt |
| P22457 | Coagulation factor VII | F7 | 48961 | SwissProt |
| P23196 | DNA-(apurinic or apyrimidinic site) endonuclease | APEX1 | 35570 | SwissProt |
| P23206 | Collagen alpha-1(X) chain | COL10A1 | 65546 | SwissProt |
| P23956 | V-type proton ATPase 16 kDa proteolipid subunit c | ATP6V0C | 15720 | SwissProt |
| P24627 | Lactotransferrin | LTF | 78056 | SwissProt |
| P25975 | Procathepsin L | CTSL | 37347 | SwissProt |
| P26452 | 40S ribosomal protein SA | RPSA | 32884 | SwissProt |
| P26882 | Peptidyl-prolyl cis-trans isomerase D | PPID | 40620 | SwissProt |
| P27214 | Annexin A11 | ANXA11 | 54018 | SwissProt |
| P27479 | Polyunsaturated fatty acid lipoxygenase ALOX15 | ALOX15 | 75124 | SwissProt |
| P27674 | Solute carrier family 2 facilitated glucose transporter member 1 | SLC2A1 | 54132 | SwissProt |
| P28782 | Protein S100-A8 | S100A8 | 10460 | SwissProt |
| **P28800** | **Alpha-2-antiplasmin** | **SERPINF2** | **54711** | **SwissProt** |
| P28801 | Glutathione S-transferase P | GSTP1 | 23613 | SwissProt |
| P29702 | Protein farnesyltransferase/geranylgeranyltransferase type-1 subunit alpha | FNTA | 43845 | SwissProt |
| P30932 | CD9 antigen | CD9 | 25258 | SwissProt |
| P31081 | 60 kDa heat shock protein mitochondrial | HSPD1 | 61108 | SwissProt |
| P31404 | V-type proton ATPase catalytic subunit A | ATP6V1A | 68344 | SwissProt |
| P31754 | Uridine 5'-monophosphate synthase | UMPS | 52229 | SwissProt |
| P31976 | Ezrin | EZR | 68760 | SwissProt |
| P33433 | Histidine-rich glycoprotein (Fragments) | HRG | 44471 | SwissProt |
| P33672 | Proteasome subunit beta type-3 | PSMB3 | 22993 | SwissProt |

| | | | | |
|---|---|---|---|---|
| P34933 | Heat shock-related 70 kDa protein 2 | HSPA2 | 69740 | SwissProt |
| **P34955** | **Alpha-1-antiproteinase** | **SERPINA1** | **46104** | **SwissProt** |
| P35445 | Cartilage oligomeric matrix protein | COMP | 82362 | SwissProt |
| P37141 | Glutathione peroxidase 3 | GPX3 | 25663 | SwissProt |
| P38657 | Protein disulfide-isomerase A3 | PDIA3 | 56930 | SwissProt |
| **P41361** | **Antithrombin-III** | **SERPINC1** | **52347** | **SwissProt** |
| P41541 | General vesicular transport factor p115 | USO1 | 107515 | SwissProt |
| P42916 | Collectin-43 | CL43 | 33616 | SwissProt |
| P43033 | Platelet-activating factor acetylhydrolase IB subunit beta | PAFAH1B1 | 46613 | SwissProt |
| P46196 | Mitogen-activated protein kinase 1 | MAPK1 | 41376 | SwissProt |
| P47865 | Aquaporin-1 | AQP1 | 28800 | SwissProt |
| P48034 | Aldehyde oxidase 1 | AOX1 | 147611 | SwissProt |
| P48616 | Vimentin | VIM | 53728 | SwissProt |
| P48644 | Retinal dehydrogenase 1 | ALDH1A1 | 54806 | SwissProt |
| P49907 | Selenoprotein P | SELENOP | 45488 | SwissProt |
| P50227 | Sulfotransferase 1A1 | SULT1A1 | 34184 | SwissProt |
| P50397 | Rab GDP dissociation inhibitor beta | GDI2 | 50488 | SwissProt |
| **P50448** | **Factor XIIa inhibitor** | **N/A** | **51723** | **SwissProt** |
| P52193 | Calreticulin | CALR | 48039 | SwissProt |
| P52556 | Flavin reductase (NADPH) | BLVRB | 22132 | SwissProt |
| P52897 | Prostaglandin F synthase 2 | N/A | 36742 | SwissProt |
| P52898 | Dihydrodiol dehydrogenase 3 | N/A | 36784 | SwissProt |
| P53712 | Integrin beta-1 | ITGB1 | 88094 | SwissProt |
| P54149 | Mitochondrial peptide methionine sulfoxide reductase | MSRA | 25818 | SwissProt |
| P55906 | Transforming growth factor-beta-induced protein ig-h3 | TGFBI | 74408 | SwissProt |
| P56652 | Inter-alpha-trypsin inhibitor heavy chain H3 | ITIH3 | 99551 | SwissProt |
| P56701 | 26S proteasome non-ATPase regulatory subunit 2 | PSMD2 | 100258 | SwissProt |
| P60661 | Myosin light polypeptide 6 | MYL6 | 16930 | SwissProt |
| P60712 | Actin cytoplasmic 1 | ACTB | 41737 | SwissProt |
| P61157 | Actin-related protein 3 | ACTR3 | 47371 | SwissProt |
| P61223 | Ras-related protein Rap-1b | RAP1B | 20825 | SwissProt |
| P61284 | 60S ribosomal protein L12 | RPL12 | 17819 | SwissProt |
| P61286 | Polyadenylate-binding protein 1 | PABPC1 | 70671 | SwissProt |
| P61585 | Transforming protein RhoA | RHOA | 21768 | SwissProt |
| P62157 | Calmodulin | CALM | 16838 | SwissProt |
| P62261 | 14-3-3 protein epsilon | YWHAE | 29174 | SwissProt |
| P62739 | Actin aortic smooth muscle | ACTA2 | 42009 | SwissProt |

| | | | | |
|---|---|---|---|---|
| P62803 | Histone H4 | N/A | 11367 | SwissProt |
| P62808 | Histone H2B type 1 | N/A | 13906 | SwissProt |
| P62833 | Ras-related protein Rap-1A | RAP1A | 20987 | SwissProt |
| P62871 | Guanine nucleotide-binding protein G(I)/G(S)/G(T) subunit beta-1 | GNB1 | 37377 | SwissProt |
| P62935 | Peptidyl-prolyl cis-trans isomerase A | PPIA | 17869 | SwissProt |
| P62992 | Ubiquitin-40S ribosomal protein S27a | RPS27A | 17965 | SwissProt |
| P62998 | Ras-related C3 botulinum toxin substrate 1 | RAC1 | 21450 | SwissProt |
| P63048 | Ubiquitin-60S ribosomal protein L40 | UBA52 | 14728 | SwissProt |
| P63097 | Guanine nucleotide-binding protein G(i) subunit alpha-1 | GNAI1 | 40361 | SwissProt |
| P63103 | 14-3-3 protein zeta/delta | YWHAZ | 27745 | SwissProt |
| P63243 | Receptor of activated protein C kinase 1 | RACK1 | 35077 | SwissProt |
| P63258 | Actin cytoplasmic 2 | ACTG1 | 41793 | SwissProt |
| P67774 | Serine/threonine-protein phosphatase 2A catalytic subunit alpha isoform | PPP2CA | 35594 | SwissProt |
| P68103 | Elongation factor 1-alpha 1 | EEF1A1 | 50141 | SwissProt |
| P68138 | Actin alpha skeletal muscle | ACTA1 | 42051 | SwissProt |
| P68250 | 14-3-3 protein beta/alpha | YWHAB | 28081 | SwissProt |
| P68252 | 14-3-3 protein gamma | YWHAG | 28253 | SwissProt |
| P68401 | Platelet-activating factor acetylhydrolase IB subunit alpha2 | PAFAH1B2P68402 | 25569 | SwissProt |
| P68509 | 14-3-3 protein eta | YWHAH | 28212 | SwissProt |
| P79103 | 40S ribosomal protein S4 | RPS4 | 29598 | SwissProt |
| P79134 | Annexin A6 | ANXA6 | 75907 | SwissProt |
| P79334 | Glycogen phosphorylase muscle form | PYGM | 97293 | SwissProt |
| P80012 | von Willebrand factor (Fragment) | VWF | 102600 | SwissProt |
| P80109 | Phosphatidylinositol-glycan-specific phospholipase D | GPLD1 | 92602 | SwissProt |
| P81125 | Alpha-soluble NSF attachment protein | NAPA | 33225 | SwissProt |
| P81187 | Complement factor B | CFB | 85366 | SwissProt |
| P81287 | Annexin A5 | ANXA5 | 36089 | SwissProt |
| **P81644** | **Apolipoprotein A-II** | **APOA2** | **11202** | **SwissProt** |
| P81947 | Tubulin alpha-1B chain | N/A | 50152 | SwissProt |
| P81948 | Tubulin alpha-4A chain | TUBA4A | 49924 | SwissProt |
| P82943 | Regakine-1 | N/A | 10281 | SwissProt |
| P84080 | ADP-ribosylation factor 1 | ARF1 | 20697 | SwissProt |
| P84081 | ADP-ribosylation factor 2 | ARF2 | 20746 | SwissProt |

| | | | | |
|---|---|---|---|---|
| P98140 | Coagulation factor XII | F12 | 67160 | SwissProt |
| **Q03247** | **Apolipoprotein E** | **APOE** | **35980** | **SwissProt** |
| Q03763 | Desmoglein-1 | DSG1 | 112243 | SwissProt |
| Q04467 | Isocitrate dehydrogenase [NADP] mitochondrial | IDH2 | 50739 | SwissProt |
| **Q05443** | **Lumican** | **LUM** | **38756** | **SwissProt** |
| Q05718 | Insulin-like growth factor-binding protein 6 | IGFBP6 | 24967 | SwissProt |
| Q06805 | Tyrosine-protein kinase receptor Tie-1 | TIE1 | 124953 | SwissProt |
| Q08D91 | Keratin type II cytoskeletal 75 | KRT75 | 59036 | SwissProt |
| Q08DA1 | Sodium/potassium-transporting ATPase subunit alpha-1 | ATP1A1 | 112643 | SwissProt |
| Q08DN8 | Flotillin-1 | FLOT1 | 47353 | SwissProt |
| Q08DP0 | Phosphoglucomutase-1 | PGM1 | 61589 | SwissProt |
| Q08DQ2 | Glutamine--fructose-6-phosphate aminotransferase [isomerizing] 2 | GFPT2 | 77081 | SwissProt |
| Q08DX7 | Protein spinster homolog 1 | SPNS1 | 56520 | SwissProt |
| Q08E11 | Peptidyl-prolyl cis-trans isomerase C | PPIC | 22811 | SwissProt |
| Q08E20 | S-formylglutathione hydrolase | ESD | 31548 | SwissProt |
| Q0IIF7 | Ubiquitin carboxyl-terminal hydrolase 14 | USP14 | 56013 | SwissProt |
| Q0IIG5 | ATP-dependent 6-phosphofructokinase muscle type | PFKM | 85294 | SwissProt |
| Q0IIG7 | Ras-related protein Rab-5A | RAB5A | 23689 | SwissProt |
| Q0P594 | Serine/threonine-protein phosphatase 2A catalytic subunit beta isoform | PPP2CB | 35561 | SwissProt |
| Q0P5A6 | 26S proteasome non-ATPase regulatory subunit 5 | PSMD5 | 56043 | SwissProt |
| Q0P5F9 | 2-aminomuconic semialdehyde dehydrogenase | ALDH8A1 | 53363 | SwissProt |
| Q0P5J4 | Keratin type I cytoskeletal 25 | KRT25 | 49313 | SwissProt |
| Q0P5J6 | Keratin type I cytoskeletal 27 | KRT27 | 49907 | SwissProt |
| Q0V8B6 | Tripeptidyl-peptidase 1 | TPP1 | 61352 | SwissProt |
| Q0V8F1 | Coronin-7 | CORO7 | 99252 | SwissProt |
| Q0VCA8 | 3-hydroxyanthranilate 3 4-dioxygenase | HAAO | 32493 | SwissProt |
| Q0VCG9 | Pentraxin-related protein PTX3 | PTX3 | 42021 | SwissProt |
| Q0VCK0 | Bifunctional purine biosynthesis protein ATIC | ATIC | 64483 | SwissProt |
| Q0VCL3 | Ubiquitin-like-conjugating enzyme ATG3 | ATG3 | 35752 | SwissProt |
| Q0VCM4 | Glycogen phosphorylase liver form | PYGL | 97456 | SwissProt |
| **Q0VCM5** | **Inter-alpha-trypsin inhibitor heavy chain H1** | **ITIH1** | **101237** | **SwissProt** |

| | | | | |
|---|---|---|---|---|
| Q0VCN1 | NmrA-like family domain-containing protein 1 | NMRAL1 | 33154 | SwissProt |
| Q0VCP3 | Olfactomedin-like protein 3 | OLFML3 | 45886 | SwissProt |
| Q0VCU1 | Cytoplasmic aconitate hydratase | ACO1 | 98204 | SwissProt |
| Q0VCX1 | Complement C1s subcomponent | C1S | 76609 | SwissProt |
| Q0VCX2 | Endoplasmic reticulum chaperone BiP | HSPA5 | 72400 | SwissProt |
| Q0VCX4 | Catenin beta-1 | CTNNB1 | 85511 | SwissProt |
| Q0VD19 | Sphingomyelin phosphodiesterase | SMPD1 | 69392 | SwissProt |
| Q148C9 | Heme-binding protein 1 | HEBP1 | 21230 | SwissProt |
| Q148F1 | Cofilin-2 | CFL2 | 18737 | SwissProt |
| Q148H5 | Keratin type II cytoskeletal 71 | KRT71 | 57408 | SwissProt |
| **Q148H6** | **Keratin type I cytoskeletal 28** | **KRT28** | **50775** | **SwissProt** |
| Q148H7 | Keratin type II cytoskeletal 79 | KRT79 | 57721 | SwissProt |
| Q148J6 | Actin-related protein 2/3 complex subunit 4 | ARPC4 | 19667 | SwissProt |
| Q17QH6 | Collectin-11 | COLEC11 | 28333 | SwissProt |
| Q17R06 | Ras-related protein Rab-21 | RAB21 | 24146 | SwissProt |
| Q1JP75 | L-xylulose reductase | DCXR | 25650 | SwissProt |
| Q1JP79 | Actin-related protein 2/3 complex subunit 1A | ARPC1A | 41539 | SwissProt |
| Q1JPA6 | Arylamine N-acetyltransferase 1 | NAT1 | 34199 | SwissProt |
| Q1JPJ2 | Xaa-Pro aminopeptidase 1 | XPNPEP1 | 69790 | SwissProt |
| Q1LZA3 | Asparagine synthetase [glutamine-hydrolyzing] | ASNS | 64220 | SwissProt |
| Q1RMJ6 | Rho-related GTP-binding protein RhoC | RHOC | 22006 | SwissProt |
| Q1RMR4 | Ras-related protein Rab-15 | RAB15 | 24425 | SwissProt |
| Q1RMU3 | Prolyl 4-hydroxylase subunit alpha-1 | P4HA1 | 61010 | SwissProt |
| Q24K22 | Hepatocyte growth factor-like protein | MST1 | 79973 | SwissProt |
| Q27965 | Heat shock 70 kDa protein 1B | HSPA1B | 70229 | SwissProt |
| Q27967 | Secreted phosphoprotein 24 | SPP2 | 23134 | SwissProt |
| Q27975 | Heat shock 70 kDa protein 1A | HSPA1A | 70259 | SwissProt |
| Q27991 | Myosin-10 | MYH10 | 229097 | SwissProt |
| Q28007 | Dihydropyrimidine dehydrogenase [NADP(+)] | DPYD | 111697 | SwissProt |
| Q28017 | Platelet-activating factor acetylhydrolase | PLA2G7 | 50133 | SwissProt |
| Q28035 | Glutathione S-transferase A1 | GSTA1 | 25452 | SwissProt |
| **Q28065** | **C4b-binding protein alpha chain** | **C4BPA** | **68887** | **SwissProt** |
| **Q28085** | **Complement factor H** | **CFH** | **140374** | **SwissProt** |
| Q28106 | Contactin-1 | CNTN1 | 113385 | SwissProt |

| | | | | |
|---|---|---|---|---|
| **Q28107** | **Coagulation factor V** | **F5** | **248981** | **SwissProt** |
| Q28115 | Glial fibrillary acidic protein | GFAP | 49512 | SwissProt |
| Q28156 | cGMP-specific 3' 5'-cyclic phosphodiesterase | PDE5A | 98627 | SwissProt |
| Q28178 | Thrombospondin-1 | THBS1 | 129534 | SwissProt |
| Q28205 | Tubulin-specific chaperone D | TBCD | 133014 | SwissProt |
| Q28824 | Myosin light chain kinase smooth muscle | MYLK | 128825 | SwissProt |
| **Q29443** | **Serotransferrin** | **TF** | **77753** | **SwissProt** |
| Q29451 | Lysosomal alpha-mannosidase | MAN2B1 | 112919 | SwissProt |
| Q29460 | Platelet-activating factor acetylhydrolase IB subunit alpha1 | PAFAH1B3 | 25865 | SwissProt |
| **Q29RQ1** | **Complement component C7** | **C7** | **93090** | **SwissProt** |
| Q29RU2 | Oncoprotein-induced transcript 3 protein | OIT3 | 60550 | SwissProt |
| Q29RU4 | Complement component C6 | C6 | 104541 | SwissProt |
| Q29S21 | Keratin type II cytoskeletal 7 | KRT7 | 51578 | SwissProt |
| Q2HJ33 | Obg-like ATPase 1 | OLA1 | 44744 | SwissProt |
| Q2HJ49 | Moesin | MSN | 67975 | SwissProt |
| Q2HJ86 | Tubulin alpha-1D chain | TUBA1D | 50283 | SwissProt |
| Q2HJB8 | Tubulin alpha-8 chain | TUBA8 | 50054 | SwissProt |
| Q2HJF4 | Phenazine biosynthesis-like domain-containing protein | PBLD | 31883 | SwissProt |
| Q2HJG5 | Vacuolar protein sorting-associated protein 35 | VPS35 | 91751 | SwissProt |
| Q2HJH2 | Ras-related protein Rab-1B | RAB1B | 22202 | SwissProt |
| Q2HJH3 | Ester hydrolase C11orf54 homolog | N/A | 35155 | SwissProt |
| Q2HJI8 | Ras-related protein Rab-8B | RAB8B | 23656 | SwissProt |
| Q2KI42 | 26S proteasome non-ATPase regulatory subunit 11 | PSMD11 | 47464 | SwissProt |
| Q2KIA5 | Dynamin-1-like protein | DNM1L | 83352 | SwissProt |
| **Q2KIG3** | **Carboxypeptidase B2** | **CPB2** | **48822** | **SwissProt** |
| Q2KIH7 | Phenylalanine-4-hydroxylase | PAH | 51727 | SwissProt |
| Q2KIM0 | Tissue alpha-L-fucosidase | FUCA1 | 54089 | SwissProt |
| **Q2KIS7** | **Tetranectin** | **CLEC3B** | **22144** | **SwissProt** |
| **Q2KIT0** | **Protein HP-20 homolog** | **N/A** | **20646** | **SwissProt** |
| Q2KIU2 | Phosphomevalonate kinase | PMVK | 21956 | SwissProt |
| Q2KIW6 | 26S proteasome regulatory subunit 10B | PSMC6 | 44074 | SwissProt |
| Q2KJ25 | 26S proteasome non-ATPase regulatory subunit 12 | PSMD12 | 53053 | SwissProt |
| Q2KJ32 | Methanethiol oxidase | SELENBP1 | 52555 | SwissProt |
| Q2KJ33 | LIM and senescent cell antigen-like-containing domain protein 2 | LIMS2 | 38895 | SwissProt |
| Q2KJ46 | 26S proteasome non-ATPase regulatory subunit 3 | PSMD3 | 60957 | SwissProt |

| | | | | |
|---|---|---|---|---|
| Q2KJ63 | Plasma kallikrein | KLKB1 | 70994 | SwissProt |
| **Q2KJ83** | **Carboxypeptidase N catalytic chain** | **CPN1** | **52669** | **SwissProt** |
| Q2KJ93 | Cell division control protein 42 homolog | CDC42 | 21259 | SwissProt |
| Q2KJC6 | S-adenosylmethionine synthase isoform type-1 | MAT1A | 43761 | SwissProt |
| Q2KJD0 | Tubulin beta-5 chain | TUBB5 | 49671 | SwissProt |
| Q2KJD2 | Vesicle-associated membrane protein 3 | VAMP3 | 11535 | SwissProt |
| Q2KJE5 | Glyceraldehyde-3-phosphate dehydrogenase testis-specific | GAPDHS | 43288 | SwissProt |
| **Q2KJF1** | **Alpha-1B-glycoprotein** | **A1BG** | **53554** | **SwissProt** |
| Q2KJG3 | Asparagine--tRNA ligase cytoplasmic | NARS | 64399 | SwissProt |
| Q2KJH4 | WD repeat-containing protein 1 | WDR1 | 66258 | SwissProt |
| Q2KJH6 | Serpin H1 | SERPINH1 | 46507 | SwissProt |
| Q2KJH9 | 4-trimethylaminobutyraldehyde dehydrogenase | ALDH9A1 | 53977 | SwissProt |
| Q2M2T1 | Histone H2B type 1-K | H2BC12 | 13875 | SwissProt |
| Q2NKY7 | Septin-2 | SEPTIN2 | 41572 | SwissProt |
| Q2NKZ1 | T-complex protein 1 subunit eta | CCT7 | 59443 | SwissProt |
| Q2NL22 | Eukaryotic initiation factor 4A-III | EIF4A3 | 46841 | SwissProt |
| Q2NL31 | Methylthioribose-1-phosphate isomerase | MRI1 | 37826 | SwissProt |
| Q2T9X2 | T-complex protein 1 subunit delta | CCT4 | 58207 | SwissProt |
| Q2T9Y6 | Glutamate--cysteine ligase regulatory subunit | GCLM | 30533 | SwissProt |
| Q2TA29 | Ras-related protein Rab-11A | RAB11A | 24470 | SwissProt |
| Q2TA49 | Vasodilator-stimulated phosphoprotein | VASP | 40463 | SwissProt |
| Q2TBH7 | Ras-related protein Rab-4A | RAB4A | 24393 | SwissProt |
| Q2TBI0 | Lipopolysaccharide-binding protein | LBP | 53698 | SwissProt |
| Q2TBL6 | Transaldolase | TALDO1 | 37685 | SwissProt |
| Q2TBN5 | COMM domain-containing protein 9 | COMMD9 | 21781 | SwissProt |
| Q2TBQ3 | Guanidinoacetate N-methyltransferase | GAMT | 26610 | SwissProt |
| Q2TBQ8 | 6-phosphogluconolactonase | PGLS | 27547 | SwissProt |
| Q2TBU0 | Haptoglobin | HP | 44859 | SwissProt |
| Q2TBW7 | Sorting nexin-2 | SNX2 | 58454 | SwissProt |
| Q2TBX4 | Heat shock 70 kDa protein 13 | HSPA13 | 51921 | SwissProt |
| **Q2UVX4** | **Complement C3** | **C3** | **187252** | **SwissProt** |
| Q2YDE4 | Proteasome subunit alpha type-6 | PSMA6 | 27399 | SwissProt |

| | | | | |
|---|---|---|---|---|
| Q2YDG0 | G-protein coupled receptor family C group 5 member C | GPRC5C | 48459 | SwissProt |
| Q2YDJ9 | Mannose-1-phosphate guanyltransferase beta | GMPPB | 39837 | SwissProt |
| Q32KL2 | Proteasome subunit beta type-5 | PSMB5 | 28609 | SwissProt |
| Q32KN8 | Tubulin alpha-3 chain | TUBA3 | 49926 | SwissProt |
| Q32KY0 | Apolipoprotein D | APOD | 21402 | SwissProt |
| Q32L40 | T-complex protein 1 subunit alpha | TCP1 | 60207 | SwissProt |
| Q32L48 | Histone H2B type 1-N | H2BC15 | 13923 | SwissProt |
| Q32L99 | Prostaglandin reductase 2 | PTGR2 | 38400 | SwissProt |
| Q32LM0 | Endophilin-B1 | SH3GLB1 | 40777 | SwissProt |
| Q32LM2 | Small glutamine-rich tetratricopeptide repeat-containing protein alpha | SGTA | 34213 | SwissProt |
| Q32LP0 | Fermitin family homolog 3 | FERMT3 | 75783 | SwissProt |
| Q32LP2 | Radixin | RDX | 68568 | SwissProt |
| Q32PA4 | 14 kDa phosphohistidine phosphatase | PHPT1 | 13931 | SwissProt |
| Q32PF2 | ATP-citrate synthase | ACLY | 119789 | SwissProt |
| Q32PH8 | Elongation factor 1-alpha 2 | EEF1A2 | 50470 | SwissProt |
| Q32PI5 | Serine/threonine-protein phosphatase 2A 65 kDa regulatory subunit A alpha isoform | PPP2R1A | 65291 | SwissProt |
| **Q32PJ2** | **Apolipoprotein A-IV** | **APOA4** | **43018** | **SwissProt** |
| Q3B7M9 | Glycogen phosphorylase brain form | PYGB | 96340 | SwissProt |
| Q3MHH4 | Glutamine--tRNA ligase | QARS1 | 87643 | SwissProt |
| Q3MHL4 | Adenosylhomocysteinase | AHCY | 47638 | SwissProt |
| Q3MHL7 | T-complex protein 1 subunit zeta | CCT6A | 57956 | SwissProt |
| Q3MHM5 | Tubulin beta-4B chain | TUBB4B | 49831 | SwissProt |
| Q3MHN0 | Proteasome subunit beta type-6 | PSMB6 | 25542 | SwissProt |
| **Q3MHN2** | **Complement component C9** | **C9** | **61998** | **SwissProt** |
| **Q3MHN5** | **Vitamin D-binding protein** | **GC** | **53342** | **SwissProt** |
| Q3MHP2 | Ras-related protein Rab-11B | RAB11B | 24488 | SwissProt |
| Q3MHR0 | Acyl-protein thioesterase 1 | LYPLA1 | 24596 | SwissProt |
| Q3SWW8 | Thrombospondin-4 | THBS4 | 105974 | SwissProt |
| Q3SWX7 | Annexin A3 | ANXA3 | 36140 | SwissProt |
| Q3SWY2 | Integrin-linked protein kinase | ILK | 51447 | SwissProt |
| **Q3SX14** | **Gelsolin** | **GSN** | **80731** | **SwissProt** |
| Q3SYU2 | Elongation factor 2 | EEF2 | 95368 | SwissProt |
| Q3SYV4 | Adenylyl cyclase-associated protein 1 | CAP1 | 51273 | SwissProt |
| Q3SYZ4 | Aspartate--tRNA ligase cytoplasmic | DARS1 | 57036 | SwissProt |
| Q3SYZ6 | Xylulose kinase | XYLB | 53305 | SwissProt |

| | | | | |
|---|---|---|---|---|
| Q3SZ54 | Eukaryotic initiation factor 4A-I | EIF4A1 | 46154 | SwissProt |
| **Q3SZ57** | **Alpha-fetoprotein** | **AFP** | **68588** | **SwissProt** |
| Q3SZ62 | Phosphoglycerate mutase 1 | PGAM1 | 28852 | SwissProt |
| Q3SZ65 | Eukaryotic initiation factor 4A-II | EIF4A2 | 46402 | SwissProt |
| Q3SZB7 | Fructose-1 6-bisphosphatase 1 | FBP1 | 36728 | SwissProt |
| Q3SZD7 | Carbonyl reductase [NADPH] 1 | CBR1 | 30533 | SwissProt |
| Q3SZF2 | ADP-ribosylation factor 4 | ARF4 | 20528 | SwissProt |
| Q3SZH7 | Leukotriene A-4 hydrolase | LTA4H | 69306 | SwissProt |
| Q3SZI4 | 14-3-3 protein theta | YWHAQ | 27764 | SwissProt |
| Q3SZJ0 | Argininosuccinate lyase | ASL | 52743 | SwissProt |
| Q3SZJ4 | Prostaglandin reductase 1 | PTGR1 | 35706 | SwissProt |
| Q3SZJ9 | Phosphomannomutase 2 | PMM2 | 27997 | SwissProt |
| Q3SZK8 | Na(+)/H(+) exchange regulatory cofactor NHE-RF1 | SLC9A3R1 | 39603 | SwissProt |
| Q3SZP2 | Microtubule-associated protein RP/EB family member 2 | MAPRE2 | 36988 | SwissProt |
| Q3SZP7 | Villin-1 | VIL1 | 92802 | SwissProt |
| **Q3SZR3** | **Alpha-1-acid glycoprotein** | **ORM1** | **23182** | **SwissProt** |
| Q3SZV3 | Elongation factor 1-gamma | EEF1G | 50378 | SwissProt |
| **Q3SZV7** | **Hemopexin** | **HPX** | **52209** | **SwissProt** |
| Q3T004 | Serum amyloid P-component | APCS | 25183 | SwissProt |
| Q3T014 | Bisphosphoglycerate mutase | BPGM | 30061 | SwissProt |
| **Q3T052** | **Inter-alpha-trypsin inhibitor heavy chain H4** | **ITIH4** | **101513** | **SwissProt** |
| Q3T054 | GTP-binding nuclear protein Ran | RAN | 24423 | SwissProt |
| Q3T056 | L-lactate dehydrogenase A-like 6B | LDHAL6B | 41592 | SwissProt |
| Q3T0A3 | Complement factor D | CFD | 27878 | SwissProt |
| Q3T0C6 | Sodium/potassium-transporting ATPase subunit beta-3 | ATP1B3 | 31521 | SwissProt |
| Q3T0D0 | Heterogeneous nuclear ribonucleoprotein K | HNRNPK | 51019 | SwissProt |
| Q3T0F5 | Ras-related protein Rab-7a | RAB7A | 23544 | SwissProt |
| Q3T0L2 | Endoplasmic reticulum resident protein 44 | ERP44 | 46836 | SwissProt |
| Q3T0P6 | Phosphoglycerate kinase 1 | PGK1 | 44538 | SwissProt |
| Q3T0R1 | 40S ribosomal protein S18 | RPS18 | 17719 | SwissProt |
| Q3T0S5 | Fructose-bisphosphate aldolase B | ALDOB | 39543 | SwissProt |
| Q3T0T7 | GTP-binding protein SAR1b | SAR1B | 22409 | SwissProt |
| Q3T0V9 | Deoxyribose-phosphate aldolase | DERA | 35246 | SwissProt |
| Q3T0W4 | Protein phosphatase 1 regulatory subunit 7 | PPP1R7 | 41388 | SwissProt |

| | | | | |
|---|---|---|---|---|
| Q3T0X5 | Proteasome subunit alpha type-1 | PSMA1 | 29586 | SwissProt |
| Q3T0Y5 | Proteasome subunit alpha type-2 | PSMA2 | 25899 | SwissProt |
| Q3T0Z7 | Dihydropteridine reductase | QDPR | 25504 | SwissProt |
| Q3T145 | Malate dehydrogenase cytoplasmic | MDH1 | 36438 | SwissProt |
| Q3T149 | Heat shock protein beta-1 | HSPB1 | 22393 | SwissProt |
| **Q3Y5Z3** | **Adiponectin** | **ADIPOQ** | **26133** | **SwissProt** |
| Q3ZBA8 | Protein NDRG2 | NDRG2 | 39208 | SwissProt |
| **Q3ZBD7** | **Glucose-6-phosphate isomerase** | **GPI** | **62855** | **SwissProt** |
| Q3ZBF7 | Prostaglandin E synthase 3 | PTGES3 | 18697 | SwissProt |
| Q3ZBG0 | Proteasome subunit alpha type-7 | PSMA7 | 27869 | SwissProt |
| Q3ZBH0 | T-complex protein 1 subunit beta | CCT2 | 57475 | SwissProt |
| Q3ZBM5 | Sorting nexin-5 | SNX5 | 46819 | SwissProt |
| Q3ZBN5 | Asporin | ASPN | 42119 | SwissProt |
| Q3ZBT1 | Transitional endoplasmic reticulum ATPase | VCP | 89330 | SwissProt |
| Q3ZBT5 | Syntaxin-7 | STX7 | 29656 | SwissProt |
| Q3ZBU7 | Tubulin beta-4A chain | TUBB4A | 49586 | SwissProt |
| Q3ZBV8 | Threonine--tRNA ligase 1 cytoplasmic | TARS1 | 83492 | SwissProt |
| Q3ZBW5 | Rho-related GTP-binding protein RhoB | RHOB | 22123 | SwissProt |
| Q3ZC07 | Actin alpha cardiac muscle 1 | ACTC1 | 42019 | SwissProt |
| Q3ZC09 | Beta-enolase | ENO3 | 47096 | SwissProt |
| Q3ZC42 | Alcohol dehydrogenase class-3 | ADH5 | 39677 | SwissProt |
| Q3ZC84 | Cytosolic non-specific dipeptidase | CNDP2 | 52655 | SwissProt |
| Q3ZCD0 | CD81 antigen | CD81 | 25854 | SwissProt |
| Q3ZCI9 | T-complex protein 1 subunit theta | CCT8 | 59610 | SwissProt |
| Q3ZCJ2 | Aldo-keto reductase family 1 member A1 | AKR1A1 | 36617 | SwissProt |
| Q3ZCJ7 | Tubulin alpha-1C chain | TUBA1C | 49857 | SwissProt |
| Q3ZCJ8 | Dipeptidyl peptidase 1 | CTSC | 51949 | SwissProt |
| Q3ZCK9 | Proteasome subunit alpha type-4 | PSMA4 | 29484 | SwissProt |
| Q3ZEJ6 | Serpin A3-3 | SERPINA3-3 | 46326 | SwissProt |
| Q56JW4 | Adenine phosphoribosyltransferase | APRT | 19537 | SwissProt |
| Q56JZ9 | Glia maturation factor gamma | GMFG | 16770 | SwissProt |
| Q58CQ2 | Actin-related protein 2/3 complex subunit 1B | ARPC1B | 40976 | SwissProt |
| Q58CQ9 | Pantetheinase | VNN1 | 56947 | SwissProt |

| | | | | |
|---|---|---|---|---|
| Q58CS8 | N-acetylglucosamine-1-phosphotransferase subunit gamma | GNPTG | 33783 | SwissProt |
| Q58CY6 | Prostamide/prostaglandin F synthase | PRXL2B | 21497 | SwissProt |
| Q58D31 | Sorbitol dehydrogenase | SORD | 38099 | SwissProt |
| **Q58D62** | **Fetuin-B** | **FETUB** | **42663** | **SwissProt** |
| Q58D84 | Follistatin-related protein 1 | FSTL1 | 34856 | SwissProt |
| Q58DD0 | Angiotensin-converting enzyme 2 | ACE2 | 93067 | SwissProt |
| Q58DK4 | Triokinase/FMN cyclase | TKFC | 59131 | SwissProt |
| Q58DS9 | Ras-related protein Rab-5C | RAB5C | 23467 | SwissProt |
| Q59A32 | Trifunctional purine biosynthetic protein adenosine-3 | GART | 107907 | SwissProt |
| Q5E947 | Peroxiredoxin-1 | PRDX1 | 22210 | SwissProt |
| Q5E956 | Triosephosphate isomerase | TPI1 | 26690 | SwissProt |
| Q5E973 | 60S ribosomal protein L18 | RPL18 | 21535 | SwissProt |
| Q5E983 | Elongation factor 1-beta | EEF1B | 24805 | SwissProt |
| Q5E987 | Proteasome subunit alpha type-5 | PSMA5 | 26411 | SwissProt |
| Q5E997 | F-actin-capping protein subunit alpha-2 | CAPZA2 | 32979 | SwissProt |
| Q5E998 | Cathepsin L2 | CTSV | 37393 | SwissProt |
| Q5E9A3 | Poly(rC)-binding protein 1 | PCBP1 | 37498 | SwissProt |
| Q5E9B1 | L-lactate dehydrogenase B chain | LDHB | 36724 | SwissProt |
| Q5E9B5 | Actin gamma-enteric smooth muscle | ACTG2 | 41877 | SwissProt |
| Q5E9B7 | Chloride intracellular channel protein 1 | CLIC1 | 26992 | SwissProt |
| Q5E9D5 | Destrin | DSTN | 18506 | SwissProt |
| Q5E9E1 | PDZ and LIM domain protein 1 | PDLIM1 | 35881 | SwissProt |
| Q5E9F5 | Transgelin-2 | TAGLN2 | 22426 | SwissProt |
| Q5E9F7 | Cofilin-1 | CFL1 | 18519 | SwissProt |
| Q5E9F9 | 26S proteasome regulatory subunit 7 | PSMC2 | 48634 | SwissProt |
| Q5E9I6 | ADP-ribosylation factor 3 | ARF3 | 20601 | SwissProt |
| Q5E9P9 | Serine hydroxymethyltransferase cytosolic | SHMT1 | 52978 | SwissProt |
| Q5E9R3 | EH domain-containing protein 1 | EHD1 | 60682 | SwissProt |
| Q5E9S9 | Sodium-coupled neutral amino acid transporter 5 | SLC38A5 | 51958 | SwissProt |
| Q5E9Z2 | Hyaluronan-binding protein 2 | HABP2 | 62441 | SwissProt |
| Q5EA01 | Beta-1 4-glucuronyltransferase 1 | B4GAT1 | 47231 | SwissProt |
| Q5EA20 | 4-hydroxyphenylpyruvate dioxygenase | HPD | 44963 | SwissProt |

| | | | | |
|---|---|---|---|---|
| Q5EA61 | Creatine kinase B-type | CKB | 42719 | SwissProt |
| Q5EA79 | Galactose mutarotase | GALM | 37614 | SwissProt |
| Q5EA88 | Glycerol-3-phosphate dehydrogenase [NAD(+)] cytoplasmic | GPD1 | 37648 | SwissProt |
| Q5EAD2 | D-3-phosphoglycerate dehydrogenase | PHGDH | 56452 | SwissProt |
| Q5I597 | Betaine--homocysteine S-methyltransferase 1 | BHMT | 44878 | SwissProt |
| Q5NTB3 | Coagulation factor XI | F11 | 69872 | SwissProt |
| **Q5XQN5** | **Keratin type II cytoskeletal 5** | **KRT5** | **62937** | **SwissProt** |
| Q6B856 | Tubulin beta-2B chain | TUBB2B | 49953 | SwissProt |
| Q6EWQ7 | Eukaryotic translation initiation factor 5A-1 | EIF5A | 16832 | SwissProt |
| Q6Q137 | Septin-7 | SEPTIN7 | 50680 | SwissProt |
| Q6URK6 | Cadherin-5 | CDH5 | 87467 | SwissProt |
| Q71SP7 | Fatty acid synthase | FASN | 274552 | SwissProt |
| Q76LV1 | Heat shock protein HSP 90-beta | HSP90AB1 | 83253 | SwissProt |
| Q76LV2 | Heat shock protein HSP 90-alpha | HSP90AA1 | 84731 | SwissProt |
| Q7SIB2 | Collagen alpha-1(IV) chain | COL4A1 | 160412 | SwissProt |
| Q7SIB3 | Collagen alpha-2(IV) chain (Fragment) | COL4A2 | 25061 | SwissProt |
| Q7SIH1 | Alpha-2-macroglobulin | A2M | 167575 | SwissProt |
| Q8MJ50 | Osteoclast-stimulating factor 1 | OSTF1 | 23842 | SwissProt |
| Q8SPJ1 | Junction plakoglobin | JUP | 81821 | SwissProt |
| Q8SPP7 | Peptidoglycan recognition protein 1 | PGLYRP1 | 21063 | SwissProt |
| Q8SQA4 | Adhesion G protein-coupled receptor E5 | ADGRE5 | 80322 | SwissProt |
| Q92176 | Coronin-1A | CORO1A | 50979 | SwissProt |
| Q95114 | Lactadherin | MFGE8 | 47411 | SwissProt |
| Q95115 | Signal transducer and activator of transcription 5A | STAT5A | 90700 | SwissProt |
| **Q95121** | **Pigment epithelium-derived factor** | **SERPINF1** | **46229** | **SwissProt** |
| Q95122 | Monocyte differentiation antigen CD14 | CD14 | 39667 | SwissProt |
| Q95JC7 | Neutral amino acid transporter B(0) | SLC1A5 | 56448 | SwissProt |
| **Q95M17** | **Acidic mammalian chitinase** | **CHIA** | **52129** | **SwissProt** |
| Q95M18 | Endoplasmin | HSP90B1 | 92427 | SwissProt |
| Q9BGI2 | Peroxiredoxin-4 | PRDX4 | 30741 | SwissProt |
| Q9BGI3 | Peroxiredoxin-2 | PRDX2 | 21946 | SwissProt |
| Q9GKN7 | Ammonium transporter Rh type A | RHAG | 46719 | SwissProt |
| Q9GMB8 | Serine--tRNA ligase cytoplasmic | SARS1 | 58605 | SwissProt |

| | | | | |
|---|---|---|---|---|
| Q9N0V4 | Glutathione S-transferase Mu 1 | GSTM1 | 25635 | SwissProt |
| Q9N179 | Protein 4.1 | EPB41 | 69258 | SwissProt |
| **Q9N2I2** | **Plasma serine protease inhibitor** | **SERPINA5** | **45297** | **SwissProt** |
| Q9TRY0 | Peptidyl-prolyl cis-trans isomerase FKBP4 | FKBP4 | 51529 | SwissProt |
| **Q9TT36** | **Thyroxine-binding globulin** | **SERPINA7** | **46017** | **SwissProt** |
| **Q9TTE1** | **Serpin A3-1** | **SERPINA3-1** | **46237** | **SwissProt** |
| Q9TTJ5 | Regucalcin | RGN | 33308 | SwissProt |
| Q9TU25 | Ras-related C3 botulinum toxin substrate 2 | RAC2 | 21424 | SwissProt |
| Q9TUM3 | Signal transducer and activator of transcription 5B | STAT5B | 89991 | SwissProt |
| Q9XSA7 | Chloride intracellular channel protein 4 | CLIC4 | 28727 | SwissProt |
| Q9XSC6 | Creatine kinase M-type | CKM | 42989 | SwissProt |
| Q9XSG3 | Isocitrate dehydrogenase [NADP] cytoplasmic | IDH1 | 46785 | SwissProt |
| Q9XSJ4 | Alpha-enolase | ENO1 | 47326 | SwissProt |
| Q9XTA2 | Prolyl endopeptidase | PREP | 80641 | SwissProt |
| A0A0A0MP92 | Serpin A3-7 | SERPINA3-7 | 47016 | TrEMBL |
| A0A0A8J2N9 | Amino acid transporter | ASCT2 | 56448 | TrEMBL |
| A0A0F6QMJ3 | Complement component 5 | C5 | 188809 | TrEMBL |
| A0A0F6QNP7 | Complement C3 | C3 | 187181 | TrEMBL |
| A0A0M4MD57 | Haptoglobin | HP | 44859 | TrEMBL |
| A0A140T843 | Beta-2-glycoprotein 1 | APOH | 38245 | TrEMBL |
| A0A140T851 | Vitamin K-dependent protein C | PROC | 50709 | TrEMBL |
| A0A140T872 | Dihydrofolate reductase | DHFR | 21603 | TrEMBL |
| A0A140T874 | 2-aminomuconic semialdehyde dehydrogenase | ALDH8A1 | 53349 | TrEMBL |
| A0A140T881 | Apolipoprotein E | APOE | 37424 | TrEMBL |
| A0A140T887 | Uridine 5'-monophosphate synthase | UMPS | 52282 | TrEMBL |
| A0A140T891 | Pantetheinase | VNN1 | 56980 | TrEMBL |
| A0A140T894 | 14-3-3 protein beta/alpha | YWHAB | 28082 | TrEMBL |
| A0A140T896 | Serine hydroxymethyltransferase | SHMT1 | 52969 | TrEMBL |
| A0A140T897 | Albumin | ALB | 69324 | TrEMBL |
| A0A140T8A5 | Isocitrate dehydrogenase [NADP] | IDH1 | 46759 | TrEMBL |
| A0A140T8C8 | Kininogen-1 | KNG1 | 68965 | TrEMBL |
| A0A1B0Z542 | Heat shock 27 kDa protein 1 | HspB1 | 22359 | TrEMBL |
| A0A1C9EIX3 | Heat shock protein beta-1 | HSPB1 | 22365 | TrEMBL |
| A0A1C9EIX6 | Heat shock protein beta-1 | HSPB1 | 22403 | TrEMBL |
| A0A1K0FUD3 | Globin C1 | GLNC1 | 15184 | TrEMBL |
| A0A2Q9 | Alanine--tRNA ligase (Fragment) | N/A | 96196 | TrEMBL |
| A0A3B0J3V0 | Adiponectin D | ADID | 26133 | TrEMBL |

| | | | | |
|---|---|---|---|---|
| A0A3Q1LFG8 | F-actin-capping protein subunit alpha | CAPZA1 | 33821 | TrEMBL |
| A0A3Q1LFR2 | Glutathione transferase | GSTM4 | 25656 | TrEMBL |
| A0A3Q1LFR7 | Amino acid transporter | SLC1A5 | 55709 | TrEMBL |
| A0A3Q1LGM4 | Complement factor I | CFI | 64667 | TrEMBL |
| A0A3Q1LI40 | Complement component C7 | C7 | 92990 | TrEMBL |
| A0A3Q1LJG6 | Aldehyde oxidase | AOX1 | 147222 | TrEMBL |
| A0A3Q1LJU5 | EMAP like 2 | EML2 | 98456 | TrEMBL |
| A0A3Q1LJZ4 | FAT atypical cadherin 1 | FAT1 | 511840 | TrEMBL |
| A0A3Q1LK49 | Inter-alpha-trypsin inhibitor heavy chain H2 | ITIH2 | 96775 | TrEMBL |
| A0A3Q1LKB9 | Carboxypeptidase B2 | CPB2 | 44247 | TrEMBL |
| A0A3Q1LKF1 | Catalase | CAT | 59980 | TrEMBL |
| A0A3Q1LKF4 | Serine hydroxymethyltransferase | SHMT1 | 51508 | TrEMBL |
| A0A3Q1LKL7 | Adhesion G protein-coupled receptor E5 | ADGRE5 | 85238 | TrEMBL |
| A0A3Q1LKR8 | E1 ubiquitin-activating enzyme | UBA1 | 129453 | TrEMBL |
| A0A3Q1LLU1 | von Willebrand factor | VWF | 307986 | TrEMBL |
| A0A3Q1LM32 | Complement component C6 | C6 | 105116 | TrEMBL |
| A0A3Q1LMS5 | Heat shock cognate 71 kDa protein | HSPA8 | 71764 | TrEMBL |
| A0A3Q1LMT0 | Phosphopyruvate hydratase | ENO3 | 44851 | TrEMBL |
| A0A3Q1LMV5 | Glutathione S-transferase | GSTP1 | 26746 | TrEMBL |
| A0A3Q1LMZ4 | Neogenin 1 | NEO1 | 146767 | TrEMBL |
| A0A3Q1LP81 | Dihydropyrimidine dehydrogenase [NADP(+)] | DPYD | 105234 | TrEMBL |
| A0A3Q1LPF0 | Apolipoprotein E | APOE | 36039 | TrEMBL |
| A0A3Q1LPT9 | STEAP3 metalloreductase | STEAP3 | 60091 | TrEMBL |
| A0A3Q1LPX4 | Hydroxymethylglutaryl-CoA synthase | HMGCS1 | 57472 | TrEMBL |
| A0A3Q1LQ21 | Inter-alpha-trypsin inhibitor heavy chain H3 | ITIH3 | 99156 | TrEMBL |
| A0A3Q1LQI3 | Plexin B2 | PLXNB2 | 220480 | TrEMBL |
| A0A3Q1LQY7 | STEAP3 metalloreductase | STEAP3 | 54077 | TrEMBL |
| A0A3Q1LRD1 | Phosphoglucomutase-1 | PGM1 | 61575 | TrEMBL |
| A0A3Q1LRL7 | Xylulose kinase | XYLB | 58262 | TrEMBL |
| A0A3Q1LS55 | Alpha-1-acid glycoprotein | ORM1 | 21441 | TrEMBL |
| A0A3Q1LS65 | Homogentisate 1 2-dioxygenase | HGD | 49821 | TrEMBL |
| A0A3Q1LS74 | Complement factor H | CFH | 140330 | TrEMBL |
| A0A3Q1LSB6 | ATP-citrate synthase | ACLY | 120866 | TrEMBL |
| A0A3Q1LSP4 | Complement C8 alpha chain | C8A | 65499 | TrEMBL |
| A0A3Q1LT55 | Coagulation factor XI | F11 | 71641 | TrEMBL |
| A0A3Q1LTB9 | Coagulation factor XIII A chain | F13A1 | 75598 | TrEMBL |
| A0A3Q1LTF3 | Histidine ammonia-lyase | HAL | 68616 | TrEMBL |
| A0A3Q1LTP0 | Adenosine kinase | ADK | 38933 | TrEMBL |
| A0A3Q1LU13 | Dihydropteridine reductase | QDPR | 23756 | TrEMBL |

| | | | | |
|---|---|---|---|---|
| A0A3Q1LUD3 | Dihydropyrimidine dehydrogenase [NADP(+)] | DPYD | 109110 | TrEMBL |
| A0A3Q1LUD9 | Uncharacterized protein | DIAPH1 | 140882 | TrEMBL |
| A0A3Q1LUE9 | Ig-like domain-containing protein | N/A | 16709 | TrEMBL |
| A0A3Q1LUJ8 | Sodium/potassium-transporting ATPase subunit alpha | ATP1A1 | 115726 | TrEMBL |
| A0A3Q1LUY6 | GDP-mannose pyrophosphorylase A | GMPPA | 46519 | TrEMBL |
| A0A3Q1LVC7 | Ezrin | EZR | 78752 | TrEMBL |
| A0A3Q1LVC8 | Xaa-Pro aminopeptidase 1 | XPNPEP1 | 70895 | TrEMBL |
| A0A3Q1LVM5 | Inter-alpha-trypsin inhibitor heavy chain H1 | ITIH1 | 100465 | TrEMBL |
| A0A3Q1LVU1 | Ferritin | FTH1 | 24678 | TrEMBL |
| A0A3Q1LVV7 | Fibrinogen alpha chain | FGA | 89852 | TrEMBL |
| A0A3Q1LW12 | Phosphatidylinositol-glycan-specific phospholipase D | GPLD1 | 93001 | TrEMBL |
| A0A3Q1LW84 | Serine/threonine-protein phosphatase 2A 65 kDa regulatory subunit A alpha isoform | PPP2R1A | 64855 | TrEMBL |
| A0A3Q1LW96 | Inter-alpha-trypsin inhibitor heavy chain H1 | ITIH1 | 100823 | TrEMBL |
| A0A3Q1LWV8 | Ig-like domain-containing protein | N/A | 16434 | TrEMBL |
| A0A3Q1LX69 | L-xylulose reductase | DCXR | 24720 | TrEMBL |
| A0A3Q1LXH8 | Urocanate hydratase | UROC1 | 80328 | TrEMBL |
| A0A3Q1LXM2 | Collagen type XV alpha 1 chain | COL15A1 | 141554 | TrEMBL |
| A0A3Q1LXP4 | Beta-2-glycoprotein 1 | APOH | 37438 | TrEMBL |
| A0A3Q1LXR2 | Ras-related C3 botulinum toxin substrate 1 | RAC1 | 23491 | TrEMBL |
| A0A3Q1LY19 | L-lactate dehydrogenase | N/A | 36583 | TrEMBL |
| A0A3Q1LYE7 | RAB1A member RAS onco family | RAB1A | 21851 | TrEMBL |
| A0A3Q1LYV8 | LIM and senescent cell antigen-like-containing domain protein | LIMS1 | 39825 | TrEMBL |
| A0A3Q1LZS5 | Tetranectin | CLEC3B | 22293 | TrEMBL |
| A0A3Q1LZU8 | Glucosidase II alpha subunit | GANAB | 106921 | TrEMBL |
| A0A3Q1M0L3 | RNA helicase | EIF4A2 | 41961 | TrEMBL |
| A0A3Q1M0L5 | Uncharacterized protein | N/A | 22869 | TrEMBL |
| A0A3Q1M0N0 | Collagen type XV alpha 1 chain | COL15A1 | 142692 | TrEMBL |
| A0A3Q1M0R0 | Adhesion G protein-coupled receptor E5 | ADGRE5 | 90693 | TrEMBL |
| A0A3Q1M0U5 | Receptor protein-tyrosine kinase | TIE1 | 125367 | TrEMBL |
| A0A3Q1M0V5 | Phosphopyruvate hydratase | ENO3 | 62126 | TrEMBL |
| A0A3Q1M0W4 | Transforming growth factor beta receptor 3 | TGFBR3 | 93127 | TrEMBL |

| | | | | |
|---|---|---|---|---|
| A0A3Q1M119 | 14-3-3 protein epsilon | YWHAE | 27419 | TrEMBL |
| A0A3Q1M168 | Alpha-1 4 glucan phosphorylase | PYGL | 98157 | TrEMBL |
| A0A3Q1M1U2 | Serine/threonine-protein phosphatase 2A 65 kDa regulatory subunit A alpha isoform | PPP2R1A | 64172 | TrEMBL |
| A0A3Q1M1Z2 | Tubulin alpha chain | TUBA4A | 46450 | TrEMBL |
| A0A3Q1M2A8 | Complement factor I | CFI | 64923 | TrEMBL |
| A0A3Q1M2H4 | Aldehyde dehydrogenase family 16 member A1 | ALDH16A1 | 79615 | TrEMBL |
| A0A3Q1M3A4 | Uncharacterized protein | LOC528040 | 190321 | TrEMBL |
| A0A3Q1M3K7 | Ras-related protein Rab-7a | RAB7A | 26385 | TrEMBL |
| A0A3Q1M3N0 | Apolipoprotein A-IV | APOA4 | 43078 | TrEMBL |
| A0A3Q1M478 | C-1-tetrahydrofolate synthase cytoplasmic | MTHFD1 | 101522 | TrEMBL |
| A0A3Q1M4K3 | Ubiquitin-like domain-containing protein | LOC101902760 | 8634 | TrEMBL |
| A0A3Q1M4L0 | HipN domain-containing protein | N/A | 22849 | TrEMBL |
| A0A3Q1M4P8 | Collagen type XV alpha 1 chain | COL15A1 | 141251 | TrEMBL |
| A0A3Q1M558 | Actin alpha cardiac muscle 1 | ACTC1 | 43569 | TrEMBL |
| A0A3Q1M559 | Histidine ammonia-lyase | HAL | 76058 | TrEMBL |
| A0A3Q1M564 | Adiponectin | ADIPOQ | 35094 | TrEMBL |
| A0A3Q1M5H7 | Mannose receptor C type 2 | MRC2 | 167227 | TrEMBL |
| A0A3Q1M5Q6 | Extracellular matrix protein 1 | ECM1 | 63177 | TrEMBL |
| A0A3Q1M5R4 | L-lactate dehydrogenase | LDHB | 37471 | TrEMBL |
| A0A3Q1M5W5 | FAT atypical cadherin 1 | FAT1 | 507127 | TrEMBL |
| A0A3Q1M616 | Vitamin K-dependent protein C | PROC | 51837 | TrEMBL |
| A0A3Q1M6H3 | Alpha-soluble NSF attachment protein | NAPA | 36297 | TrEMBL |
| A0A3Q1M6N7 | Leukotriene A(4) hydrolase | LTA4H | 67283 | TrEMBL |
| A0A3Q1M6P2 | Thyroglobulin | TG | 303305 | TrEMBL |
| A0A3Q1M7B6 | Parvin beta | PARVB | 45803 | TrEMBL |
| A0A3Q1M7H6 | Contactin-1 | CNTN1 | 112715 | TrEMBL |
| A0A3Q1M7W9 | Dynamin-1-like protein | DNM1L | 78918 | TrEMBL |
| A0A3Q1M7Y3 | Ras-related C3 botulinum toxin substrate 1 | RAC1 | 30473 | TrEMBL |
| A0A3Q1M857 | PKS_ER domain-containing protein | N/A | 40138 | TrEMBL |
| A0A3Q1M8Y3 | FAT atypical cadherin 1 | FAT1 | 486621 | TrEMBL |
| A0A3Q1M913 | Phosphoribosylformylglycinamidine synthase | PFAS | 144247 | TrEMBL |
| A0A3Q1M944 | Collectin-11 | COLEC11 | 28550 | TrEMBL |
| A0A3Q1M9B3 | 60S ribosomal protein L18 | RPL18 | 23008 | TrEMBL |
| A0A3Q1M9D2 | Sulfurtransferase | MPST | 62972 | TrEMBL |
| A0A3Q1M9Q8 | ADP-ribosylation factor | N/A | 20733 | TrEMBL |

| | | | | |
|---|---|---|---|---|
| A0A3Q1MA31 | Inter-alpha-trypsin inhibitor heavy chain H4 | ITIH4 | 99646 | TrEMBL |
| A0A3Q1MA45 | Uncharacterized protein | N/A | 46719 | TrEMBL |
| A0A3Q1MAU7 | S-formylglutathione hydrolase | ESD | 32740 | TrEMBL |
| A0A3Q1MAY2 | Histidine ammonia-lyase | HAL | 71258 | TrEMBL |
| A0A3Q1MB39 | Adenosine kinase | ADK | 40403 | TrEMBL |
| A0A3Q1MB70 | Apolipoprotein M | APOM | 25020 | TrEMBL |
| A0A3Q1MBF0 | Neogenin 1 | NEO1 | 158594 | TrEMBL |
| A0A3Q1MBY4 | cAMP-dependent protein kinase type I-alpha regulatory subunit | PRKAR1A | 42491 | TrEMBL |
| A0A3Q1MC58 | Dihydropyrimidine dehydrogenase [NADP(+)] | DPYD | 109126 | TrEMBL |
| A0A3Q1MCW2 | Extracellular matrix protein 1 | ECM1 | 60395 | TrEMBL |
| A0A3Q1MDM5 | Sodium/potassium-transporting ATPase subunit alpha | ATP1A1 | 109830 | TrEMBL |
| A0A3Q1MF14 | Complement factor I | CFI | 68905 | TrEMBL |
| A0A3Q1MF21 | Periostin | POSTN | 89849 | TrEMBL |
| A0A3Q1MFI5 | Dihydropyrimidinase-related protein 2 | DPYSL2 | 73535 | TrEMBL |
| A0A3Q1MFJ2 | Uncharacterized protein | LOC112441469 | 24693 | TrEMBL |
| A0A3Q1MFJ3 | Methanethiol oxidase | SELENBP1 | 49252 | TrEMBL |
| A0A3Q1MFP4 | Neogenin 1 | NEO1 | 154044 | TrEMBL |
| A0A3Q1MFY4 | Sodium/potassium-transporting ATPase subunit alpha | ATP1A1 | 112713 | TrEMBL |
| A0A3Q1MG04 | Fibrinogen beta chain | FGB | 57467 | TrEMBL |
| A0A3Q1MG08 | MBL associated serine protease 1 | MASP1 | 79100 | TrEMBL |
| A0A3Q1MGI8 | Ribonucleoside-diphosphate reductase | RRM1 | 89720 | TrEMBL |
| A0A3Q1MGL5 | Uncharacterized protein | N/A | 35989 | TrEMBL |
| A0A3Q1MHR3 | Exostosin-2 | EXT2 | 84578 | TrEMBL |
| A0A3Q1MHX8 | RAB2A member RAS onco family | RAB2A | 22470 | TrEMBL |
| A0A3Q1MIA4 | 26S proteasome non-ATPase regulatory subunit 2 | PSMD2 | 99646 | TrEMBL |
| A0A3Q1MIB0 | IQ motif containing GTPase activating protein 1 | IQGAP1 | 180837 | TrEMBL |
| A0A3Q1MIF4 | Complement factor I | CFI | 67044 | TrEMBL |
| A0A3Q1MIS1 | Gelsolin | GSN | 90140 | TrEMBL |
| A0A3Q1MIW0 | Alpha-fetoprotein | AFP | 67862 | TrEMBL |
| A0A3Q1MIW2 | Gelsolin | GSN | 81946 | TrEMBL |
| A0A3Q1MJQ0 | Solute carrier family 29 member 1 | SLC29A1 | 54397 | TrEMBL |
| A0A3Q1MJT2 | Alpha-1B-glycoprotein | A1BG | 61960 | TrEMBL |
| A0A3Q1MKR2 | Glutamyl-prolyl-tRNA synthetase 1 | EPRS1 | 170743 | TrEMBL |
| A0A3Q1ML32 | Carbonyl reductase (NADPH) | CBR1 | 30970 | TrEMBL |

| | | | | |
|---|---|---|---|---|
| A0A3Q1MLQ5 | Solute carrier family 2 facilitated glucose transporter member 1 | SLC2A1 | 53903 | TrEMBL |
| A0A3Q1MLQ7 | Talin 1 | TLN1 | 270746 | TrEMBL |
| A0A3Q1MLX2 | Regucalcin | RGN | 33540 | TrEMBL |
| A0A3Q1MM52 | Argininosuccinate synthase | ASS1 | 47954 | TrEMBL |
| A0A3Q1MM55 | Glucose-6-phosphate 1-dehydrogenase | G6PD | 63252 | TrEMBL |
| A0A3Q1MN33 | Glutamate--cysteine ligase | GCLC | 72739 | TrEMBL |
| A0A3Q1MNN2 | Aldo_ket_red domain-containing protein | AKR7A2 | 39978 | TrEMBL |
| A0A3Q1MNN6 | Uncharacterized protein | LOC506828 | 159845 | TrEMBL |
| A0A3Q1MNW6 | Uncharacterized protein | N/A | 36283 | TrEMBL |
| A0A3Q1MP36 | Sulfotransferase | SULT1C4 | 35934 | TrEMBL |
| A0A3Q1MP40 | Complement component C7 | C7 | 90093 | TrEMBL |
| A0A3Q1MP45 | Complement factor H | CFH | 140969 | TrEMBL |
| A0A3Q1MP67 | 14-3-3 protein epsilon | YWHAE | 30107 | TrEMBL |
| A0A3Q1MP70 | Glucose-6-phosphate isomerase | GPI | 63636 | TrEMBL |
| A0A3Q1MPB3 | Fatty acid synthase | FASN | 269854 | TrEMBL |
| A0A3Q1MPF1 | Rab GDP dissociation inhibitor | GDI2 | 51069 | TrEMBL |
| A0A3Q1MPG8 | 1 4-alpha-glucan branching enzyme | GBE1 | 81005 | TrEMBL |
| A0A3Q1MQ59 | Solute carrier family 3 member 2 | SLC3A2 | 61994 | TrEMBL |
| A0A3Q1MQ68 | Dihydropyrimidinase-related protein 2 | DPYSL2 | 58147 | TrEMBL |
| A0A3Q1MQ74 | Plexin B2 | PLXNB2 | 204950 | TrEMBL |
| A0A3Q1MQM3 | ST13 Hsp70 interacting protein | ST13 | 42371 | TrEMBL |
| A0A3Q1MQS3 | 14-3-3 protein theta | YWHAQ | 29054 | TrEMBL |
| A0A3Q1MQV3 | Thrombospondin-1 | THBS1 | 129878 | TrEMBL |
| A0A3Q1MQZ2 | ATP-citrate synthase | ACLY | 119747 | TrEMBL |
| A0A3Q1MQZ5 | Phosphatidylinositol-glycan-specific phospholipase D | GPLD1 | 84709 | TrEMBL |
| A0A3Q1MR36 | Anion exchange protein | SLC4A1 | 101001 | TrEMBL |
| A0A3Q1MRK8 | Phosphodiesterase | PDE5A | 94788 | TrEMBL |
| A0A3Q1MRL0 | PKS_ER domain-containing protein | N/A | 40227 | TrEMBL |
| A0A3Q1MSP0 | Glutamyl-prolyl-tRNA synthetase 1 | EPRS1 | 162402 | TrEMBL |
| A0A3Q1MTI5 | Epoxide hydrolase 2 | EPHX2 | 62041 | TrEMBL |
| A0A3Q1MU33 | Annexin | ANXA5 | 35876 | TrEMBL |
| A0A3Q1MUA3 | Collagen type XV alpha 1 chain | COL15A1 | 137512 | TrEMBL |
| A0A3Q1MUJ8 | Plexin B2 | PLXNB2 | 228356 | TrEMBL |
| A0A3Q1MWA3 | Ras-related protein Rab-1B | RAB1B | 32301 | TrEMBL |
| A0A3Q1MZJ9 | Apolipoprotein A-IV | APOA4 | 45639 | TrEMBL |
| A0A3Q1N147 | RAB1A member RAS onco family | RAB1A | 24625 | TrEMBL |

| | | | | |
|---|---|---|---|---|
| A0A3Q1N1A7 | Carboxypeptidase B2 | CPB2 | 44774 | TrEMBL |
| A0A3Q1N1A8 | Uncharacterized protein | N/A | 38228 | TrEMBL |
| A0A3Q1N1T6 | Glucosidase II alpha subunit | GANAB | 109162 | TrEMBL |
| A0A3Q1N308 | Methanethiol oxidase | SELENBP1 | 53035 | TrEMBL |
| A0A3Q1N461 | Complement factor I | CFI | 65979 | TrEMBL |
| A0A3Q1N4H3 | Sushi domain-containing protein | N/A | 26810 | TrEMBL |
| A0A3Q1N521 | Neogenin 1 | NEO1 | 150301 | TrEMBL |
| A0A3Q1N5X6 | FAM20C golgi associated secretory pathway kinase | FAM20C | 80522 | TrEMBL |
| A0A3Q1N764 | Collectin-11 | COLEC11 | 26058 | TrEMBL |
| A0A3Q1N7G4 | Homogentisate 1 2-dioxygenase | HGD | 49525 | TrEMBL |
| A0A3Q1N894 | F-actin-capping protein subunit alpha | CAPZA1 | 33174 | TrEMBL |
| A0A3Q1N8E3 | Ras-related protein Rab-5C | RAB5C | 24595 | TrEMBL |
| A0A3Q1N9Y5 | Sulfhydryl oxidase | QSOX1 | 86164 | TrEMBL |
| A0A3Q1NCY0 | S-adenosylmethionine synthase | MAT1A | 41938 | TrEMBL |
| A0A3Q1NEF8 | Ras-related C3 botulinum toxin substrate 1 | RAC1 | 28432 | TrEMBL |
| A0A3Q1NGI1 | Xylulose kinase | XYLB | 55470 | TrEMBL |
| A0A3Q1NJ03 | Serpin family A member 10 | SERPINA10 | 47190 | TrEMBL |
| A0A3Q1NJ70 | ATP-dependent 6-phosphofructokinase | PFKP | 97498 | TrEMBL |
| A0A3Q1NJB1 | Ceruloplasmin | CP | 120779 | TrEMBL |
| A0A3Q1NJR8 | Antithrombin-III | SERPINC1 | 60069 | TrEMBL |
| A0A3Q1NM28 | Hyaluronoglucosaminidase | CEMIP | 146012 | TrEMBL |
| A0A3Q1NNJ3 | Flavin reductase (NADPH) | BLVRB | 20511 | TrEMBL |
| A0A3Q1NNK1 | Periostin | POSTN | 90023 | TrEMBL |
| A0A3Q1NNN2 | Solute carrier family 7 member 5 | SLC7A5 | 65697 | TrEMBL |
| A0A3S5ZPA0 | Exostosin-2 | EXT2 | 89752 | TrEMBL |
| A0A3S5ZPB5 | Extracellular matrix protein 1 | ECM1 | 58384 | TrEMBL |
| A0A3S5ZPD5 | Glutathione transferase | GSTT2 | 31633 | TrEMBL |
| A0A3S5ZPM3 | 6-phosphogluconate dehydrogenase decarboxylating | PGD | 60780 | TrEMBL |
| A0A452DHZ3 | Plasma serine protease inhibitor | SERPINA5 | 43568 | TrEMBL |
| A0A452DI24 | Rab GDP dissociation inhibitor | GDI2 | 48136 | TrEMBL |
| A0A452DI25 | Hemopexin | HPX | 52166 | TrEMBL |
| A0A452DI31 | Phosphopyruvate hydratase | ENO3 | 48310 | TrEMBL |
| A0A452DI66 | Prothrombin | F2 | 70575 | TrEMBL |
| A0A452DIB6 | Actin-related protein 2/3 complex subunit 4 | ARPC4 | 20893 | TrEMBL |
| A0A452DIH7 | Tubulin alpha chain | TUBA4A | 49481 | TrEMBL |
| A0A452DIQ5 | GLOBIN domain-containing protein | HBA1 | 14162 | TrEMBL |
| A0A452DIU4 | Tetranectin | CLEC3B | 30567 | TrEMBL |

| | | | | |
|---|---|---|---|---|
| A0A452DIX3 | Triosephosphate isomerase | TPI1 | 30577 | TrEMBL |
| A0A452DJ62 | Thrombospondin-4 | THBS4 | 105946 | TrEMBL |
| A0A452DJK6 | Serpin A3-1 | SERPINA3-1 | 52495 | TrEMBL |
| A0A452DKI9 | Xaa-Pro aminopeptidase 1 | XPNPEP1 | 74680 | TrEMBL |
| A0A493UA87 | Aldo-keto reductase family 1 member B1 | AKR1B1 | 36050 | TrEMBL |
| A0A5A4UAK7 | Heat shock protein 70 | hsp70 | 70246 | TrEMBL |
| A0A5A4UBP5 | Heat shock protein 70 | hsp70 | 70245 | TrEMBL |
| A0A6B9SBR8 | Ig lamda chain variable region (Fragment) | N/A | 11783 | TrEMBL |
| A0A6B9SCV7 | Ig lamda chain variable region (Fragment) | N/A | 11936 | TrEMBL |
| A0A8J8YKL2 | Slit-like 2 | VASN | 71707 | TrEMBL |
| A0JN91 | Exostoses (Multiple) 2 | EXT2 | 81915 | TrEMBL |
| A1L528 | RAB1A member RAS onco family | RAB1A | 22678 | TrEMBL |
| A1L548 | Ubiquitinyl hydrolase 1 (Fragment) | USP5 | 80066 | TrEMBL |
| A2T1U6 | Aminopeptidase B | AP-B | 71976 | TrEMBL |
| A2VDN8 | Coronin | CORO1C | 53126 | TrEMBL |
| A4FUD0 | C-1-tetrahydrofolate synthase cytoplasmic | MTHFD1 | 101177 | TrEMBL |
| A4FV50 | LIM and senescent cell antigen-like-containing domain protein | MGC142792 | 38422 | TrEMBL |
| A4FV56 | RNPEP protein | RNPEP | 72155 | TrEMBL |
| A4IFA5 | VASN protein | VASN | 71707 | TrEMBL |
| A4IFI0 | IGLL1 protein | IGLL1 | 24764 | TrEMBL |
| A4IFM8 | Actin alpha 1 skeletal muscle | ACTA1 | 42023 | TrEMBL |
| A5D7C6 | Prolyl endopeptidase | PREP | 80741 | TrEMBL |
| A5D7E1 | EH domain containing 3 | EHD3 | 60931 | TrEMBL |
| A5D7E8 | Protein disulfide-isomerase | PDIA3 | 56930 | TrEMBL |
| A5D7J0 | ACTA2 protein | ACTA2 | 42037 | TrEMBL |
| A5D7J6 | Calreticulin | CALR | 48098 | TrEMBL |
| A5D7L1 | C-type lectin domain containing 11A | CLEC11A | 35615 | TrEMBL |
| A5D7R6 | ITIH2 protein | ITIH2 | 106186 | TrEMBL |
| A5D7S0 | NAPA protein | NAPA | 33205 | TrEMBL |
| A5D973 | Alpha isoform of regulatory subunit A protein phosphatase 2 | PPP2R1A | 65282 | TrEMBL |
| A5D984 | Pyruvate kinase | PKM | 57949 | TrEMBL |
| A5D9B6 | Syntenin | SDCBP | 32415 | TrEMBL |
| A5D9E8 | Mimecan | OGN | 34196 | TrEMBL |
| A5PJ69 | SERPINA10 protein | SERPINA10 | 51988 | TrEMBL |
| A5PJR3 | Dicarbonyl/L-xylulose reductase | DCXR | 25677 | TrEMBL |
| A5PJT7 | ECM1 protein | ECM1 | 57637 | TrEMBL |
| A5PKC2 | SHBG protein | SHBG | 43316 | TrEMBL |

| | | | | |
|---|---|---|---|---|
| A5PKM0 | Glutathione transferase (Fragment) | GSTM2 | 26842 | TrEMBL |
| A6H7D3 | KRT18 protein (Fragment) | KRT18 | 49248 | TrEMBL |
| A6QL77 | Procollagen-proline 4-dioxygenase | P4HA1 | 60928 | TrEMBL |
| A6QLB3 | ITGA2B protein | ITGA2B | 113702 | TrEMBL |
| A6QLB7 | Adenylyl cyclase-associated protein | CAP1 | 51202 | TrEMBL |
| A6QLQ6 | PARVB protein | PARVB | 41712 | TrEMBL |
| A6QLS9 | RAB10 protein | RAB10 | 22541 | TrEMBL |
| A6QLT9 | Alanine--tRNA ligase | AARS1 | 106655 | TrEMBL |
| A6QNJ8 | GANAB protein (Fragment) | GANAB | 109085 | TrEMBL |
| A6QNL0 | Monocyte differentiation antigen CD14 | CD14 | 39922 | TrEMBL |
| A6QNW8 | Adhesion G protein-coupled receptor E5 | ADGRE5 | 80305 | TrEMBL |
| A6QP30 | CPN2 protein | CPN2 | 60040 | TrEMBL |
| A6QPH7 | AKR7A2 protein | AKR7A2 | 40508 | TrEMBL |
| A6QPP2 | SERPIND1 protein | SERPIND1 | 55207 | TrEMBL |
| A6QQP5 | SPN protein | SPN | 49618 | TrEMBL |
| A6QR28 | Phosphoserine aminotransferase | PSAT1 | 40529 | TrEMBL |
| A7E3Q2 | Heat shock 70kDa protein 1A | HSPA2 | 69809 | TrEMBL |
| A7E3S8 | Heat shock 70kD protein binding protein | ST13 | 41445 | TrEMBL |
| A7MAZ2 | STX12 protein | STX12 | 31356 | TrEMBL |
| A7MBE8 | Beta-ureidopropionase 1 | UPB1 | 42816 | TrEMBL |
| A7MBH9 | G protein subunit alpha i2 | GNAI2 | 40477 | TrEMBL |
| A7MBI6 | GLOD4 protein | GLOD4 | 33253 | TrEMBL |
| A7YW37 | CD58 protein (Fragment) | CD58 | 28707 | TrEMBL |
| A7YY67 | 10-formyltetrahydrofolate dehydrogenase | ALDH1L1 | 98738 | TrEMBL |
| A7Z067 | Receptor protein-tyrosine kinase | CSF1R | 107353 | TrEMBL |
| A8DBT6 | Monocyte differentiation antigen CD14 | CD14 | 39668 | TrEMBL |
| A8E4P3 | STOM protein | STOM | 31288 | TrEMBL |
| A8E645 | Glutamine--fructose-6-phosphate transaminase (isomerizing) | GFPT1 | 76743 | TrEMBL |
| B0JYK6 | Alpha-1 4 glucan phosphorylase | PYGM | 97288 | TrEMBL |
| B0JYL8 | Cofilin-1 | CFL1 | 18519 | TrEMBL |
| B0JYM5 | 14-3-3 protein theta | YWHAQ | 27764 | TrEMBL |
| B0JYN1 | Cathepsin L2 | CTSL2 | 37347 | TrEMBL |
| B0JYN2 | GTP-binding nuclear protein Ran | RAN | 24423 | TrEMBL |
| B0JYN3 | L-lactate dehydrogenase | LDHB | 36725 | TrEMBL |
| B0JYN6 | Alpha-2-HS-glycoprotein | AHSG | 38419 | TrEMBL |

| | | | | |
|---|---|---|---|---|
| B0JYN9 | Ras-related C3 botulinum toxin substrate 2 (Rho family small GTP binding protein Rac2) | RAC2 | 21424 | TrEMBL |
| B0JYP6 | IGK protein | IGK | 26320 | TrEMBL |
| B0JYQ0 | ALB protein | ALB | 69294 | TrEMBL |
| B0JYQ1 | Ferritin | FTH1 | 21070 | TrEMBL |
| B1PK18 | 1 4-alpha-glucan branching enzyme | N/A | 80977 | TrEMBL |
| B2KJ42 | Fructose-bisphosphatase | N/A | 36740 | TrEMBL |
| B3IVN4 | Pyruvate kinase (Fragment) | PKM | 16527 | TrEMBL |
| B3VTM3 | Lactotransferrin | N/A | 78056 | TrEMBL |
| B6VAP7 | Cell division control protein 42 homolog | CDC42 | 21243 | TrEMBL |
| B8Y9S9 | Fibronectin | N/A | 262424 | TrEMBL |
| B8Y9T0 | Fibronectin | N/A | 249128 | TrEMBL |
| B8YB76 | Homogentisate 1 2-dioxygenase | HGD | 49997 | TrEMBL |
| B9VPZ5 | Lactotransferrin | LF | 78056 | TrEMBL |
| B9X245 | Factor VIII | F8 | 263460 | TrEMBL |
| C7FE01 | Lactoferrin (Fragment) | N/A | 76275 | TrEMBL |
| D1Z308 | Periostin | POSTN | 93194 | TrEMBL |
| D4QBB3 | Hemoglobin beta | HBB | 15979 | TrEMBL |
| D4QBB4 | Globin A1 | HBB | 15954 | TrEMBL |
| E1B726 | Plasminogen | PLG | 91243 | TrEMBL |
| E1B805 | Uncharacterized protein | LOC528040 | 186351 | TrEMBL |
| E1B9F6 | Elongation factor 1-alpha | N/A | 48049 | TrEMBL |
| E1B9H5 | Transforming growth factor beta receptor 3 | TGFBR3 | 94587 | TrEMBL |
| E1BB08 | EMAP like 2 | EML2 | 70990 | TrEMBL |
| E1BB91 | Collagen type VI alpha 3 chain | COL6A3 | 339586 | TrEMBL |
| E1BBT0 | Aldo-keto reductase family 1 member D1 | AKR1D1 | 37510 | TrEMBL |
| E1BBY7 | Heat shock 70 kDa protein 4 | HSPA4 | 94509 | TrEMBL |
| E1BCW0 | HGF activator | HGFAC | 69560 | TrEMBL |
| E1BCW3 | ATP-dependent 6-phosphofructokinase | PFKP | 102968 | TrEMBL |
| E1BD93 | Urocanate hydratase | UROC1 | 82813 | TrEMBL |
| E1BEN4 | GDP-mannose pyrophosphorylase A | GMPPA | 46208 | TrEMBL |
| E1BFV0 | Karyopherin subunit beta 1 | KPNB1 | 97227 | TrEMBL |
| E1BGJ0 | LDL receptor related protein 1 | LRP1 | 504920 | TrEMBL |
| E1BH06 | Complement component 4A | C4A | 192796 | TrEMBL |
| E1BH17 | Glutathione transferase | GSTM4 | 25688 | TrEMBL |
| E1BI98 | Collagen type VI alpha 1 chain | COL6A1 | 108671 | TrEMBL |
| E1BIG6 | Transferrin receptor protein 1 | TFRC | 85441 | TrEMBL |
| E1BJ49 | MBL associated serine protease 2 | MASP2 | 76385 | TrEMBL |
| E1BJ78 | Sulfotransferase | SULT1C4 | 34900 | TrEMBL |

| | | | | |
|---|---|---|---|---|
| E1BJB1 | Tubulin beta chain | TUBB2A | 49907 | TrEMBL |
| E1BJK2 | Tubulin beta chain | TUBB1 | 49987 | TrEMBL |
| E1BKL9 | FAT atypical cadherin 1 | FAT1 | 505872 | TrEMBL |
| E1BKM4 | Programmed cell death 6 interacting protein | PDCD6IP | 96877 | TrEMBL |
| E1BKX1 | Hyaluronoglucosaminidase | CEMIP | 152767 | TrEMBL |
| E1BMG9 | 10-formyltetrahydrofolate dehydrogenase | ALDH1L1 | 98753 | TrEMBL |
| E1BMJ0 | Serpin family G member 1 | SERPING1 | 51772 | TrEMBL |
| E1BMK7 | GB1/RHD3-type G domain-containing protein | LOC107131333 | 72653 | TrEMBL |
| E1BNR0 | Apolipoprotein B | APOB | 515521 | TrEMBL |
| E1BP91 | Aminopeptidase | NPEPPS | 103600 | TrEMBL |
| E1BPI1 | Transporter | SLC6A11 | 70065 | TrEMBL |
| E9RHW1 | Heat shock protein beta-1 | HSPB1 | 22393 | TrEMBL |
| F1MAV0 | Fibrinogen beta chain | FGB | 54314 | TrEMBL |
| F1MB08 | Phosphopyruvate hydratase | ENO1 | 54151 | TrEMBL |
| F1MBC5 | Coagulation factor IX | F9 | 50358 | TrEMBL |
| F1MC48 | IQ motif containing GTPase activating protein 1 | IQGAP1 | 182781 | TrEMBL |
| F1MC75 | Parvin beta | PARVB | 40337 | TrEMBL |
| F1MCN3 | Adhesion G protein-coupled receptor E5 | ADGRE5 | 90442 | TrEMBL |
| F1MD66 | Annexin | ANXA11 | 54212 | TrEMBL |
| F1MDH3 | Talin 1 | TLN1 | 268202 | TrEMBL |
| F1MEF6 | Integrin subunit alpha 4 | ITGA4 | 115539 | TrEMBL |
| F1MG05 | Elongation factor 1-gamma | EEF1G | 50362 | TrEMBL |
| F1MGU7 | Fibrinogen gamma-B chain | FGG | 50232 | TrEMBL |
| F1MH27 | Chitinase | CHIA | 52069 | TrEMBL |
| F1MHP6 | Adenylosuccinate lyase | ADSL | 55465 | TrEMBL |
| F1MHT1 | Glycogen debranching enzyme | AGL | 174086 | TrEMBL |
| F1MI32 | Sodium/potassium-transporting ATPase subunit alpha | ATP1A1 | 112686 | TrEMBL |
| F1MIE6 | Cation-independent mannose-6-phosphate receptor | IGF2R | 274541 | TrEMBL |
| F1MJ12 | Complement C1s subcomponent | C1S | 77382 | TrEMBL |
| F1MJ28 | Alpha-1 4 glucan phosphorylase | PYGM | 97279 | TrEMBL |
| F1MJH1 | Gelsolin | GSN | 80703 | TrEMBL |
| F1MJQ1 | Ras-related protein Rab-7a | RAB7A | 23516 | TrEMBL |
| F1MJQ3 | Alpha-amylase | AMY2B | 57366 | TrEMBL |
| F1MKG2 | Collagen type VI alpha 2 chain | COL6A2 | 105177 | TrEMBL |
| F1MKS5 | Histidine-rich glycoprotein | HRG | 60743 | TrEMBL |
| F1MLI8 | Podocalyxin | PODXL | 53870 | TrEMBL |
| F1MLW8 | Uncharacterized protein | LOC100847119 | 24624 | TrEMBL |
| F1MM83 | 6-phosphogluconolactonase | PGLS | 27531 | TrEMBL |

| | | | | |
|---|---|---|---|---|
| F1MM86 | Complement component C6 | C6 | 104537 | TrEMBL |
| F1MMJ5 | Fermitin family homolog 3 | FERMT3 | 74675 | TrEMBL |
| F1MMK0 | Catechol O-methyltransferase | COMT | 48042 | TrEMBL |
| F1MMK2 | Glucose-6-phosphate 1-dehydrogenase | G6PD | 62414 | TrEMBL |
| F1MMK9 | Protein AMBP | KIF12 | 53002 | TrEMBL |
| F1MNA6 | Formimidoyltransferase-cyclodeaminase | FTCD | 60005 | TrEMBL |
| F1MNL4 | Neogenin 1 | NEO1 | 159757 | TrEMBL |
| F1MNM2 | Phosphatidylinositol-glycan-specific phospholipase D | GPLD1 | 93588 | TrEMBL |
| F1MNN7 | Lipopolysaccharide-binding protein | LBP | 53683 | TrEMBL |
| F1MNV5 | Kininogen-1 | KNG1 | 48422 | TrEMBL |
| F1MPD1 | Mannose receptor C type 2 | MRC2 | 166149 | TrEMBL |
| F1MPH3 | Thyroglobulin | TG | 302144 | TrEMBL |
| F1MPK6 | Tenascin XB | TNXB | 442587 | TrEMBL |
| F1MQ37 | Myosin heavy chain 9 | MYH9 | 227202 | TrEMBL |
| F1MRY9 | Aldehyde oxidase | AOX1 | 144703 | TrEMBL |
| F1MRZ8 | Pleckstrin | PLEK | 40057 | TrEMBL |
| F1MS05 | Cytoplasmic aconitate hydratase | ACO1 | 98232 | TrEMBL |
| F1MS32 | Apolipoprotein D | APOD | 24032 | TrEMBL |
| F1MSF1 | Ribonucleoside-diphosphate reductase | RRM1 | 89991 | TrEMBL |
| F1MSW0 | Meprin A subunit | MEP1A | 81400 | TrEMBL |
| F1MT41 | Prostaglandin F2 receptor inhibitor | PTGFRN | 98660 | TrEMBL |
| F1MTN1 | Integrin beta | ITGB3 | 86536 | TrEMBL |
| F1MTP5 | WD repeat-containing protein 1 | WDR1 | 66277 | TrEMBL |
| F1MU24 | Alpha-1 4 glucan phosphorylase | PYGB | 96339 | TrEMBL |
| F1MU79 | Peptidylprolyl isomerase | FKBP4 | 51560 | TrEMBL |
| F1MUC5 | Collagen type XV alpha 1 chain | COL15A1 | 139871 | TrEMBL |
| F1MUT4 | Coagulation factor XI | F11 | 69886 | TrEMBL |
| F1MVI0 | Contactin-1 | CNTN1 | 113306 | TrEMBL |
| F1MVP0 | ADAM metallopeptidase with thrombospondin type 1 motif 13 | ADAMTS13 | 102845 | TrEMBL |
| F1MVS9 | MBL associated serine protease 1 | MASP1 | 81299 | TrEMBL |
| F1MW44 | Coagulation factor XIII A chain | F13A1 | 82804 | TrEMBL |
| F1MWE0 | Proteasome 26S subunit ATPase 3 | PSMC3 | 49317 | TrEMBL |
| F1MWU9 | Heat shock protein family A (Hsp70) member 6 | HSPA6 | 70571 | TrEMBL |
| F1MX69 | Phosphoglycerate mutase | BPGM | 30070 | TrEMBL |

| | | | | |
|---|---|---|---|---|
| F1MX81 | Receptor protein-tyrosine kinase | TIE1 | 123897 | TrEMBL |
| F1MX87 | Complement C8 alpha chain | C8A | 66277 | TrEMBL |
| F1MXQ3 | FAM20C golgi associated secretory pathway kinase | FAM20C | 65851 | TrEMBL |
| F1MY85 | Complement C5a anaphylatoxin | C5 | 188795 | TrEMBL |
| F1MYP9 | STEAP3 metalloreductase | STEAP3 | 57886 | TrEMBL |
| F1MYX2 | Apolipoprotein M | APOM | 21158 | TrEMBL |
| F1MZ96 | Uncharacterized protein | N/A | 26664 | TrEMBL |
| F1MZC0 | Aldo_ket_red domain-containing protein | AKR7A2 | 40566 | TrEMBL |
| F1MZN7 | Sorting nexin-2 | SNX2 | 58435 | TrEMBL |
| F1MZY2 | Glutamine--fructose-6-phosphate transaminase (isomerizing) | GFPT1 | 76752 | TrEMBL |
| F1N076 | Ceruloplasmin | CP | 122395 | TrEMBL |
| F1N0I3 | Coagulation factor V | F5 | 239544 | TrEMBL |
| F1N0R5 | von Willebrand factor | VWF | 308143 | TrEMBL |
| F1N0W1 | Uncharacterized protein | DIAPH1 | 134730 | TrEMBL |
| F1N102 | Complement component C8 beta chain | C8B | 66686 | TrEMBL |
| F1N169 | Filamin A | FLNA | 280925 | TrEMBL |
| F1N1I6 | Gelsolin | GSN | 92084 | TrEMBL |
| F1N2B5 | Solute carrier family 3 member 2 | SLC3A2 | 59011 | TrEMBL |
| F1N2L9 | 4-trimethylaminobutyraldehyde dehydrogenase | ALDH9A1 | 56704 | TrEMBL |
| F1N3A1 | Thrombospondin-1 | THBS1 | 129392 | TrEMBL |
| F1N3F7 | RELT TNF receptor | RELT | 45475 | TrEMBL |
| F1N3P2 | Ubiquitin carboxyl-terminal hydrolase | USP5 | 95815 | TrEMBL |
| F1N3Q7 | Apolipoprotein A-IV | APOA4 | 52291 | TrEMBL |
| F1N441 | Aldehyde dehydrogenase family 16 member A1 | ALDH16A1 | 85226 | TrEMBL |
| F1N468 | Adenosine kinase | ADK | 38532 | TrEMBL |
| F1N4K1 | Phosphoribosylformylglycinamidine synthase | PFAS | 147728 | TrEMBL |
| F1N4M7 | Complement factor I | CFI | 68013 | TrEMBL |
| F1N549 | Dihydropyrimidine dehydrogenase [NADP(+)] | DPYD | 111838 | TrEMBL |
| F1N5M2 | Vitamin D-binding protein | GC | 53356 | TrEMBL |
| F1N647 | Fatty acid synthase | FASN | 274408 | TrEMBL |
| F1N6Y1 | Glucosidase II alpha subunit | GANAB | 109472 | TrEMBL |
| F2Z4D5 | Ras-related protein Rab-11A | RAB11A | 24394 | TrEMBL |
| F2Z4F5 | Dipeptidyl peptidase 3 | DPP3 | 83802 | TrEMBL |
| F5XVA9 | von Willebrand factor | VWF | 307947 | TrEMBL |
| F6PZI3 | Solute carrier family 29 member 1 | SLC29A1 | 57937 | TrEMBL |

| | | | | |
|---|---|---|---|---|
| F6Q334 | Alpha-soluble NSF attachment protein | NAPA | 56576 | TrEMBL |
| F6Q751 | Glutathione transferase | GSTM4 | 25654 | TrEMBL |
| F6QGM3 | Sulfurtransferase | MPST | 36159 | TrEMBL |
| F6QHN4 | Prolyl endopeptidase | PREP | 81443 | TrEMBL |
| F6QND5 | Fibrinogen alpha chain | FGA | 94734 | TrEMBL |
| F6QPG2 | Anion exchange protein | SLC4A1 | 105766 | TrEMBL |
| F6QS88 | Epoxide hydrolase 2 | EPHX2 | 62721 | TrEMBL |
| F6R4P6 | Serpin family D member 1 | SERPIND1 | 62705 | TrEMBL |
| F6R6T6 | Adhesion G protein-coupled receptor E5 | ADGRE5 | 85509 | TrEMBL |
| F6RJG0 | Hydroxymethylglutaryl-CoA synthase | HMGCS1 | 63531 | TrEMBL |
| F6RMV5 | Leucine rich alpha-2-glycoprotein 1 | LRG1 | 39193 | TrEMBL |
| F6RQK3 | Maleylacetoacetate isomerase | GSTZ1 | 24205 | TrEMBL |
| F6S1Q0 | Keratin 18 | KRT18 | 47965 | TrEMBL |
| G3MYK9 | Semaphorin 3G | SEMA3G | 85909 | TrEMBL |
| G3MZL1 | Cadherin-5 | CDH5 | 87463 | TrEMBL |
| G3N0S9 | Uncharacterized protein | N/A | 21488 | TrEMBL |
| G3X6L9 | Glutamyl-prolyl-tRNA synthetase 1 | EPRS1 | 169930 | TrEMBL |
| G3X6N3 | Serotransferrin | TF | 77738 | TrEMBL |
| G3X6Y4 | Osteomodulin | OMD | 49143 | TrEMBL |
| G3X743 | Arginyl aminopeptidase | RNPEP | 72141 | TrEMBL |
| G3X755 | Plexin B2 | PLXNB2 | 215014 | TrEMBL |
| G3X7N4 | Phosphoglycerate kinase | PGK1 | 43123 | TrEMBL |
| G5E507 | Heat shock protein HSP 90-beta | HSP90AB1 | 81723 | TrEMBL |
| G5E5A8 | Fibronectin | FN1 | 275548 | TrEMBL |
| G5E5A9 | Fibronectin | FN1 | 285245 | TrEMBL |
| G5E5B0 | Fibronectin | FN1 | 275417 | TrEMBL |
| G5E5C8 | Transaldolase | TALDO1 | 37681 | TrEMBL |
| G5E5T5 | Uncharacterized protein | N/A | 56043 | TrEMBL |
| G5E5U7 | S-adenosylmethionine synthase | MAT1A | 43789 | TrEMBL |
| G5E5V0 | Carboxypeptidase N catalytic chain | CPN1 | 52635 | TrEMBL |
| G5E5W1 | Coagulation factor VIII | F8 | 263542 | TrEMBL |
| G5E619 | GTP-binding nuclear protein Ran | N/A | 24467 | TrEMBL |
| G8JKX4 | Actin aortic smooth muscle | ACTA2 | 41777 | TrEMBL |
| H2CNR1 | Peptidoglycan-recognition protein | PGLYRP1 | 21037 | TrEMBL |
| H7BWW0 | Coronin | CORO1B | 54253 | TrEMBL |
| I7CT57 | Vitamin D-binding protein | N/A | 53328 | TrEMBL |
| J9QD97 | Periostin variant 9 | N/A | 93160 | TrEMBL |
| J9QDG2 | Periostin variant 8 | N/A | 89748 | TrEMBL |

| | | | | |
|---|---|---|---|---|
| K8FK38 | Heat Shock Protein 70 (Fragment) | HSP70 | 69421 | TrEMBL |
| M1ZMP2 | Aldehyde oxidase | N/A | 147310 | TrEMBL |
| M5FKF4 | Insulin-like growth factor binding protein acid labile subunit | IGFALS | 65993 | TrEMBL |
| O18977 | Tenascin-X | TN-X | 447387 | TrEMBL |
| Q08DL0 | SLC3A2 protein | SLC3A2 | 63183 | TrEMBL |
| Q08DW4 | Mannan-binding lectin serine peptidase 1 (C4/C2 activating component of Ra-reactive factor) | MASP1 | 81269 | TrEMBL |
| Q09TE3 | Insulin-like growth factor binding protein acid labile subunit | IGFALS | 65969 | TrEMBL |
| Q0V8K9 | Solute carrier family 29 (Nucleoside transporters) member 1 (Fragment) | SLC29A1 | 49196 | TrEMBL |
| Q0VCS8 | Glutathione transferase | GSTT3 | 27277 | TrEMBL |
| Q17QK4 | Epoxide hydrolase 2 cytoplasmic | EPHX2 | 62691 | TrEMBL |
| Q17R18 | Adenosine kinase | ADK | 38580 | TrEMBL |
| Q1JP95 | Proteasome 26S ATPase subunit 3 (Fragment) | PSMC3 | 47995 | TrEMBL |
| Q1JPA2 | Elongation factor 1-gamma (Fragment) | EEF1G | 50231 | TrEMBL |
| Q1JPD0 | Complement component 8 alpha polypeptide (Fragment) | C8A | 32642 | TrEMBL |
| Q1JPE5 | Tumor necrosis factor receptor superfamily member 19-like (Fragment) | TNFRSF19L | 37696 | TrEMBL |
| Q1JPG7 | Pyruvate kinase | PKLR | 56870 | TrEMBL |
| **Q1RMN8** | **Immunoglobulin light chain lambda gene cluster** | **IGL@** | **24536** | **TrEMBL** |
| Q1RMN9 | C4b-binding protein alpha-like | LOC510860 | 21997 | TrEMBL |
| **Q28194** | **Thrombospondin-1 (Fragment)** | **N/A** | **25015** | **TrEMBL** |
| Q2HJB6 | Procollagen C-endopeptidase enhancer | PCOLCE | 48211 | TrEMBL |
| **Q2HJF0** | **Serotransferrin-like** | **LOC525947** | **69183** | **TrEMBL** |
| **Q2KIF2** | **Leucine-rich alpha-2-glycoprotein 1** | **LRG1** | **38348** | **TrEMBL** |
| **Q2KIH2** | **ApoN protein** | **APON** | **28535** | **TrEMBL** |
| Q2KIH5 | Complement component 8 alpha polypeptide | C8A | 66335 | TrEMBL |
| Q2KIM6 | Maleylacetoacetate isomerase | GSTZ1 | 24113 | TrEMBL |
| Q2KIW4 | Lecithin-cholesterol acyltransferase | LCAT | 49844 | TrEMBL |
| Q32LP1 | Ribonucleoside-diphosphate reductase | RRM1 | 90019 | TrEMBL |
| Q32PA1 | CD59 glycoprotein | CD59 | 13663 | TrEMBL |

| | | | | |
|---|---|---|---|---|
| **Q32PI4** | **Complement factor I** | **CFI** | **68933** | **TrEMBL** |
| Q3MHG3 | Sulfurtransferase | MPST | 33193 | TrEMBL |
| **Q3MHH8** | **Alpha-amylase** | **AMY2A** | **57408** | **TrEMBL** |
| Q3MHK9 | Fascin | FSCN1 | 54785 | TrEMBL |
| Q3SYR0 | Serpin peptidase inhibitor clade A (Alpha-1 antiproteinase antitrypsin) member 7 | SERPINA7 | 46035 | TrEMBL |
| Q3SZ81 | PSMC3 protein (Fragment) | PSMC3 | 48094 | TrEMBL |
| Q3SZQ8 | Endopin 2 | SERPINA3-7 | 47034 | TrEMBL |
| Q3SZZ9 | FGG protein | FGG | 49167 | TrEMBL |
| Q3T085 | Mimecan | OGN | 34197 | TrEMBL |
| Q3T0F0 | SLC3A2 protein | SLC3A2 | 59445 | TrEMBL |
| Q3T0T9 | Carbonyl reductase (NADPH) | MGC127133 | 31707 | TrEMBL |
| Q3T0Y1 | Ubiquitin thioesterase | OTUB1 | 31308 | TrEMBL |
| Q3T101 | IGL@ protein | IGL@ | 24640 | TrEMBL |
| Q3ZBG1 | Ras-related protein Rab-14 | RAB14 | 23897 | TrEMBL |
| Q3ZBG2 | AKR1C4 protein | AKR1C4 | 36732 | TrEMBL |
| Q3ZBQ9 | Apolipoprotein M | APOM | 13027 | TrEMBL |
| **Q3ZBS7** | **Vitronectin** | **VTN** | **53575** | **TrEMBL** |
| Q3ZC30 | Sulfotransferase | SULT1E1 | 34660 | TrEMBL |
| Q3ZC79 | Hydroxymethylglutaryl-CoA synthase (Fragment) | HMGCS1 | 62059 | TrEMBL |
| Q3ZC83 | Solute carrier family 29 (Nucleoside transporters) member 1 | SLC29A1 | 49882 | TrEMBL |
| Q3ZC85 | Plasmalemma vesicle associated protein | PLVAP | 49768 | TrEMBL |
| Q3ZC87 | Pyruvate kinase (Fragment) | PKM2 | 61428 | TrEMBL |
| Q3ZCI4 | 6-phosphogluconate dehydrogenase decarboxylating | PGD | 53077 | TrEMBL |
| Q58CV8 | Sulfotransferase | SULT1C2 | 34914 | TrEMBL |
| Q58DL9 | Phospholipid transfer protein | PLTP | 56428 | TrEMBL |
| Q58DT9 | Alpha 2 actin | ACTA2 | 45221 | TrEMBL |
| Q5EA54 | Solute carrier family 3 (Activators of dibasic and neutral amino acid transport) member 2 | SLC3A2 | 63211 | TrEMBL |
| Q5EA67 | Inter-alpha (Globulin) inhibitor H4 (Plasma Kallikrein-sensitive glycoprotein) | ITIH4 | 101510 | TrEMBL |
| Q5GN72 | Alpha-1-acid glycoprotein | AGP | 23158 | TrEMBL |
| Q5J801 | Endopin 2B | N/A | 47002 | TrEMBL |
| Q5W1P3 | Putative calcium-dependent phospholipase A2 (Fragment) | PLA2G2D2 | 15241 | TrEMBL |
| Q68RU0 | Ovarian and testicular apolipoprotein N | ApoN | 29091 | TrEMBL |
| Q6LBN7 | Lactoferrin (Fragment) | N/A | 75182 | TrEMBL |

| | | | | |
|---|---|---|---|---|
| Q6QRN7 | PP1201 protein | TMBIM1 | 33949 | TrEMBL |
| Q6T182 | Sex hormone-binding globulin (Fragment) | SHBG | 40094 | TrEMBL |
| Q702I4 | CD97 antigen transcript variant | cd97 | 90459 | TrEMBL |
| Q861V5 | Peptidyl-prolyl cis-trans isomerase (Fragment) | N/A | 16926 | TrEMBL |
| Q8HZ62 | Prostaglandin F synthase-like2 protein | PGFSL2 | 36790 | TrEMBL |
| Q9BGU1 | Histidine-rich glycoprotein | BTHRG | 61948 | TrEMBL |
| Q9TU26 | Blood-brain barrier large neutral amino acid transporter | SLC7A5 | 55111 | TrEMBL |
| Q9XSW5 | Anion exchange protein | SLC4A1 | 104375 | TrEMBL |
| V6F7X3 | Apolipoprotein A-IV | APOA4 | 42990 | TrEMBL |
| V6F9A2 | Apolipoprotein A-I preproprotein | APOA1 | 30276 | TrEMBL |

N/A: non annotated gene.

**Table S2.** List of the 180 of the Contaminants Database (Max Planck Institute) included in the SPROUTS_DB.

| Accession | Description | Gene | Organism | Section of UniProt or other DB |
|---|---|---|---|---|
| P00761 | Trypsin | N/A | Sus scrofa | SwissProt |
| Q86Y46 | Keratin, type II cytoskeletal 73 | KRT73 | Homo sapiens | SwissProt |
| P19013 | Keratin, type II cytoskeletal 4 | KRT4 | Homo sapiens | SwissProt |
| Q8N1N4 | Keratin, type II cytoskeletal 78 | KRT78 | Homo sapiens | SwissProt |
| P15636 | Protease 1 | N/A | Achromobacter lyticus | SwissProt |
| P09870 | Clostripain | cloSI | Hathewaya histolytica | SwissProt |
| Q9R4J4 | Peptidyl-Asp metalloendopeptidase (Fragment) | N/A | Pseudomonas fragi | SwissProt |
| P0C1U8 | Glutamyl endopeptidase | sspA | Staphylococcu s aureus | SwissProt |
| P00766 | Chymotrypsinogen A | N/A | Bos taurus | SwissProt |
| P13717 | Nuclease | nucA | Serratia marcescens | SwissProt |
| Q9U6Y5 | GFP-like fluorescent chromoprotein FP506 | N/A | Zoanthus sp. | SwissProt |
| P21578 | Yellow fluorescent protein | luxY | Aliivibrio fischeri | SwissProt |
| O76009 | Keratin, type I cuticular Ha3-I | KRT33A | Homo sapiens | SwissProt |
| O76011 | Keratin, type I cuticular Ha4 | KRT34 | Homo sapiens | SwissProt |
| O76013 | Keratin, type I cuticular Ha6 | KRT36 | Homo sapiens | SwissProt |
| O76014 | Keratin, type I cuticular Ha7 | KRT37 | Homo sapiens | SwissProt |
| O76015 | Keratin, type I cuticular Ha8 | KRT38 | Homo sapiens | SwissProt |
| P08779 | Keratin, type I cytoskeletal 16 | KRT16 | Homo sapiens | SwissProt |
| Q14525 | Keratin, type I cuticular Ha3-II | KRT33B | Homo sapiens | SwissProt |
| Q14532 | Keratin, type I cuticular Ha2 | KRT32 | Homo sapiens | SwissProt |

| | | | | |
|---|---|---|---|---|
| Q15323 | Keratin, type I cuticular Ha1 | KRT31 | Homo sapiens | SwissProt |
| Q92764 | Keratin, type I cuticular Ha5 | KRT35 | Homo sapiens | SwissProt |
| Q14533 | Keratin, type II cuticular Hb1 | KRT81 | Homo sapiens | SwissProt |
| Q9NSB4 | Keratin, type II cuticular Hb2 | KRT82 | Homo sapiens | SwissProt |
| P78385 | Keratin, type II cuticular Hb3 | KRT83 | Homo sapiens | SwissProt |
| Q9NSB2 | Keratin, type II cuticular Hb4 | KRT84 | Homo sapiens | SwissProt |
| P78386 | Keratin, type II cuticular Hb5 | KRT85 | Homo sapiens | SwissProt |
| O43790 | Keratin, type II cuticular Hb6 | KRT86 | Homo sapiens | SwissProt |
| Q6A162 | Keratin, type I cytoskeletal 40 | KRT40 | Homo sapiens | SwissProt |
| O76011 | Keratin, type I cuticular Ha1 | KRT31 | Homo sapiens | SwissProt |
| O76011 | Keratin, type I cuticular Ha4 | KRT34 | Homo sapiens | SwissProt |
| P78385 | Keratin, type II cuticular Hb3 | KRT83 | Homo sapiens | SwissProt |
| Q9NSB2 | Keratin, type II cuticular Hb4 | KRT84 | Homo sapiens | SwissProt |
| O76009 | Keratin, type I cuticular Ha3-I | KRT33A | Homo sapiens | SwissProt |
| Q6A163 | Keratin, type I cytoskeletal 39 | KRT39 | Homo sapiens | SwissProt |
| P04264 | Keratin, type II cytoskeletal 1 | KRT1 | Homo sapiens | SwissProt |
| P13647 | Keratin, type II cytoskeletal 5 | KRT5 | Homo sapiens | SwissProt |
| P35908 | Keratin, type II cytoskeletal 2 epidermal | KRT2 | Homo sapiens | SwissProt |
| P13645 | Keratin, type I cytoskeletal 10 | KRT10 | Homo sapiens | SwissProt |
| P35527 | Keratin, type I cytoskeletal 9 | KRT9 | Homo sapiens | SwissProt |
| Q6E0U4 | Dermokine | DMKN | Homo sapiens | SwissProt |
| P02533 | Keratin, type I cytoskeletal 14 | KRT14 | Homo sapiens | SwissProt |
| P02538 | Keratin, type II cytoskeletal 6A | KRT6A | Homo sapiens | SwissProt |
| P48668 | Keratin, type II cytoskeletal 6C | KRT6C | Homo sapiens | SwissProt |
| P04259 | Keratin, type II cytoskeletal 6B | KRT6B | Homo sapiens | SwissProt |
| Q6E0U4 | Dermokine | DMKN | Homo sapiens | SwissProt |
| Q3SZH5 | Angiotensinogen | AGT | Bos taurus | TrEMBL |
| Q1A7A4 | Complement component C5a (Fragment) | N/A | Bos taurus | TrEMBL |
| Q2YDI2 | Origin recognition complex subunit 4 | ORC4 | Bos taurus | SwissProt |
| Q2KJC7 | Periostin | POSTN | Bos taurus | TrEMBL |
| P02777 | Platelet factor 4 | PF4 | Bos taurus | SwissProt |
| Q6T181 | Sex hormone-binding globulin (Fragment) | SHBG | Bos taurus | TrEMBL |
| P21752 | Thymosin beta-10 | TMSB10 | Bos taurus | SwissProt |
| Q2KIU3 | Protein HP-25 homolog 2 | N/A | Bos taurus | SwissProt |
| P67983 | Metallothionein-1A | MT1A | Bos taurus | SwissProt |
| Q862S4 | Similar to pro alpha 1(I) collagen (Fragment) | N/A | Bos taurus | TrEMBL |
| Q5KR48 | Tropomyosin beta chain | TPM2 | Bos taurus | SwissProt |
| P56652 | Inter-alpha-trypsin inhibitor heavy chain H3 | ITIH3 | Bos taurus | SwissProt |
| P13646 | Keratin, type I cytoskeletal 13 | KRT13 | Homo sapiens | SwissProt |
| Q04695 | Keratin, type I cytoskeletal 17 | KRT17 | Homo sapiens | SwissProt |
| A2I7N0 | Serpin A3-4 | SERPINA3-4 | Bos taurus | SwissProt |
| P02768 | Albumin | ALB | Homo sapiens | SwissProt |

| | | | | |
|---|---|---|---|---|
| P02081 | Hemoglobin fetal subunit beta | N/A | Bos taurus | SwissProt |
| Q9TRI1 | Inter-alpha-trypsin inhibitor HC2 component homolog (Fragment) | N/A | Bos taurus | TrEMBL |
| Q3KUS7 | Complement factor B (Fragment) | BF | Bos taurus | TrEMBL |
| Q2TBQ1 | Coagulation factor XIII B chain | F13B | Bos taurus | TrEMBL |
| Q05B55 | IGK protein | IGK | Bos taurus | TrEMBL |
| P01044 | Kininogen-1 | KNG1 | Bos taurus | SwissProt |
| Q29443 | Serotransferrin | TF | Bos taurus | SwissProt |
| P02662 | Alpha-S1-casein | CSN1S1 | Bos taurus | SwissProt |
| P02663 | Alpha-S2-casein | CSN1S2 | Bos taurus | SwissProt |
| P02666 | Beta-casein | CSN2 | Bos taurus | SwissProt |
| P02668 | Kappa-casein | CSN3 | Bos taurus | SwissProt |
| P31096 | Osteopontin | SPP1 | Bos taurus | SwissProt |
| P02754 | Beta-lactoglobulin | LGB | Bos taurus | SwissProt |
| P00711 | Alpha-lactalbumin | LALBA | Bos taurus | SwissProt |
| P62894 | Cytochrome c | CYCS | Bos taurus | SwissProt |
| P19001 | Keratin, type I cytoskeletal 19 | Krt19 | Mus musculus | SwissProt |
| Q8VED5 | Keratin, type II cytoskeletal 79 | Krt79 | Mus musculus | SwissProt |
| Q61726 | Keratin type II (Fragment) | Krt83 | Mus musculus | TrEMBL |
| Q3ZAW8 | Keratin 16 | Krt16 | Mus musculus | TrEMBL |
| P50446 | Keratin, type II cytoskeletal 6A | Krt6a | Mus musculus | SwissProt |
| Q497I4 | Keratin, type I cuticular Ha5 | Krt35 | Mus musculus | SwissProt |
| Q9D312 | Keratin, type I cytoskeletal 20 | Krt20 | Mus musculus | SwissProt |
| P08730 | Keratin, type I cytoskeletal 13 | Krt13 | Mus musculus | SwissProt |
| Q922U2 | Keratin, type II cytoskeletal 5 | Krt5 | Mus musculus | SwissProt |
| Q8BGZ7 | Keratin, type II cytoskeletal 75 | Krt75 | Mus musculus | SwissProt |
| Q9QWL7 | Keratin, type I cytoskeletal 17 | Krt17 | Mus musculus | SwissProt |
| Q6IME9 | Keratin, type II cytoskeletal 72 | Krt72 | Mus musculus | SwissProt |
| Q6NXH9 | Keratin, type II cytoskeletal 73 | Krt73 | Mus musculus | SwissProt |
| Q6IFX4 | Keratin, type I cytoskeletal 39 | Krt39 | Mus musculus | SwissProt |
| P07744 | Keratin, type II cytoskeletal 4 | Krt4 | Mus musculus | SwissProt |
| Q6IFZ6 | Keratin, type II cytoskeletal 1b | Krt77 | Mus musculus | SwissProt |
| Q6IFX2 | Keratin, type I cytoskeletal 42 | Krt42 | Mus musculus | SwissProt |
| Q9R0H5 | Keratin, type II cytoskeletal 71 | Krt71 | Mus musculus | SwissProt |
| Q3TTY5 | Keratin, type II cytoskeletal 2 epidermal | Krt2 | Mus musculus | SwissProt |
| Q0VBK2 | Keratin, type II cytoskeletal 80 | Krt80 | Mus musculus | SwissProt |
| P02535 | Keratin, type I cytoskeletal 10 | Krt10 | Mus musculus | SwissProt |
| Q61782 | Type I epidermal keratin mRNA, 3'end (Fragment) | N/A | Mus musculus | TrEMBL |
| Q61765 | Keratin, type I cuticular Ha1 | Krt31 | Mus musculus | SwissProt |
| Q99PS0 | Keratin, type I cytoskeletal 23 | Krt23 | Mus musculus | SwissProt |
| Q9D646 | Keratin, type I cuticular Ha4 | Krt34 | Mus musculus | SwissProt |
| P05784 | Keratin, type I cytoskeletal 18 | Krt18 | Mus musculus | SwissProt |
| Q9DCV7 | Keratin, type II cytoskeletal 7 | Krt7 | Mus musculus | SwissProt |
| Q9Z2K1 | Keratin, type I cytoskeletal 16 | Krt16 | Mus musculus | SwissProt |

| | | | | |
|---|---|---|---|---|
| P07477 | Serine protease 1 | PRSS1 | Homo sapiens | SwissProt |
| P05787 | Keratin, type II cytoskeletal 8 | KRT8 | Homo sapiens | SwissProt |
| Q6KB66 | Keratin, type II cytoskeletal 80 | KRT80 | Homo sapiens | SwissProt |
| Q7Z794 | Keratin, type II cytoskeletal 1b | KRT77 | Homo sapiens | SwissProt |
| Q9BYR9 | Keratin-associated protein 2-4 | KRTAP2-4 | Homo sapiens | SwissProt |
| Q9BYQ5 | Keratin-associated protein 4-6 | KRTAP4-6 | Homo sapiens | SwissProt |
| Q9BYR8 | Keratin-associated protein 3-1 | KRTAP3-1 | Homo sapiens | SwissProt |
| Q9BYQ7 | Keratin-associated protein 4-1 | KRTAP4-1 | Homo sapiens | SwissProt |
| Q3LI72 | Keratin-associated protein 19-5 | KRTAP19-5 | Homo sapiens | SwissProt |
| Q9BYR4 | Keratin-associated protein 4-3 | KRTAP4-3 | Homo sapiens | SwissProt |
| Q9BYQ8 | Keratin-associated protein 4-9 | KRTAP4-9 | Homo sapiens | SwissProt |
| P60413 | Keratin-associated protein 10-12 | KRTAP10-12 | Homo sapiens | SwissProt |
| P19012 | Keratin, type I cytoskeletal 15 | KRT15 | Homo sapiens | SwissProt |
| Q2M2I5 | Keratin, type I cytoskeletal 24 | KRT24 | Homo sapiens | SwissProt |
| O95678 | Keratin, type II cytoskeletal 75 | KRT75 | Homo sapiens | SwissProt |
| Q01546 | Keratin, type II cytoskeletal 2 oral | KRT76 | Homo sapiens | SwissProt |
| Q99456 | Keratin, type I cytoskeletal 12 | KRT12 | Homo sapiens | SwissProt |
| P35900 | Keratin, type I cytoskeletal 20 | KRT20 | Homo sapiens | SwissProt |
| Q3SY84 | Keratin, type II cytoskeletal 71 | KRT71 | Homo sapiens | SwissProt |
| Q8N1A0 | Keratin-like protein KRT222 | KRT222 | Homo sapiens | SwissProt |
| Q8N1N4 | Keratin, type II cytoskeletal 78 | KRT78 | Homo sapiens | SwissProt |
| Q5XKE5 | Keratin, type II cytoskeletal 79 | KRT79 | Homo sapiens | SwissProt |
| P12035 | Keratin, type II cytoskeletal 3 | KRT3 | Homo sapiens | SwissProt |
| Q9C075 | Keratin, type I cytoskeletal 23 | KRT23 | Homo sapiens | SwissProt |
| P08729 | Keratin, type II cytoskeletal 7 | KRT7 | Homo sapiens | SwissProt |
| Q7Z3Y8 | Keratin, type I cytoskeletal 27 | KRT27 | Homo sapiens | SwissProt |
| Q7RTS7 | Keratin, type II cytoskeletal 74 | KRT74 | Homo sapiens | SwissProt |
| Q7Z3Y9 | Keratin, type I cytoskeletal 26 | KRT26 | Homo sapiens | SwissProt |
| Q7Z3Z0 | Keratin, type I cytoskeletal 25 | KRT25 | Homo sapiens | SwissProt |
| Q7Z3Y7 | Keratin, type I cytoskeletal 28 | KRT28 | Homo sapiens | SwissProt |
| P08727 | Keratin, type I cytoskeletal 19 | KRT19 | Homo sapiens | SwissProt |
| Q14CN4 | Keratin, type II cytoskeletal 72 | KRT72 | Homo sapiens | SwissProt |
| Q3KNV1 | KRT7 protein | KRT7 | Homo sapiens | TrEMBL |
| Q86YZ3 | Hornerin | HRNR | Homo sapiens | SwissProt |
| P20930 | Filaggrin | FLG | Homo sapiens | SwissProt |
| Q5D862 | Filaggrin-2 | FLG2 | Homo sapiens | SwissProt |
| P22629 | Streptavidin | N/A | Streptomyces avidinii | SwissProt |
| Q1RMK2 | IGHM protein | N/A | Bos taurus | TrEMBL |
| A2AB72 | Keratin complex 1, acidic, gene 2 | Krt32 | Mus musculus | TrEMBL |
| A2A4G1 | Keratin complex 1, acidic, gene 15 | Krt15 | Mus musculus | TrEMBL |
| Q9H552 | Keratin-8-like protein 1 | N/A | Homo sapiens | TrEMBL |
| XP_986630 | keratin complex 1, acidic, gene 3 | Krt33b | Mus musculus | REFSEQ |

| | | | | |
|---|---|---|---|---|
| XP_001474382 | RIKEN cDNA 1110025L11 | 1110025L11Rik | Mus musculus | REFSEQ |
| XP_092267 | similar to Keratin, type II cytoskeletal 8 | LOC150739 | Homo sapiens | REFSEQ |
| XP_932229 | similar to Keratin, type II cytoskeletal 2 oral | KRT126P | Homo sapiens | REFSEQ |
| HIT000016045 | Similar to Keratin, type II cytoskeletal 8 | N/A | Homo sapiens | H-INV |
| HIT000292931 | Similar to Keratin, type II cytoskeletal 8 | N/A | Homo sapiens | H-INV |
| HIT000015463 | Similar to Keratin 18 | PTPN14 | Homo sapiens | H-INV |
| ENSP00000377550 | 46 kDa protein | KRT13 | Homo sapiens | ENSEMBL |
| ENSBTAP00000006074 | 81 kDa protein | N/A | Bos taurus | ENSEMBL |
| ENSBTAP00000038329 | 9 kDa protein | N/A | Bos taurus | ENSEMBL |
| XP_001252647 | similar to endopin 2B | N/A | Bos taurus | REFSEQ |
| ENSBTAP00000007350 | similar to Complement C4-A precursor | N/A | Bos taurus | ENSEMBL |
| ENSBTAP00000038253 | 63 kDa protein | N/A | Bos taurus | ENSEMBL |
| ENSBTAP00000023402 | 46 kDa protein | N/A | Bos taurus | ENSEMBL |
| ENSBTAP00000024466 | 44 kDa protein | N/A | Bos taurus | ENSEMBL |
| ENSBTAP00000023055 | 68 kDa protein | N/A | Bos taurus | ENSEMBL |
| ENSBTAP00000018229 | 54 kDa protein | N/A | Bos taurus | ENSEMBL |
| ENSBTAP00000016046 | similar to fibulin-1 C isoform 1 | N/A | Bos taurus | ENSEMBL |
| ENSBTAP00000024462 | 47 kDa protein | N/A | Bos taurus | ENSEMBL |
| ENSBTAP00000014147 | 12 kDa protein | N/A | Bos taurus | ENSEMBL |
| ENSBTAP00000033053 | 15 kDa protein | N/A | Bos taurus | ENSEMBL |
| ENSBTAP00000001528 | similar to intersectin long isoform 4 | N/A | Bos taurus | ENSEMBL |
| ENSBTAP00000037665 | similar to Pregnancy zone protein, partial | N/A | Bos taurus | ENSEMBL |
| ENSBTAP00000031900 | 121 kDa protein | N/A | Bos taurus | ENSEMBL |
| ENSBTAP00000031360 | 55 kDa protein | N/A | Bos taurus | ENSEMBL |
| ENSBTAP00000018574 | 55 kDa protein | N/A | Bos taurus | ENSEMBL |
| ENSBTAP00000032840 | similar to apolipoprotein B, partial | N/A | Bos taurus | ENSEMBL |
| ENSBTAP00000011227 | 15 kDa protein | N/A | Bos taurus | ENSEMBL |
| ENSBTAP00000025008 | hypothetical protein | N/A | Bos taurus | ENSEMBL |
| ENSBTAP00000034412 | similar to C4b-binding protein alpha chain | N/A | Bos taurus | ENSEMBL |
| ENSBTAP00000013050 | hypothetical protein | N/A | Bos taurus | ENSEMBL |
| ENSBTAP00000016285 | similar to peptidoglycan recognition protein L | N/A | Bos taurus | ENSEMBL |

| | | | | |
|---|---|---|---|---|
| ENSBTAP00000024146 | similar to alpha-2-macroglobulin isoform 1 | N/A | Bos taurus | ENSEMBL |
| XP_585019 | similar to afamin | N/A | Bos taurus | REFSEQ |

N/A: non annotated gene.

**Table S3.** List of the proteins quantified by Peaks X-Pro software. For each protein is reported: Accession Number as in the UniProt database, protein name, gene code, score, number of the characterized peptides, number of unique peptides and average area.

| Accession | Protein name | Gene code | Score (-10lgP) | Peptides | Unique | Avg Area |
|---|---|---|---|---|---|---|
| Q7SIH1 | Alpha-2-macroglobulin | A2M | 666.87 | 247 | 247 | 1.59E+10 |
| P02769 | Albumin | ALB | 547.39 | 96 | 89 | 5.53E+09 |
| P12763 | Alpha-2-HS-glycoprotein | AHSG | 520.17 | 49 | 49 | 1.56E+09 |
| Q2UVX4 | Complement C3 | C3 | 590.1 | 116 | 116 | 1.27E+09 |
| P34955 | Alpha-1-antiproteinase | SERPINA1 | 399.32 | 30 | 30 | 1.24E+09 |
| P01966 | Hemoglobin subunit alpha | HBA | 310.25 | 10 | 10 | 9.70E+08 |
| P00735 | Prothrombin | F2 | 481.91 | 50 | 50 | 6.58E+08 |
| P02081 | Hemoglobin fetal subunit beta | N/A | 395.25 | 25 | 21 | 4.67E+08 |
| P41361 | Antithrombin-III | SERPINC1 | 419.64 | 28 | 28 | 3.72E+08 |
| Q29443 | Serotransferrin | TF | 480.42 | 49 | 46 | 2.54E+08 |
| P17697 | Clusterin | CLU | 345.87 | 18 | 18 | 1.87E+08 |
| Q95121 | Pigment epithelium-derived factor | SERPINF1 | 382 | 20 | 20 | 1.42E+08 |
| Q3MHL4 | Adenosylhomocysteinase | AHCY | 363.79 | 23 | 23 | 1.04E+08 |
| Q58D62 | Fetuin-B | FETUB | 320.17 | 14 | 14 | 1.02E+08 |
| Q05443 | Lumican | LUM | 283.6 | 14 | 14 | 9.17E+07 |
| P01017 | Angiotensinogen | AGT | 411.61 | 27 | 7 | 8.96E+07 |
| Q2KJ83 | Carboxypeptidase N catalytic chain | CPN1 | 340.47 | 13 | 13 | 8.39E+07 |
| P10096 | Glyceraldehyde-3-phosphate dehydrogenase | GAPDH | 365.54 | 15 | 13 | 8.31E+07 |
| Q3T052 | Inter-alpha-trypsin inhibitor heavy chain H4 | ITIH4 | 423.89 | 31 | 31 | 7.46E+07 |
| Q28107 | Coagulation factor V | F5 | 442.83 | 50 | 50 | 7.09E+07 |
| Q3SZ57 | Alpha-fetoprotein | AFP | 404.84 | 29 | 28 | 6.91E+07 |
| O46375 | Transthyretin | TTR | 323.88 | 9 | 9 | 6.85E+07 |
| Q3Y5Z3 | Adiponectin | ADIPOQ | 234.24 | 6 | 6 | 6.48E+07 |
| P06868 | Plasminogen | PLG | 402.08 | 38 | 38 | 6.33E+07 |
| P56652 | Inter-alpha-trypsin inhibitor heavy chain H3 | ITIH3 | 362.22 | 23 | 23 | 6.32E+07 |
| P15497 | Apolipoprotein A-I | APOA1 | 308.48 | 18 | 18 | 5.10E+07 |
| Q9XSJ4 | Alpha-enolase | ENO1 | 379.4 | 23 | 17 | 4.78E+07 |
| P81187 | Complement factor B | CFB | 393.22 | 27 | 27 | 4.24E+07 |
| Q0VCU1 | Cytoplasmic aconitate hydratase | ACO1 | 412.13 | 36 | 36 | 4.03E+07 |
| Q03247 | Apolipoprotein E | APOE | 297.9 | 11 | 11 | 3.87E+07 |
| P28800 | Alpha-2-antiplasmin | SERPINF2 | 308.14 | 12 | 12 | 3.70E+07 |
| Q3ZCJ2 | Aldo-keto reductase family 1 member A1 | AKR1A1 | 375.77 | 23 | 22 | 3.63E+07 |
| Q3T0P6 | Phosphoglycerate kinase 1 | PGK1 | 364.33 | 19 | 19 | 3.61E+07 |

| | | | | | | |
|---|---|---|---|---|---|---|
| Q9N2I2 | Plasma serine protease inhibitor | SERPINA5 | 330.85 | 13 | 13 | 3.25E+07 |
| Q3SX14 | Gelsolin | GSN | 374.98 | 23 | 23 | 2.91E+07 |
| P00978 | Protein AMBP | AMBP | 312.06 | 11 | 11 | 2.83E+07 |
| Q32LP0 | Fermitin family homolog 3 | FERMT3 | 393.59 | 22 | 22 | 2.75E+07 |
| Q2KJF1 | Alpha-1B-glycoprotein | A1BG | 374.41 | 17 | 17 | 2.67E+07 |
| P02584 | Profilin-1 | PFN1 | 246.79 | 7 | 7 | 2.46E+07 |
| P62935 | Peptidyl-prolyl cis-trans isomerase A | PPIA | 273.82 | 10 | 10 | 2.42E+07 |
| P52898 | Dihydrodiol dehydrogenase 3 | N/A | 362.45 | 16 | 16 | 2.40E+07 |
| P01267 | Thyroglobulin | TG | 477.61 | 61 | 61 | 2.40E+07 |
| P68103 | Elongation factor 1-alpha 1 | EEF1A1 | 303.67 | 11 | 11 | 2.32E+07 |
| O46415 | Ferritin light chain | FTL | 250.13 | 6 | 6 | 2.28E+07 |
| A6H768 | Galactokinase | GALK1 | 356.39 | 15 | 15 | 2.23E+07 |
| Q2KIG3 | Carboxypeptidase B2 | CPB2 | 317.74 | 15 | 15 | 2.21E+07 |
| Q92176 | Coronin-1A | CORO1A | 304.36 | 16 | 16 | 2.17E+07 |
| Q3SZR3 | Alpha-1-acid glycoprotein | ORM1 | 283.21 | 8 | 8 | 2.10E+07 |
| Q2KJH9 | 4-trimethylaminobutyraldehyde dehydrogenase | ALDH9A1 | 277.31 | 13 | 13 | 2.04E+07 |
| Q2KJH4 | WD repeat-containing protein 1 | WDR1 | 339.39 | 17 | 17 | 1.94E+07 |
| Q27975 | Heat shock 70 kDa protein 1A | HSPA1A | 370.73 | 19 | 11 | 1.91E+07 |
| Q27965 | Heat shock 70 kDa protein 1B | HSPA1B | 370.73 | 19 | 11 | 1.91E+07 |
| P52556 | Flavin reductase (NADPH) | BLVRB | 325.49 | 9 | 9 | 1.80E+07 |
| Q0VCK0 | Bifunctional purine biosynthesis protein ATIC | ATIC | 322.53 | 20 | 20 | 1.78E+07 |
| P14568 | Argininosuccinate synthase | ASS1 | 289.04 | 12 | 12 | 1.77E+07 |
| Q3SZV7 | Hemopexin | HPX | 328.93 | 12 | 12 | 1.75E+07 |
| P19120 | Heat shock cognate 71 kDa protein | HSPA8 | 380.74 | 21 | 14 | 1.73E+07 |
| A3KMV5 | Ubiquitin-like modifier-activating enzyme 1 | UBA1 | 389.49 | 27 | 27 | 1.65E+07 |
| Q3SYV4 | Adenylyl cyclase-associated protein 1 | CAP1 | 333.75 | 14 | 14 | 1.54E+07 |
| P81948 | Tubulin alpha-4A chain | TUBA4A | 325.66 | 15 | 5 | 1.46E+07 |
| Q3T054 | GTP-binding nuclear protein Ran | RAN | 195.51 | 8 | 8 | 1.45E+07 |
| Q3SYU2 | Elongation factor 2 | EEF2 | 315.15 | 23 | 23 | 1.44E+07 |
| Q3MHN2 | Complement component C9 | C9 | 295.57 | 15 | 15 | 1.39E+07 |
| P61223 | Ras-related protein Rap-1b | RAP1B | 238.08 | 11 | 11 | 1.30E+07 |
| P07589 | Fibronectin | FN1 | 417.14 | 38 | 38 | 1.28E+07 |
| P28801 | Glutathione S-transferase P | GSTP1 | 245.61 | 6 | 6 | 1.24E+07 |
| P02672 | Fibrinogen alpha chain | FGA | 300.49 | 15 | 15 | 1.22E+07 |
| P37141 | Glutathione peroxidase 3 | GPX3 | 229.26 | 7 | 7 | 1.21E+07 |
| O18879 | Glutathione S-transferase A2 | GSTA2 | 148.7 | 4 | 3 | 1.20E+07 |
| Q2KJD0 | Tubulin beta-5 chain | TUBB5 | 360.2 | 20 | 4 | 1.06E+07 |
| P12799 | Fibrinogen gamma-B chain | FGG | 264.17 | 11 | 11 | 1.03E+07 |
| Q76LV1 | Heat shock protein HSP 90-beta | HSP90AB1 | 348.54 | 23 | 11 | 1.01E+07 |
| Q29RQ1 | Complement component C7 | C7 | 323.89 | 18 | 18 | 9.92E+06 |
| Q9TTJ5 | Regucalcin | RGN | 241.4 | 9 | 9 | 9.82E+06 |
| P12378 | UDP-glucose 6-dehydrogenase | UGDH | 290.35 | 16 | 16 | 9.40E+06 |
| Q76LV2 | Heat shock protein HSP 90- | HSP90AA1 | 353.2 | 26 | 15 | 8.99E+06 |

| | | | | | | |
|---|---|---|---|---|---|---|
| | alpha | | | | | |
| Q95M17 | Acidic mammalian chitinase | CHIA | 254.93 | 10 | 10 | 8.95E+06 |
| P19217 | Sulfotransferase 1E1 | SULT1E1 | 206.1 | 6 | 6 | 8.81E+06 |
| Q5EA79 | Galactose mutarotase | GALM | 265.7 | 7 | 7 | 8.78E+06 |
| P17690 | Beta-2-glycoprotein 1 | APOH | 265.16 | 9 | 9 | 8.36E+06 |
| P00741 | Coagulation factor IX | F9 | 242.92 | 10 | 10 | 7.92E+06 |
| P19879 | Mimecan | OGN | 214.97 | 8 | 8 | 7.54E+06 |
| Q5EAD2 | D-3-phosphoglycerate dehydrogenase | PHGDH | 293.94 | 12 | 12 | 7.49E+06 |
| P81644 | Apolipoprotein A-II | APOA2 | 202.89 | 6 | 6 | 7.02E+06 |
| A7E3W2 | Galectin-3-binding protein | LGALS3BP | 240.19 | 10 | 10 | 6.86E+06 |
| P01044 | Kininogen-1 | KNG1 | 331.08 | 21 | 7 | 6.83E+06 |
| P16116 | Aldo-keto reductase family 1 member B1 | AKR1B1 | 229.55 | 10 | 9 | 6.54E+06 |
| Q08DP0 | Phosphoglucomutase-1 | PGM1 | 288.05 | 12 | 12 | 6.42E+06 |
| E1BF81 | Corticosteroid-binding globulin | SERPINA6 | 233.44 | 7 | 7 | 6.19E+06 |
| Q3ZC42 | Alcohol dehydrogenase class-3 | ADH5 | 274.43 | 11 | 11 | 5.89E+06 |
| Q5E9F7 | Cofilin-1 | CFL1 | 247.13 | 8 | 7 | 5.45E+06 |
| P02070 | Hemoglobin subunit beta | HBB | 228.08 | 8 | 4 | 5.43E+06 |
| Q5E9B7 | Chloride intracellular channel protein 1 | CLIC1 | 223.31 | 9 | 8 | 5.41E+06 |
| Q0VCM5 | Inter-alpha-trypsin inhibitor heavy chain H1 | ITIH1 | 290.35 | 12 | 12 | 5.40E+06 |
| P13605 | Fibromodulin | FMOD | 216.76 | 7 | 7 | 5.36E+06 |
| Q28035 | Glutathione S-transferase A1 | GSTA1 | 162.5 | 5 | 4 | 5.35E+06 |
| Q5EA01 | Beta-1 4-glucuronyltransferase 1 | B4GAT1 | 216.19 | 5 | 5 | 5.29E+06 |
| Q3SZD7 | Carbonyl reductase [NADPH] 1 | CBR1 | 254.05 | 10 | 10 | 5.07E+06 |
| Q3ZBD7 | Glucose-6-phosphate isomerase | GPI | 262.55 | 11 | 11 | 4.96E+06 |
| P50227 | Sulfotransferase 1A1 | SULT1A1 | 237.14 | 9 | 9 | 4.95E+06 |
| P38657 | Protein disulfide-isomerase A3 | PDIA3 | 266.74 | 13 | 13 | 4.79E+06 |
| Q28106 | Contactin-1 | CNTN1 | 327.43 | 14 | 14 | 4.66E+06 |
| A7MBI7 | Catechol O-methyltransferase | COMT | 247.32 | 7 | 7 | 4.61E+06 |
| P02676 | Fibrinogen beta chain | FGB | 313.24 | 16 | 16 | 4.43E+06 |
| P12260 | Coagulation factor XIII A chain (Fragment) | F13A1 | 185 | 4 | 4 | 4.24E+06 |
| P22226 | Cathelicidin-1 | CATHL1 | 214.94 | 6 | 6 | 4.20E+06 |
| P63048 | Ubiquitin-60S ribosomal protein L40 | UBA52 | 164.96 | 4 | 4 | 4.16E+06 |
| P0CG53 | Polyubiquitin-B | UBB | 164.96 | 4 | 4 | 4.16E+06 |
| P0CH28 | Polyubiquitin-C | UBC | 164.96 | 4 | 4 | 4.16E+06 |
| P62992 | Ubiquitin-40S ribosomal protein S27a | RPS27A | 164.96 | 4 | 4 | 4.16E+06 |
| Q2KJC6 | S-adenosylmethionine synthase isoform type-1 | MAT1A | 245.62 | 6 | 6 | 4.08E+06 |
| G3MYZ3 | Afamin | AFM | 275.48 | 14 | 14 | 3.95E+06 |
| Q32KY0 | Apolipoprotein D | APOD | 126.93 | 3 | 3 | 3.94E+06 |
| Q5E956 | Triosephosphate isomerase | TPI1 | 286.03 | 10 | 10 | 3.92E+06 |
| Q3MHN5 | Vitamin D-binding protein | GC | 353.07 | 22 | 3 | 3.90E+06 |
| Q5E9Z2 | Hyaluronan-binding protein 2 | HABP2 | 209.38 | 9 | 9 | 3.74E+06 |
| Q5E9B1 | L-lactate dehydrogenase B chain | LDHB | 267.6 | 9 | 7 | 3.71E+06 |

| | | | | | | |
|---|---|---|---|---|---|---|
| P50448 | Factor XIIa inhibitor | N/A | 212.68 | 5 | 5 | 3.67E+06 |
| Q2TA49 | Vasodilator-stimulated phosphoprotein | VASP | 224.75 | 8 | 8 | 3.56E+06 |
| Q9N0V4 | Glutathione S-transferase Mu 1 | GSTM1 | 185 | 5 | 5 | 3.54E+06 |
| P30932 | CD9 antigen | CD9 | 148.75 | 2 | 2 | 3.53E+06 |
| Q0VCM4 | Glycogen phosphorylase liver form | PYGL | 263.84 | 14 | 12 | 3.51E+06 |
| O77742 | Osteomodulin | OMD | 97.42 | 2 | 2 | 3.50E+06 |
| P00745 | Vitamin K-dependent protein C (Fragment) | PROC | 256.8 | 8 | 8 | 3.48E+06 |
| Q56JW4 | Adenine phosphoribosyltransferase | APRT | 241.4 | 7 | 7 | 3.41E+06 |
| Q3SZB7 | Fructose-1 6-bisphosphatase 1 | FBP1 | 266.6 | 9 | 9 | 3.39E+06 |
| Q9TTE1 | Serpin A3-1 | SERPINA3-1 | 245.71 | 9 | 3 | 3.27E+06 |
| Q1JP75 | L-xylulose reductase | DCXR | 172.39 | 4 | 4 | 3.27E+06 |
| Q2KJ33 | LIM and senescent cell antigen-like-containing domain protein 2 | LIMS2 | 109.34 | 2 | 2 | 3.23E+06 |
| O77783 | Exostosin-2 | EXT2 | 294.87 | 14 | 14 | 3.18E+06 |
| Q9TT36 | Thyroxine-binding globulin | SERPINA7 | 199.23 | 5 | 5 | 3.18E+06 |
| Q3SWY2 | Integrin-linked protein kinase | ILK | 202.91 | 11 | 11 | 3.09E+06 |
| P00435 | Glutathione peroxidase 1 | GPX1 | 193.02 | 6 | 6 | 3.01E+06 |
| A6QLR4 | Flotillin-2 | FLOT2 | 230.63 | 12 | 12 | 2.96E+06 |
| O02659 | Mannose-binding protein C | MBL | 227.8 | 5 | 5 | 2.49E+06 |
| Q0VCX2 | Endoplasmic reticulum chaperone BiP | HSPA5 | 292.02 | 13 | 11 | 2.43E+06 |
| Q9XSC6 | Creatine kinase M-type | CKM | 243.47 | 6 | 5 | 2.42E+06 |
| Q28065 | C4b-binding protein alpha chain | C4BPA | 189.17 | 5 | 5 | 2.41E+06 |
| Q5E9F5 | Transgelin-2 | TAGLN2 | 198.57 | 8 | 8 | 2.41E+06 |
| P33433 | Histidine-rich glycoprotein (Fragment) | HRG | 190.53 | 4 | 4 | 2.39E+06 |
| P08169 | Cation-independent mannose-6-phosphate receptor | IGF2R | 317.72 | 24 | 24 | 2.34E+06 |
| Q0VCX1 | Complement C1s subcomponent | C1S | 220.07 | 7 | 7 | 2.30E+06 |
| Q3MHM5 | Tubulin beta-4B chain | TUBB4B | 355.23 | 19 | 1 | 2.30E+06 |
| Q32PI5 | Serine/threonine-protein phosphatase 2A 65 kDa regulatory subunit A alpha isoform | PPP2R1A | 245.4 | 9 | 9 | 2.30E+06 |
| Q0P5F9 | 2-aminomuconic semialdehyde dehydrogenase | ALDH8A1 | 298.98 | 10 | 10 | 2.28E+06 |
| Q2TBQ3 | Guanidinoacetate N-methyltransferase | GAMT | 188.77 | 5 | 5 | 2.25E+06 |
| Q3SZV3 | Elongation factor 1-gamma | EEF1G | 227.66 | 7 | 7 | 2.22E+06 |
| Q2KJH6 | Serpin H1 | SERPINH1 | 231.96 | 7 | 7 | 2.21E+06 |
| P11151 | Lipoprotein lipase | LPL | 204.76 | 5 | 5 | 2.13E+06 |
| P48644 | Aldehyde dehydrogenase 1A1 | ALDH1A1 | 224.88 | 10 | 10 | 2.08E+06 |
| Q3MHP2 | Ras-related protein Rab-11B | RAB11B | 108.69 | 3 | 3 | 2.08E+06 |
| Q2TA29 | Ras-related protein Rab-11A | RAB11A | 108.69 | 3 | 3 | 2.08E+06 |
| Q2KJ93 | Cell division control protein 42 homolog | CDC42 | 166.73 | 5 | 4 | 2.05E+06 |
| O46414 | Ferritin heavy chain | FTH1 | 192.07 | 5 | 5 | 2.01E+06 |
| Q3ZC84 | Cytosolic non-specific | CNDP2 | 220.02 | 8 | 8 | 1.96E+06 |

| | | | | | | |
|---|---|---|---|---|---|---|
| | dipeptidase | | | | | |
| P50397 | Rab GDP dissociation inhibitor beta | GDI2 | 289.99 | 11 | 8 | 1.95E+06 |
| A3KN12 | Adenylosuccinate lyase | ADSL | 151.31 | 3 | 3 | 1.94E+06 |
| Q2TBX4 | Heat shock 70 kDa protein 13 | HSPA13 | 205.89 | 7 | 7 | 1.92E+06 |
| Q32PF2 | ATP-citrate synthase | ACLY | 207.64 | 10 | 10 | 1.91E+06 |
| Q95M18 | Endoplasmin | HSP90B1 | 241.11 | 12 | 11 | 1.90E+06 |
| A1A4J1 | ATP-dependent 6-phosphofructokinase liver type | PFKL | 291.72 | 12 | 12 | 1.89E+06 |
| Q08DA1 | Sodium/potassium-transporting ATPase subunit alpha-1 | ATP1A1 | 298.17 | 13 | 13 | 1.88E+06 |
| P12820 | Angiotensin-converting enzyme | ACE | 224.6 | 9 | 9 | 1.86E+06 |
| A5D7I4 | Exostosin-1 | EXT1 | 186.39 | 5 | 5 | 1.85E+06 |
| Q3T014 | Bisphosphoglycerate mutase | BPGM | 160.09 | 3 | 3 | 1.84E+06 |
| Q95122 | Monocyte differentiation antigen CD14 | CD14 | 199.28 | 5 | 5 | 1.82E+06 |
| Q1JPJ2 | Xaa-Pro aminopeptidase 1 | XPNPEP1 | 268.26 | 10 | 10 | 1.81E+06 |
| Q3T149 | Heat shock protein beta-1 | HSPB1 | 216.98 | 8 | 8 | 1.80E+06 |
| P61585 | Transforming protein RhoA | RHOA | 219.13 | 7 | 7 | 1.80E+06 |
| P52193 | Calreticulin | CALR | 233.14 | 7 | 7 | 1.74E+06 |
| Q27967 | Secreted phosphoprotein 24 | SPP2 | 204.59 | 7 | 7 | 1.74E+06 |
| O77834 | Peroxiredoxin-6 | PRDX6 | 235.92 | 9 | 9 | 1.73E+06 |
| Q3T0Z7 | Dihydropteridine reductase | QDPR | 201.7 | 6 | 6 | 1.66E+06 |
| P35445 | Cartilage oligomeric matrix protein | COMP | 256.57 | 14 | 12 | 1.55E+06 |
| Q58D31 | Sorbitol dehydrogenase | SORD | 235.29 | 7 | 7 | 1.52E+06 |
| Q5E9P9 | Serine hydroxymethyltransferase cytosolic | SHMT1 | 225.74 | 8 | 8 | 1.51E+06 |
| Q06805 | Tyrosine-protein kinase receptor Tie-1 | TIE1 | 260.54 | 10 | 10 | 1.37E+06 |
| Q2KJ32 | Methanethiol oxidase | SELENBP1 | 217.03 | 7 | 7 | 1.34E+06 |
| Q17QH6 | Collectin-11 | COLEC11 | 169.01 | 4 | 4 | 1.33E+06 |
| Q95JC7 | Neutral amino acid transporter B(0) | SLC1A5 | 185.48 | 6 | 6 | 1.30E+06 |
| Q5E9I6 | ADP-ribosylation factor 3 | ARF3 | 206.93 | 6 | 3 | 1.29E+06 |
| P84080 | ADP-ribosylation factor 1 | ARF1 | 206.93 | 6 | 3 | 1.29E+06 |
| Q32PJ2 | Apolipoprotein A-IV | APOA4 | 176.75 | 6 | 6 | 1.29E+06 |
| A6QR56 | Aldehyde dehydrogenase family 16 member A1 | ALDH16A1 | 207.51 | 8 | 8 | 1.28E+06 |
| Q29RU4 | Complement component C6 | C6 | 257.29 | 12 | 12 | 1.26E+06 |
| Q3SZP2 | Microtubule-associated protein RP/EB family member 2 | MAPRE2 | 168.98 | 5 | 5 | 1.25E+06 |
| Q71SP7 | Fatty acid synthase | FASN | 259.3 | 14 | 14 | 1.24E+06 |
| P54149 | Mitochondrial peptide methionine sulfoxide reductase | MSRA | 210.05 | 6 | 6 | 1.24E+06 |
| P13696 | Phosphatidylethanolamine-binding protein 1 | PEBP1 | 176.35 | 3 | 3 | 1.22E+06 |
| Q2TBI0 | Lipopolysaccharide-binding protein | LBP | 224.47 | 8 | 8 | 1.21E+06 |
| Q3B7M9 | Glycogen phosphorylase brain form | PYGB | 261.43 | 11 | 7 | 1.16E+06 |

| | | | | | | |
|---|---|---|---|---|---|---|
| Q148C9 | Heme-binding protein 1 | HEBP1 | 170.15 | 3 | 3 | 1.10E+06 |
| O02675 | Dihydropyrimidinase-related protein 2 | DPYSL2 | 255.17 | 9 | 9 | 1.07E+06 |
| Q9GMB8 | Serine--tRNA ligase cytoplasmic | SARS1 | 190.64 | 7 | 7 | 1.07E+06 |
| Q32L99 | Prostaglandin reductase 2 | PTGR2 | 131.43 | 3 | 3 | 1.06E+06 |
| Q9XTA2 | Prolyl endopeptidase | PREP | 155.74 | 4 | 4 | 1.02E+06 |
| P48034 | Aldehyde oxidase 1 | AOX1 | 214.8 | 10 | 10 | 1.01E+06 |
| P19858 | L-lactate dehydrogenase A chain | LDHA | 220.69 | 8 | 6 | 9.98E+05 |
| Q3SYZ6 | Xylulose kinase | XYLB | 222.89 | 8 | 8 | 9.97E+05 |
| Q2KIT0 | Protein HP-20 homolog | N/A | 130.2 | 2 | 2 | 9.92E+05 |
| Q08E11 | Peptidyl-prolyl cis-trans isomerase C | PPIC | 122.4 | 3 | 3 | 9.77E+05 |
| P53712 | Integrin beta-1 | ITGB1 | 156.09 | 3 | 3 | 9.61E+05 |
| Q28156 | cGMP-specific 3' 5'-cyclic phosphodiesterase | PDE5A | 225.39 | 8 | 8 | 9.36E+05 |
| P01131 | Low-density lipoprotein receptor | LDLR | 199.95 | 10 | 10 | 8.87E+05 |
| Q28178 | Thrombospondin-1 | THBS1 | 178.65 | 6 | 6 | 8.86E+05 |
| P00514 | cAMP-dependent protein kinase type I-alpha regulatory subunit | PRKAR1A | 186.13 | 5 | 5 | 8.85E+05 |
| Q9BGI3 | Peroxiredoxin-2 | PRDX2 | 120.68 | 2 | 2 | 8.77E+05 |
| Q3SZJ4 | Prostaglandin reductase 1 | PTGR1 | 189.16 | 5 | 5 | 8.61E+05 |
| Q2KIA5 | Dynamin-1-like protein | DNM1L | 210.36 | 7 | 7 | 8.42E+05 |
| Q2HJ33 | Obg-like ATPase 1 | OLA1 | 156.53 | 6 | 6 | 8.16E+05 |
| P61286 | Polyadenylate-binding protein 1 | PABPC1 | 180.38 | 6 | 6 | 8.08E+05 |
| P00376 | Dihydrofolate reductase | DHFR | 169.36 | 3 | 3 | 8.07E+05 |
| P80109 | Phosphatidylinositol-glycan-specific phospholipase D | GPLD1 | 171.48 | 5 | 5 | 7.95E+05 |
| Q2KJ63 | Plasma kallikrein | KLKB1 | 179.64 | 7 | 7 | 7.89E+05 |
| Q2T9Y6 | Glutamate--cysteine ligase regulatory subunit | GCLM | 126.55 | 2 | 2 | 7.68E+05 |
| Q59A32 | Trifunctional purine biosynthetic protein adenosine-3 | GART | 252.83 | 7 | 7 | 7.65E+05 |
| Q3ZCD0 | CD81 antigen | CD81 | 138.39 | 2 | 2 | 7.57E+05 |
| Q8SQA4 | Adhesion G protein-coupled receptor E5 | ADGRE5 | 187.62 | 6 | 6 | 7.32E+05 |
| Q3T0F5 | Ras-related protein Rab-7a | RAB7A | 144.35 | 4 | 4 | 7.30E+05 |
| P01045 | Kininogen-2 | KNG2 | 281.57 | 17 | 3 | 7.24E+05 |
| P00515 | cAMP-dependent protein kinase type II-alpha regulatory subunit | PRKAR2A | 190.44 | 5 | 5 | 7.15E+05 |
| Q32PA4 | 14 kDa phosphohistidine phosphatase | PHPT1 | 121.39 | 2 | 2 | 7.11E+05 |
| P19035 | Apolipoprotein C-III | APOC3 | 125.35 | 2 | 2 | 6.92E+05 |
| Q3ZBN5 | Asporin | ASPN | 150.04 | 4 | 4 | 6.29E+05 |
| P21809 | Biglycan | BGN | 139.24 | 3 | 3 | 6.22E+05 |
| Q1JPA6 | Arylamine N-acetyltransferase 1 | NAT1 | 169.21 | 7 | 7 | 6.10E+05 |
| P61157 | Actin-related protein 3 | ACTR3 | 199.22 | 4 | 4 | 5.93E+05 |
| Q1RMU3 | Prolyl 4-hydroxylase subunit alpha-1 | P4HA1 | 247.17 | 10 | 10 | 5.73E+05 |
| Q2KIS7 | Tetranectin | CLEC3B | 147.96 | 3 | 3 | 5.73E+05 |

| | | | | | | |
|---|---|---|---|---|---|---|
| Q5EA61 | Creatine kinase B-type | CKB | 202.97 | 5 | 4 | 5.73E+05 |
| Q58CQ9 | Pantetheinase | VNN1 | 178.47 | 4 | 4 | 5.65E+05 |
| P00743 | Coagulation factor X | F10 | 167.57 | 6 | 6 | 5.60E+05 |
| A7YWP4 | Histidine ammonia-lyase | HAL | 146.72 | 5 | 5 | 5.59E+05 |
| Q2TBQ8 | 6-phosphogluconolactonase | PGLS | 207.14 | 5 | 5 | 5.53E+05 |
| P60712 | Actin cytoplasmic 1 | ACTB | 348.4 | 16 | 1 | 5.43E+05 |
| Q3SZI4 | 14-3-3 protein theta | YWHAQ | 162.01 | 3 | 2 | 5.39E+05 |
| P41541 | General vesicular transport factor p115 | USO1 | 202.12 | 7 | 7 | 5.27E+05 |
| Q2YDE4 | Proteasome subunit alpha type-6 | PSMA6 | 117.62 | 3 | 3 | 5.16E+05 |
| P25975 | Procathepsin L | CTSL | 148.16 | 3 | 3 | 5.15E+05 |
| Q5E998 | Cathepsin L2 | CTSV | 148.16 | 3 | 3 | 5.15E+05 |
| P13384 | Insulin-like growth factor-binding protein 2 | IGFBP2 | 157.9 | 4 | 4 | 5.15E+05 |
| Q28085 | Complement factor H | CFH | 194.32 | 7 | 7 | 4.92E+05 |
| Q0VD19 | Sphingomyelin phosphodiesterase | SMPD1 | 104.74 | 2 | 2 | 4.77E+05 |
| Q5NTB3 | Coagulation factor XI | F11 | 152.35 | 6 | 6 | 4.72E+05 |
| Q148J6 | Actin-related protein 2/3 complex subunit 4 | ARPC4 | 83.02 | 2 | 2 | 4.71E+05 |
| Q3T0W4 | Protein phosphatase 1 regulatory subunit 7 | PPP1R7 | 128.23 | 4 | 4 | 4.57E+05 |
| P63103 | 14-3-3 protein zeta/delta | YWHAZ | 175.53 | 5 | 4 | 4.53E+05 |
| Q3SZ65 | Eukaryotic initiation factor 4A-II | EIF4A2 | 210.95 | 6 | 1 | 4.52E+05 |
| Q3SWW8 | Thrombospondin-4 | THBS4 | 231.94 | 7 | 5 | 4.50E+05 |
| P62261 | 14-3-3 protein epsilon | YWHAE | 164.5 | 5 | 4 | 4.15E+05 |
| P18902 | Retinol-binding protein 4 | RBP4 | 104.87 | 2 | 2 | 4.10E+05 |
| Q3ZBG0 | Proteasome subunit alpha type-7 | PSMA7 | 148.12 | 2 | 2 | 3.96E+05 |
| Q9TU25 | Ras-related C3 botulinum toxin substrate 2 | RAC2 | 145.64 | 5 | 2 | 3.96E+05 |
| Q32KL2 | Proteasome subunit beta type-5 | PSMB5 | 114.95 | 2 | 2 | 3.95E+05 |
| P24627 | Lactotransferrin | LTF | 213.56 | 10 | 6 | 3.90E+05 |
| P68250 | 14-3-3 protein beta/alpha | YWHAB | 153.64 | 4 | 3 | 3.89E+05 |
| Q5I597 | Betaine--homocysteine S-methyltransferase 1 | BHMT | 155.74 | 5 | 5 | 3.72E+05 |
| A2VDS1 | Selenocysteine lyase | SCLY | 151.94 | 4 | 4 | 3.70E+05 |
| Q0V8B6 | Tripeptidyl-peptidase 1 | TPP1 | 135.75 | 4 | 4 | 3.68E+05 |
| Q3MHR0 | Acyl-protein thioesterase 1 | LYPLA1 | 82.5 | 2 | 2 | 3.62E+05 |
| Q9XSG3 | Isocitrate dehydrogenase [NADP] cytoplasmic | IDH1 | 137.63 | 3 | 3 | 3.61E+05 |
| Q2TBU0 | Haptoglobin | HP | 130.26 | 2 | 2 | 3.47E+05 |
| P49907 | Selenoprotein P | SELENOP | 157.59 | 3 | 3 | 3.41E+05 |
| Q3SZJ0 | Argininosuccinate lyase | ASL | 160.29 | 4 | 4 | 3.32E+05 |
| Q5E9A3 | Poly(rC)-binding protein 1 | PCBP1 | 93.99 | 2 | 2 | 3.31E+05 |
| Q3SZH7 | Leukotriene A-4 hydrolase | LTA4H | 144.71 | 3 | 3 | 3.22E+05 |
| P17248 | Tryptophan--tRNA ligase cytoplasmic | WARS1 | 143.21 | 4 | 4 | 3.15E+05 |
| Q58DK4 | Triokinase/FMN cyclase | TKFC | 185.72 | 3 | 3 | 3.05E+05 |
| Q3ZBA8 | Protein NDRG2 | NDRG2 | 91.91 | 2 | 2 | 3.05E+05 |
| Q0VCG9 | Pentraxin-related protein PTX3 | PTX3 | 89.62 | 2 | 2 | 2.96E+05 |

| | | | | | | |
|---|---|---|---|---|---|---|
| P31754 | Uridine 5'-monophosphate synthase | UMPS | 158.19 | 5 | 5 | 2.85E+05 |
| P28782 | Protein S100-A8 | S100A8 | 81.13 | 2 | 2 | 2.82E+05 |
| P11064 | Low molecular weight phosphotyrosine protein phosphatase | ACP1 | 112.51 | 2 | 2 | 2.81E+05 |
| P56701 | 26S proteasome non-ATPase regulatory subunit 2 | PSMD2 | 176.97 | 5 | 5 | 2.69E+05 |
| Q2HJF4 | Phenazine biosynthesis-like domain-containing protein | PBLD | 110.11 | 2 | 2 | 2.65E+05 |
| A4FUA8 | F-actin-capping protein subunit alpha-1 | CAPZA1 | 155.23 | 4 | 3 | 2.64E+05 |
| Q3ZBH0 | T-complex protein 1 subunit beta | CCT2 | 154.4 | 4 | 4 | 2.62E+05 |
| P26452 | 40S ribosomal protein SA | RPSA | 162.78 | 4 | 4 | 2.56E+05 |
| Q3ZC09 | Beta-enolase | ENO3 | 278.89 | 8 | 2 | 2.46E+05 |
| Q2YDJ9 | Mannose-1-phosphate guanyltransferase beta | GMPPB | 126 | 3 | 3 | 2.44E+05 |
| P21856 | Rab GDP dissociation inhibitor alpha | GDI1 | 230.48 | 6 | 3 | 2.44E+05 |
| Q28007 | Dihydropyrimidine dehydrogenase [NADP(+)] | DPYD | 157.17 | 6 | 6 | 2.43E+05 |
| Q2TBN5 | COMM domain-containing protein 9 | COMMD9 | 94.17 | 2 | 2 | 2.42E+05 |
| Q3T004 | Serum amyloid P-component | APCS | 122.33 | 3 | 3 | 2.40E+05 |
| Q2TBW7 | Sorting nexin-2 | SNX2 | 134.42 | 5 | 4 | 2.27E+05 |
| P55906 | Transforming growth factor-beta-induced protein ig-h3 | TGFBI | 180.92 | 5 | 5 | 2.20E+05 |
| Q2KI42 | 26S proteasome non-ATPase regulatory subunit 11 | PSMD11 | 108.4 | 2 | 2 | 2.16E+05 |
| Q3T0S5 | Fructose-bisphosphate aldolase B | ALDOB | 137.96 | 3 | 3 | 2.10E+05 |
| Q29RU2 | Oncoprotein-induced transcript 3 protein | OIT3 | 162.59 | 4 | 4 | 2.10E+05 |
| A5PK51 | Nicotinate phosphoribosyltransferase | NAPRT | 195.78 | 6 | 6 | 2.01E+05 |
| P63258 | Actin cytoplasmic 2 | ACTG1 | 348.71 | 16 | 1 | 1.94E+05 |
| Q3ZCJ8 | Dipeptidyl peptidase 1 | CTSC | 122.77 | 3 | 3 | 1.89E+05 |
| Q9TRY0 | Peptidyl-prolyl cis-trans isomerase FKBP4 | FKBP4 | 146.44 | 5 | 5 | 1.82E+05 |
| P15246 | Protein-L-isoaspartate(D-aspartate) O-methyltransferase | PCMT1 | 111.5 | 2 | 2 | 1.80E+05 |
| Q32LM2 | Small glutamine-rich tetratricopeptide repeat-containing protein alpha | SGTA | 150.18 | 3 | 3 | 1.77E+05 |
| P0CB32 | Heat shock 70 kDa protein 1-like | HSPA1L | 302.15 | 11 | 1 | 1.75E+05 |
| Q8SPF8 | BPI fold-containing family B member 1 | BPIFB1 | 125.77 | 2 | 2 | 1.68E+05 |
| P02465 | Collagen alpha-2(I) chain | COL1A2 | 145.68 | 2 | 2 | 1.67E+05 |
| Q5E9F9 | 26S proteasome regulatory subunit 7 | PSMC2 | 127.44 | 4 | 4 | 1.58E+05 |
| P00432 | Catalase | CAT | 141.57 | 4 | 4 | 1.54E+05 |
| A7MBJ5 | Cullin-associated NEDD8-dissociated protein 1 | CAND1 | 138.71 | 3 | 3 | 1.49E+05 |

| | | | | | | |
|---|---|---|---|---|---|---|
| Q6URK6 | Cadherin-5 | CDH5 | 136.06 | 5 | 5 | 1.45E+05 |
| Q3ZBM5 | Sorting nexin-5 | SNX5 | 106.89 | 2 | 2 | 1.44E+05 |
| P81125 | Alpha-soluble NSF attachment protein | NAPA | 115.26 | 3 | 3 | 1.33E+05 |
| P68509 | 14-3-3 protein eta | YWHAH | 137.76 | 3 | 2 | 1.30E+05 |
| Q5E973 | 60S ribosomal protein L18 | RPL18 | 97.85 | 2 | 2 | 1.23E+05 |
| Q3SZF2 | ADP-ribosylation factor 4 | ARF4 | 181.16 | 4 | 1 | 1.14E+05 |
| Q28017 | Platelet-activating factor acetylhydrolase | PLA2G7 | 91.62 | 2 | 2 | 1.10E+05 |
| Q8SPP7 | Peptidoglycan recognition protein 1 | PGLYRP1 | 153.58 | 2 | 2 | 1.08E+05 |
| Q3SZK8 | Na(+)/H(+) exchange regulatory cofactor NHE-RF1 | SLC9A3R1 | 149.64 | 3 | 3 | 1.04E+05 |
| Q3ZCI9 | T-complex protein 1 subunit theta | CCT8 | 142.73 | 2 | 2 | 1.02E+05 |
| Q2YDG0 | G-protein coupled receptor family C group 5 member C | GPRC5C | 106.34 | 2 | 2 | 9.39E+04 |
| Q2KJD2 | Vesicle-associated membrane protein 3 | VAMP3 | 116.95 | 3 | 3 | 7.94E+04 |
| F1N152 | Serine protease HTRA1 | HTRA1 | 139.78 | 3 | 3 | 7.87E+04 |
| Q3ZBT5 | Syntaxin-7 | STX7 | 123.03 | 3 | 3 | 7.68E+04 |
| Q2TBH7 | Ras-related protein Rab-4A | RAB4A | 122.47 | 3 | 2 | 6.77E+04 |
| Q17R06 | Ras-related protein Rab-21 | RAB21 | 83.08 | 2 | 2 | 4.30E+04 |
| Q95114 | Lactadherin | MFGE8 | 85.69 | 2 | 2 | 3.53E+04 |

N/A: non annotated gene.

**Table S4.** GO terms referring to cellular components that were found to be over-represented in medium proteins using the BiNGO plugin for Cytoscape software platform (*p*-values < 0.001).

| GO terms | Protein count | corr p-value | -log (corr P value) |
|---|---|---|---|
| extracellular region | 54 | 4.28E-27 | 26.37 |
| extracellular space | 43 | 3.38E-24 | 23.47 |
| cytosol | 36 | 1.52E-07 | 6.82 |
| spherical high-density lipoprotein particle | 3 | 6.51E-05 | 4.19 |
| pigment granule | 6 | 6.51E-05 | 4.19 |
| melanosome | 6 | 6.51E-05 | 4.19 |
| high-density lipoprotein particle | 4 | 1.71E-04 | 3.77 |
| chylomicron | 3 | 3.31E-04 | 3.48 |
| fibrinogen complex | 3 | 3.31E-04 | 3.48 |
| haptoglobin-hemoglobin complex | 3 | 3.41E-04 | 3.47 |
| plasma lipoprotein particle | 4 | 3.41E-04 | 3.47 |
| lipoprotein particle | 4 | 3.41E-04 | 3.47 |
| protein-lipid complex | 4 | 3.94E-04 | 3.40 |
| hemoglobin complex | 3 | 3.94E-04 | 3.40 |
| very-low-density lipoprotein particle | 3 | 7.71E-04 | 3.11 |
| triglyceride-rich plasma lipoprotein particle | 3 | 7.71E-04 | 3.11 |

**Table S5.** GO terms referring to cellular components that were found to be under-represented in medium proteins using the BiNGO plugin for Cytoscape software platform (*p*-values < 0.05).

| GO terms | Protein count | corr p-value | -log (corr P value) |
|---|---|---|---|
| membrane | 25 | 2.72E-03 | 2.57 |
| organelle | 38 | 1.50E-02 | 1.82 |
| intracellular organelle | 38 | 3.75E-02 | 1.43 |
